# Evolutionary flexibility in the maternal-to-zygotic transition reveals alternative routes to embryogenesis across eukaryotes

**DOI:** 10.64898/2026.04.21.719843

**Authors:** Kenny A. Bogaert, Pélagie Ratchinski, Min Zheng, Fabian B. Haas, Rita A. Batista, Susana M. Coelho

## Abstract

The maternal-to-zygotic transition (MZT), during which developmental control shifts from maternal products to the zygotic genome, is a fundamental feature of embryogenesis. The prevailing model, established from animal systems, holds that maternal provisioning and nuclear-to-cytoplasmic (N/C) ratio determine the timing of zygotic genome activation (ZGA). Whether this developmental logic is conserved across multicellular eukaryotes remains unknown. Using brown algae, an independently evolved multicellular lineage spanning broad natural variation in maternal provisioning, we generated comparative embryonic transcriptomes across three species and resolved parental contributions through reciprocal crosses and allele-specific expression. Contrary to canonical animal models, all species activated their zygotic genomes immediately after fertilisation, including highly provisioned species with low N/C ratios, and showed no evidence of stable parental imprinting. These findings reveal that timing and regulation of MZT during embryogenesis is evolutionarily flexible and independently shaped across the tree of life.

## INTRODUCTION

The maternal-to-zygotic transition (MZT) represents a pivotal early developmental milestone in which control of gene expression shifts from maternally supplied transcripts and proteins to the zygotic genome. The MZT comprises two coupled processes: clearance of maternal mRNAs and proteins, and zygotic genome activation (ZGA), the onset of transcription from the zygote’s own genome. This transition involves extensive molecular reprogramming, including the degradation of maternal mRNAs and proteins, and chromatin remodeling to establish new transcriptional and regulatory landscapes that enable totipotency and subsequent cell differentiation ^1,2^.

The timing and regulation of the MZT vary widely across animals and plants ^3^. In animals, the nuclear-to-cytoplasmic (N/C) ratio has been proposed as a universal timer of zygotic genome activation (ZGA), with embryos reaching a critical N/C threshold to initiate transcription; accordingly, embryos with higher N/C ratios or slower division rates activate the zygotic genome earlier, whereas rapidly dividing embryos delay activation ^3,4^. Beyond timing, the question of whether maternal and paternal genomes contribute equally to ZGA has emerged as of considerable interest ^5^. Plants also exhibit substantially great diversity in the timing and parental regulation of the MZT ^6^. For instance, in flowering plants the extent of maternal regulation of early embryo development has been debated ^7–11^ but zygotic transcription begins relatively early in *Arabidopsis* ^12^, maize ^13^ and rice ^14,15^. In *Arabidopsis* a transient maternal dominance of transcripts, persisting into the globular stage due to asymmetric genome activation has been reported ^16^ whereas in liverworts like *Marchantia*, development of the sporophyte, which is retained on the female gametophyte, relies almost exclusively on maternal transcripts ^6^. Collectively, these studies show that although the MZT is a universal developmental process, current understanding of the mechanisms regulating its timing derives overwhelmingly from animals, with more limited insights from plants, while the regulation of this transition remains largely unexplored across other eukaryotic lineages. Expanding the study of the MZT beyond animals and plants is therefore essential for understanding the evolutionary origins and diversity of early developmental reprogramming across eukaryotes.

Brown algae (Phaeophyceae), members of the Stramenopiles, represent an independent origin of complex multicellularity with genomic and regulatory architectures distinct from those of animals and plants ^17–20^. As such, they provide a powerful system for examining the evolutionary diversity of MZT. Notably, brown algae display extensive variation in reproductive strategies, including gamete dimorphism ranging from near-isogamy (almost equal-sized gametes) to oogamy (large eggs and small sperm), and differing levels of maternal care, from freely developing zygotes to those retained on maternal tissues ^21–24^. These traits are expected to shape the degree of maternal provisioning and thus the duration of maternal regulatory control during early development, and provide an opportunity to test the universality of N/C ratio as a regulator of developmental transitions. Based on observations from animal systems, we would therefore expect that increased maternal provisioning and maternal retention would prolong maternal control through greater inheritance of maternal transcripts and maternally biased transcription, whereas limited provisioning would shorten this maternal phase.

Here, by analysing the timing of zygotic transcription and maternal transcript clearance across brown algal species spanning a reproductive spectrum, we characterise the MZT in a third independent lineage of multicellular eukaryotes and reveal how reproductive and life-history traits shape the evolution of this fundamental developmental transition. We show that, contrary to expectations derived from animals, maternal provisioning does not delay zygotic genome activation. Instead, increased maternal investment is associated with immediate and extensive activation of the zygotic genome in brown algae. We further find no evidence for stable parental imprinting, indicating that parent-of-origin effects during early embryogenesis are transient rather than maintained developmental states in this lineage. Together, these findings demonstrate that the timing and regulation of the MZT are evolutionarily flexible and shaped by reproductive strategy rather than the degree of maternal investment alone.

## RESULTS

### Transcriptomic profiling across early development in algae with contrasting gamete dimorphism

We selected three brown algal species representing different levels of gamete size dimorphism ^21,23^ (**Figure 1, A and B**). *Dictyota dichotoma* displays an oogamous reproductive system, with tiny, motile male gametes and large, non-flagellated female gametes ^25^. After spawning and fertilisation, the zygote develops independently of the maternal tissue (**Figure 1C**). *Undaria pinnatifida*, a kelp, is near-oogamous, with small male gametes and larger flagellated female gametes that remain attached to the maternal gametophyte during early zygotic development (**Figure 1D**) with the maternal attachment providing a developmental cue to the sporophyte ^26^. Finally, *Scytosiphon promiscuus* is nearly isogamous: male gametes are only slightly smaller than female gametes, and both are motile, with zygote development proceeding independently of the maternal tissue (**Figure 1E**). This gamete size gradient across species results in substantial variation in the N/C ratio of the zygotes, providing a unique opportunity to test hypotheses about how N/C ratio and maternal provisioning influence MZT timing. Interestingly, we note that brown algae occupy a wide range of N/C ratios relative to animals and land plants (**Figure 1F, Table S1**). Transmission electron microscopy reveals marked differences in nuclear ultrastructure between sexes and species (**Figure 1G**). In the oogamous species, female gamete nuclei show expanded euchromatin and a prominent nucleolus, consistent with a transcriptionally permissive state, whereas in the near-isogamous *Scytosiphon* the female nucleus is more compact and electron-dense, very similar to the male gamete largely heterochromatic nuclei. Therefore, nuclear scaling and organisation vary systematically with gamete dimorphism in brown algae, providing a structural basis for interpreting early developmental transcriptional states.

**Figure 1.**
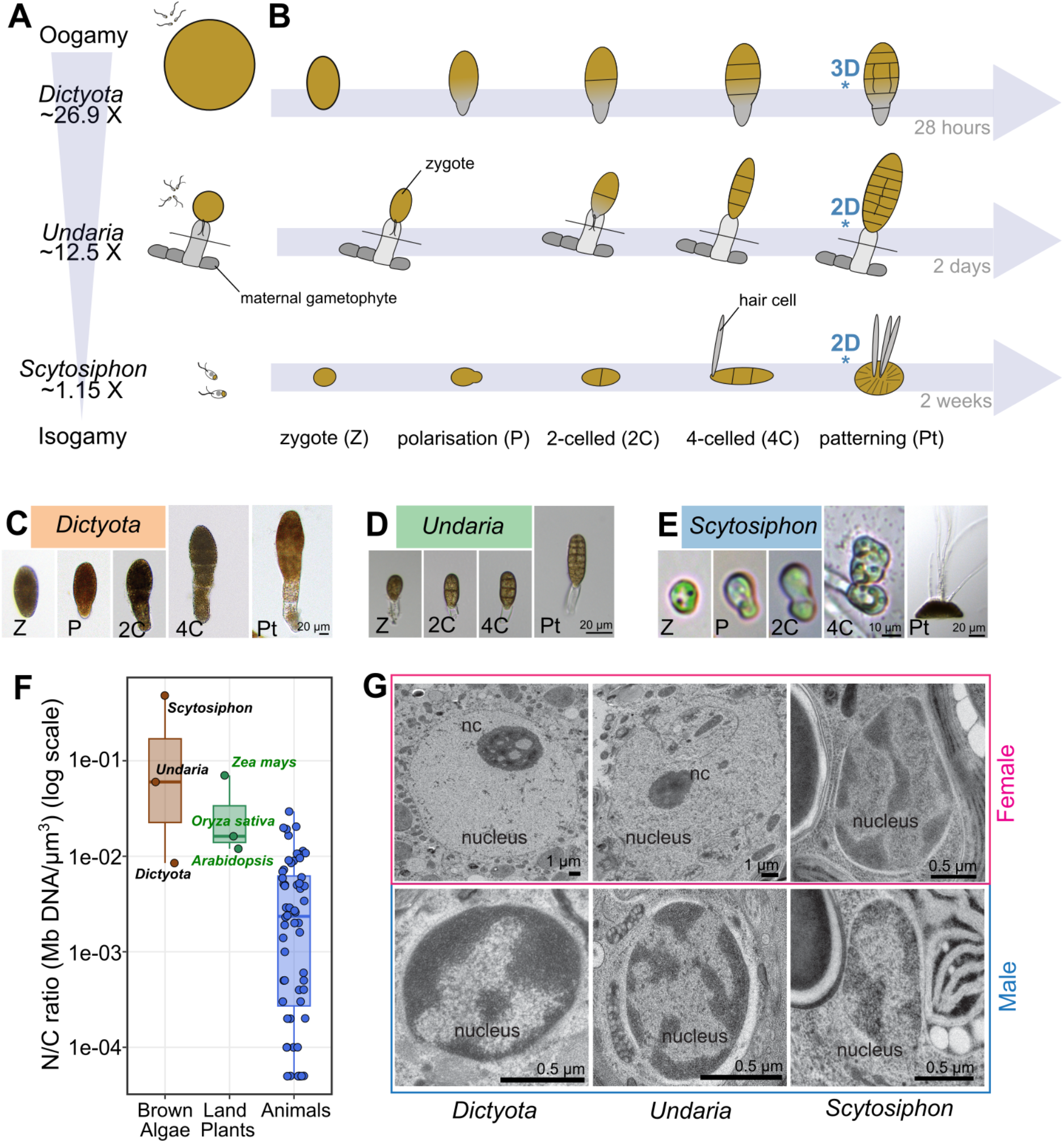
Transcriptomic profiling of early developmental stages across brown algae with distinct gamete dimorphism. (**A**) Schematic overview of fertilisation and early developmental progression in the studied brown algae (*Dictyota, Undaria* and *Scytosiphon*), spanning oogamy to isogamy. Relative gamete size dimorphism (ratio female:male diameter) is indicated and depicted with gamete sizes proportional to the relative diameter ratios. (**B**) Representative stages from zygote formation through early patterning together with approximate developmental timescales. Transitions to 2D or 3D growth (*) are indicated in blue. (**C, D, E**) Light micrographs of early developmental stages: zygote (Z), polarised zygote (P), 2-cell (2C), 4-cell (4C), and patterned stage (Pt). Scale bars as indicated. (**F**) N/C ratio (Mb DNA per µm3 cytoplasm; log scale) comparing animals, plants, and brown algae (see Table S1). (**G**) Transmission electron micrographs of female and male gamete nuclei showing the nuclear ultrastructure. nc = nucleolus.

To investigate MZT dynamics, we manually microdissected material from distinct developmental stages in three brown algal species and generated stage-specific transcriptomes by RNA-seq (**Figure 1, A and B**; **Tables S2, S3**). Samples included eggs, sperm, zygotes, germinating zygotes, two-cell embryos, four-cell embryos, and patterning embryos (see Methods). In *Dictyota*, apical-basal polarity is established between the zygote and the polarisation stage (5.5 h after fertilisation, AF)^27^, and three-dimensional (3D) growth, with cell differentiation initiated at the patterning stage. In *Undaria*, the zygote is maternally prepolarised ^28^ and the embryo transitions to two-dimensional (2D) growth at the patterning stage. Note that due to species-specific developmental dynamics, the polarisation stage could not be sampled in *Undaria*. Finally, *Scytosiphon*’s polarity is established at the polarisation stage without cell differentiation, and 2D growth emerges at the patterning stage. Developmental tempo also varies substantially across these three species, ranging from 28 hours to the patterning stage in *Dictyota*, to two days in *Undaria* and two weeks in *Scytosiphon*.

For each stage, at least three independent biological replicates were prepared and sequenced to a depth exceeding ∼18 million read pairs per library (**Table S3**). Transcriptomes from the same stage were highly reproducible across replicates (**Figure S1**). This dataset enabled us to resolve the timing and architecture of ZGA and parental transcript clearance across species with divergent reproductive strategies, providing new insight into the evolutionary conservation and diversification of early developmental reprogramming across the spectrum of maternal investment in brown algae.

### Heterochronic maternal-to-zygotic transitions across brown algae

Principal component analysis of the sampled transcriptomes revealed distinct patterns of transcriptome remodeling across species (**Figure 2A**). In *Dictyota* and *Undaria*, transcriptomic changes were progressive and homogeneous across developmental stages, suggesting continuous and coordinated remodeling following fertilisation. By contrast, in *Scytosiphon*, early developmental samples clustered together before a secondary group emerged later in development. This observation reflects a biphasic pattern of early development in which an initial period of transcriptomic change is followed by a major remodeling event at the four-cell stage, coinciding with the onset of cell differentiation (see **Figure 1**). These distinct dynamics raise the question of whether they reflect differences in zygotic genome activation and maternal transcript clearance, and if so, what drives them. To address this, we examined the timing of maternal transcript clearance and *de novo* zygotic transcription directly (**Figure 2, B to D**).

**Figure 2.**
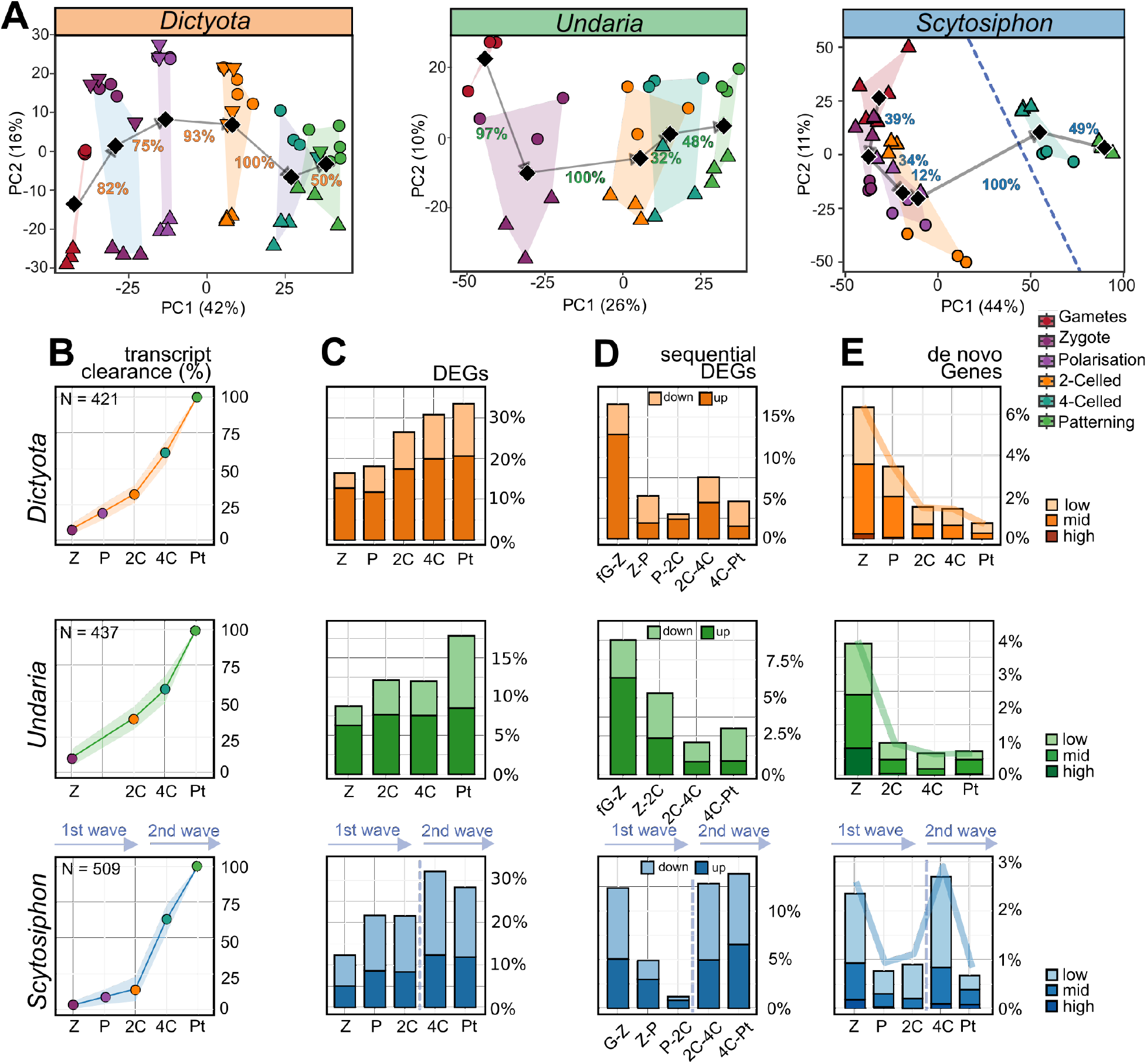
Early zygotic genome activation in brown algae. (**A**) Principal component analysis of the transcriptome from gametes to multicellular embryos. Circles and triangles: reciprocal crosses; inverted triangles: internal crosses (see also **Figure 4a**). Diamonds denote group centroids. Percentages indicate the Euclidean distance between consecutive stage centroids in PCA space, normalized to the largest inter-stage distance (=100%) following ^29^. (**B-E**) Each row corresponds to one species: *Dictyota dichotoma* (top), *Undaria pinnatifida* (middle), *Scytosiphon promiscuus* (bottom). (**B**) Progressive clearance of transcripts across development. Curves show, for each developmental stage, the cumulative percentage of transcripts that are monotonically downregulated and have decreased to below 10% of their expression level in female gametes (see Methods for definition and calculation). “Monotonically” indicates that transcript expression decreases (or remains unchanged) across successive developmental stages, without any increase at later stages. N indicates the total number of monotonically cleared genes (**C**) Number of differentially expressed genes (DEGs) relative to female gametes (fGam) (*Dictyota, Undaria*) and gametes (Gam) (*Scytosiphon*) at each developmental stage, coloured by up or down regulation (dark and light-coloured, respectively). (**D**) Sequential stage-to-stage DEGs, indicating the number of genes becoming differentially expressed at each transition, coloured by expression level (low, mid, high). (**E**) Emergence of de novo expressed genes (absent in gametes, but present at each respective developmental stage) grouped by expression level based on mean DESeq2-normalized counts: low (5-10), mid (10-100), and high (>100). In *Scytosiphon*, dashed lines denote our inferred two distinct transcriptional waves during early embryogenesis. Developmental stages: Z, zygote; P, polarisation; 2C, 2-celled; 4C, 4-celled; Pt, patterning. Note that *Undaria* lacks the polarisation stage.

To characterise the clearance of gamete-derived transcripts, we focused on transcripts that were monotonically downregulated across development (*i. e*, transcripts present in gametes and whose expression decreases (or remains unchanged) across successive developmental stages; see Methods). Consistent with transcriptome trajectories (**Figure 2D**), transcript clearance occurred progressively during embryogenesis but differed markedly in timing among species (**Figure 2b**). In *Dictyota* and *Undaria*, gamete derived transcripts declined rapidly, with steep reductions during polarisation and first cell division stages. In contrast, *Scytosiphon* showed delayed parental transcript clearance, indicating prolonged parental control during early development (2-4 cell stage) preceded by a period of relatively low transcriptomic change.

We next quantified changes in transcript abundance relative to the gametes (**Figure 2C**). For oogamous species (*Dictyota* and *Undaria*), female gametes were used as the reference, whereas in the near-isogamous *Scytosiphon*, the combined male and female gamete transcriptome served as the baseline. In all species, the number of differentially expressed genes (DEGs) (compared to gametes) increased as development progressed, but the timing of this accumulation differed. Importantly, many of these DEGs were upregulated relative to the gamete, consistent with active, *de novo* transcription (see also **Figure S2**). In *Dictyota* and *Undaria*, most DEGs emerged during polarisation and the first division, whereas *Scytosiphon* showed an additional pronounced increase substantially later, at the 4-cell stage. This pattern is consistent with an early, major transcriptional wave in *Dictyota* and *Undaria*, whereas in *Scytosiphon* a relatively modest initial wave is followed by a larger wave of *de novo* transcription at later stages (**Figure 2**).

Transcriptional changes across sequential developmental stages further supported these contrasting dynamics between species (**Figure 2D**). In *Dictyota* and *Undaria*, DEGs were evenly distributed across sequential developmental transitions, with the strongest changes occurring around fertilisation and first cell division, coinciding with the first asymmetric cell division (see **Figure 1A**). By contrast, *Scytosiphon* exhibited a more abrupt transition later in development. Similarly, the first appearance of newly expressed genes (i.e., those absent from the gamete transcriptome) revealed early activation of the zygotic genome in *Dictyota* and *Undaria* whereas the largest burst of newly expressed genes in *Scytosiphon* occurred at the 4-cell stage, when cells differentiate (**Figure 2E**). All together, these analyses indicate a single and early transcriptional activation in *Dictyota* and *Undaria*, but time-modulated, biphasic transcriptional activation in *Scytosiphon*.

### Conserved developmental programs despite extensive transcriptional rewiring

To assess whether conserved developmental dynamics are driven by shared gene regulatory programs, we performed weighted gene co-expression network analysis (WGCNA; signed network; **Figure 3A**, see Methods) separately for each species. The overall expression profile (as represented by the module eigengene values) of large modules (**Figure S2, D to F, Figure 3B**) was significantly correlated with developmental time, and several modules displayed significant overlap in orthologous genes across species (**Figure 3A**, bold modules). Notably, these conserved modules were associated with the MZT, showing either increasing expression as development progressed (‘MZT-induced modules’) or decreasing expression (likely corresponding to cleared maternal transcripts, ‘MZT repressed modules’). Moreover, MZT-associated modules were enriched in evolutionarily old genes, as determined by phylostratigraphy ^30–32^ (see Material and Methods, **Table S4-S5**), both at the level of entire modules and among hub genes. In particular, the top 25% of genes with the highest module membership were significantly enriched in old gene classes (**Figure 3A**). This suggests that deeply conserved gene families occupy central positions within developmental gene regulatory networks. Conversely, unassigned genes (grey module) were enriched in young genes, suggesting that evolutionary novelty is associated with limited co-expression network integration.

**Figure 3.**
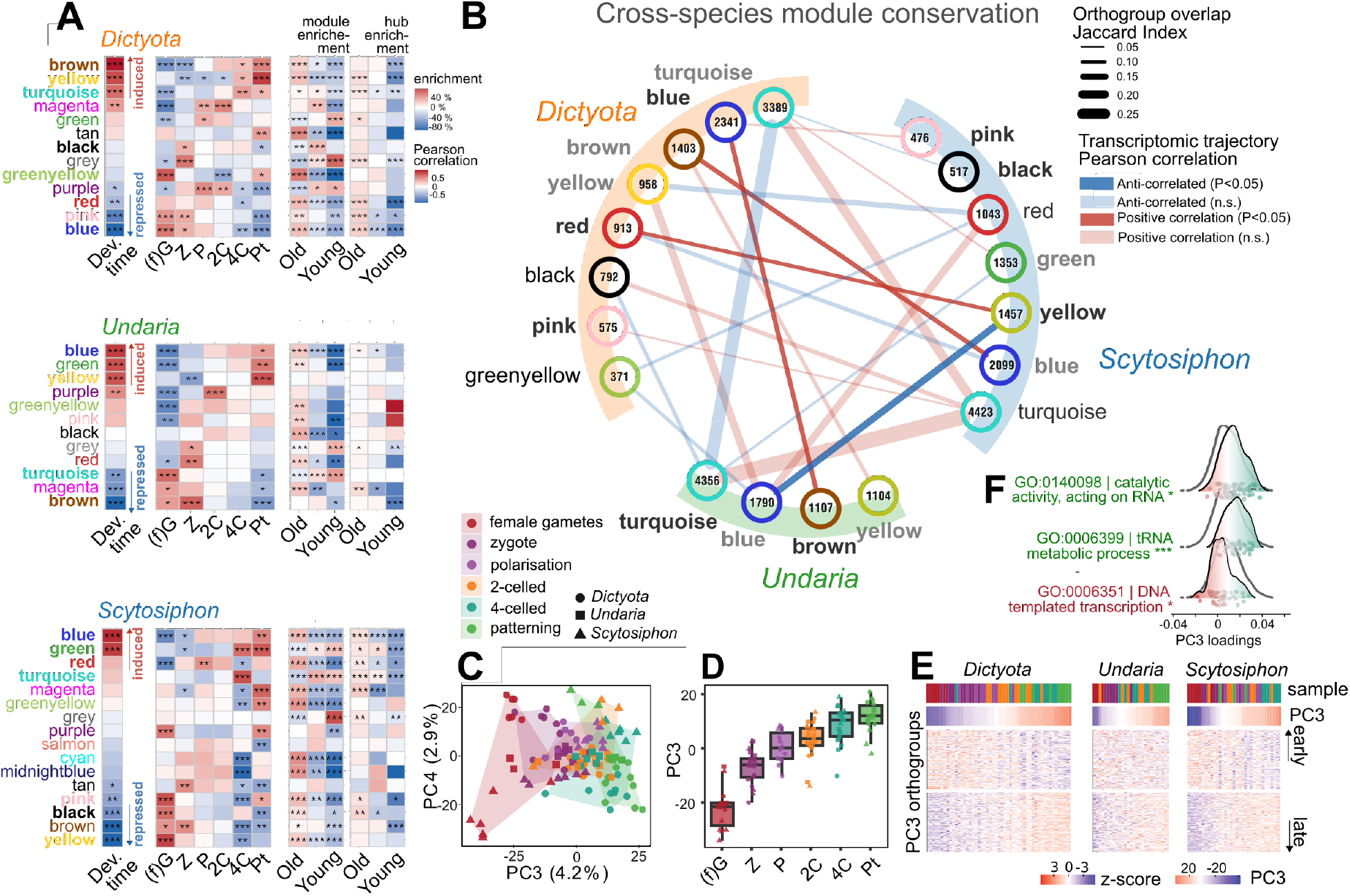
A conserved developmental transcriptional program despite gene-level divergence in brown algal embryogenesis. **(A)** Pearson correlations between WGCNA module eigengenes and developmental stages. *Dev. time*: correlation with stage as ordinal rank (1 = female gametes → 6 = patterning); red = progressively induced (late/zygotic), blue = progressively repressed. Per-stage columns capture stage-specific peaks. Asterisks: significance; bold names: cross-species conservation (see B). Adjacent panels: Fisher’s exact tests for enrichment of evolutionarily young, intermediate, or old genes across each module (left) and its top 25% hub genes (right). **(B)** Cross-species module conservation network summarising shared gene membership and temporal dynamics. Nodes: WGCNA modules (number = size in OGs). Edges: significant OG overlap (BH-adjusted p < 0.05, odds ratio ≥ 1.5, >50 shared OGs). Bold black/grey modules: negatively/positively correlated with developmental time. **(C)** PCA of orthogroup expression (TPM). PC3 separates samples by species (**Figure S3D**) from early to late stages regardless of species, capturing a conserved developmental axis. **(D)** PC3 sample loadings by stage. **(E)** Heatmaps of z-scored expression for the top 100 negative and positive PC3-loading OGs per species, illustrating a conserved early-to-late transcriptional shift. **(F)** GO enrichment of PC3-associated OGs relative to background (white).

We next quantified module conservation across species by combining two measures: overlap in orthogroups composition (Jaccard index) and similarity of expression trajectories (based on module eigengene values) (**Figure 3b, Table S6**). Several module pairs displayed both significant gene overlap and correlated temporal dynamics, indicating that parts of the developmental transcriptome are organized into conserved regulatory units despite substantial evolutionary divergence between the studied species. This conservation was most pronounced for MZT-induced modules and MZT-repressed modules. However, in some cases, modules with overlapping gene content showed anti-correlated eigengene trajectories, suggesting partial rewiring of transcriptional programs across species.

Single copy orthogroups (N=8126) (**Figure S3A, Data S5**) where compiled to enable cross-species comparison of transcriptomic profiles. PCA analysis of orthogroup expression revealed a conserved developmental progression across species and recapitulated the biphasic MZT transition in *Scytosiphon* **(Figure S3B**). When analysed jointly, the primary axes of variation (PC1 and PC2) were dominated by species-specific differences in transcript abundance (**Figure S3D**). However, PC3 separated samples by developmental stage rather than species identity (**Figure 3, C and D**), indicating that developmental progression represents a consistent source of variation across species. In line with this, mean-centering expression per orthogroup reduced species-specific offsets and enhanced the separation of developmental stages along the leading principal components (**Figure S4**). When samples were ordered by their PC3 scores, these orthogroups displayed a coordinated gradient of expression from high to low, with samples arranged according to developmental time (**Figure 3E**), consistent with PC3 capturing a common transcriptional program underlying developmental progression. Together, these results support a conserved developmental trajectory across brown algal embryogenesis, despite divergence in absolute expression levels and network organisation.

Functional enrichment analysis of orthogroups associated with the conserved developmental axis (PC3) revealed distinct biological processes at early and late stages of development. Orthogroups with negative PC3 loadings, which correspond to early developmental stages (gametes and zygotes), were enriched for transcriptional regulation (**Figure 3F, Data S6**). In contrast, orthogroups with positive PC3 loadings, corresponding to later stages, were enriched for RNA metabolism and protein synthesis. Therefore, early development is characterised by the activation of regulatory networks associated with the onset of zygotic transcription, whereas later stages are marked by increased RNA processing and translational capacity, supporting both the expansion of the zygotic transcriptome and the turnover of maternally derived transcripts.

Across all metrics, *Dictyota* and *Undaria* were consistently more similar to each other than to *Scytosiphon* (**Figure S5**). Gene expression profiles become increasingly similar between species over the course of development (**Figure S5A**). After correcting for baseline differences in expression levels, this similarity is most pronounced at mid-developmental stages, indicating that shared temporal expression patterns (rather than absolute expression levels) drive this convergence. In contrast, similarity at early stages is primarily due to a common set of highly expressed genes, likely reflecting conserved maternal contributions (**Figure S5, B to E**).

Species sharing similar levels of gamete size dimorphism might be expected to exhibit more similar co-expression networks. However, we did not observe increased network similarity between the two oogamous species (*Dictyota* and *Undaria*). This observation suggests that their phenotypic similarity may not be driven by conserved co-expression modules, but could instead reflect convergent evolution of transcriptional programs associated with oogamy. This interpretation is consistent with previous phylogenetic analyses indicating independent origins of oogamy in Laminariales and Dictyotales ^31,33,34^.

Altogether, these analyses suggest that early embryogenesis in brown algae is shaped by a conserved, MZT-associated developmental program that is robust to substantial divergence in gene expression and network architecture. While co-expression modules show only partial conservation and evidence of transcriptional rewiring, particularly between lineages, a shared developmental trajectory emerges at the systems level. This is reflected in partially conserved eigengene dynamics and a common transcriptional axis across species. This conserved program is supported by evolutionarily old genes that occupy central positions within co-expression networks, indicating deep evolutionary roots of early developmental regulation, particularly in transcriptional control and activation. At the same time, lineage-specific differences in module composition and temporal dynamics highlight flexibility in how these conserved processes are implemented, consistent with the convergent evolution of reproductive strategies such as oogamy. Therefore, similar developmental outcomes can arise from divergent regulatory architectures, underscoring the evolutionary plasticity of multicellular development.

### Parental contributions to the early embryonic transcriptome

To examine parental contributions to the early embryonic transcriptome, we performed reciprocal crosses between two genetically distinct strains for each of the three species (**Figure 4A**). Between 5700 and 16141 genes containing at least one informative SNP were identified, enabling assignment of maternal and paternal reads in the transcriptomes (**Table S8, S9, Data S1 to S3**). For each developmental stage and species, the maternal fraction of expression (maternal reads divided by total of reads assigned to a parent) was calculated for each gene in both reciprocal crosses (**Figure 4, B to D**). In parallel, allele-specific expression was tested for each gene using edgeR’s exact test on maternal and paternal read counts normalised by library size (see methods).

**Figure 4.**
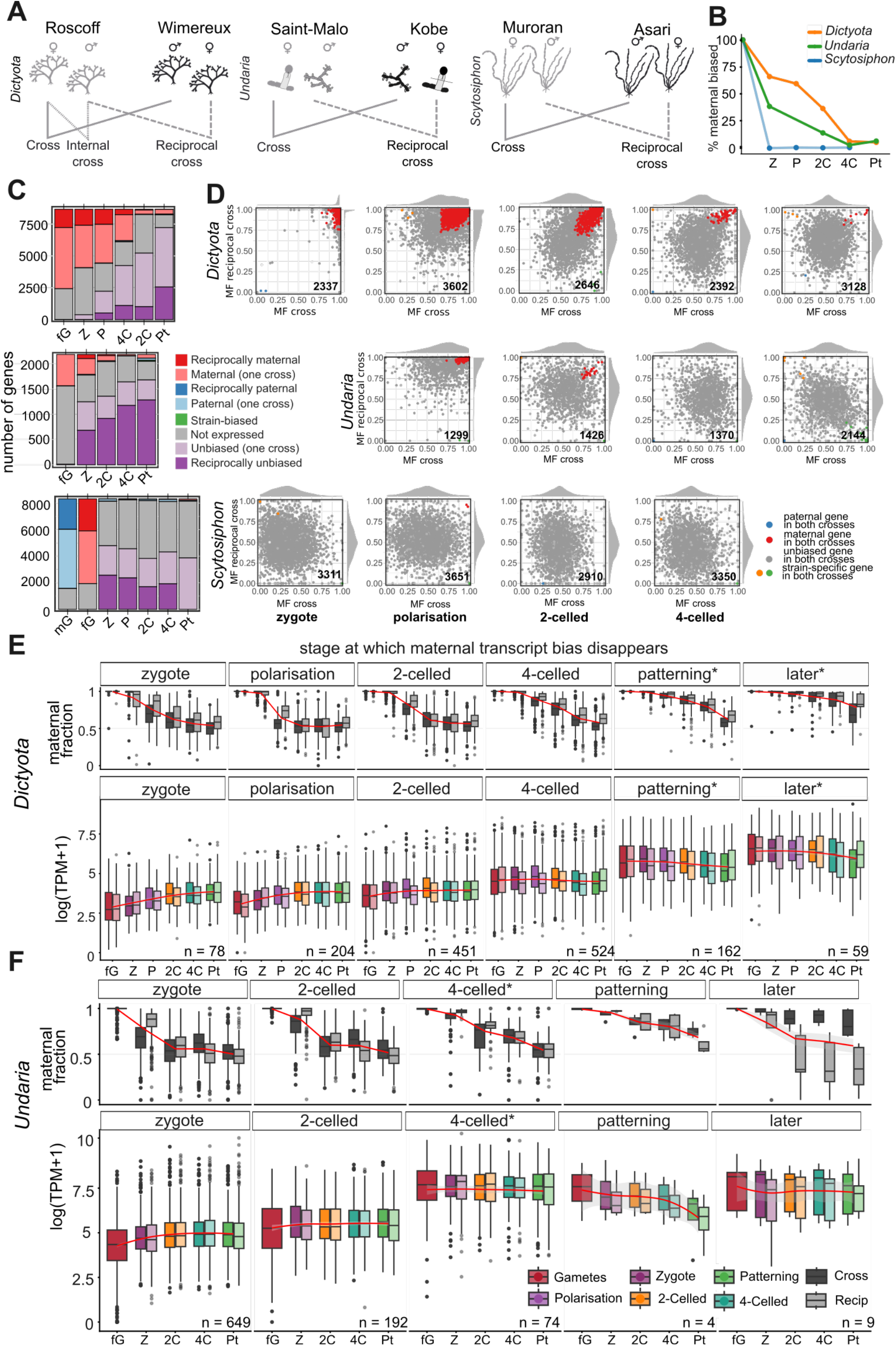
Parental genome bias in transcript abundance during early development in brown algae. **(A)** Schematic representation of the reciprocal crosses performed for each species. **(B)** Fraction of reciprocally maternally biased genes across developmental stages relative to the female gametes **(C)** Number of genes in each bias category across developmental stages, for *Scytosiphon, Undaria* and *Dictyota*. Only genes for which the presence or absence of bias could be statistically assessed in at least one developmental stage are shown. For *Scytosiphon*, the patterning stage and for *Undaria* the egg cell stage are based on a single cross due to data availability. Colors: dark red, reciprocally maternal; light red, maternally biased in one cross; dark blue, reciprocally paternal; light blue, paternally biased in one cross; green, strain-biased; purple, reciprocally unbiased; light purple, unbiased in one cross; grey tones, undetermined, not expressed, or other. **(D)** Mean maternal fraction in the cross versus reciprocal cross for individual genes in *Scytosiphon, Undaria* and *Dictyota*, across developmental stages (n genes indicated per panel). Points are coloured by bias category: red, maternally biased in both crosses; blue, paternally biased in both crosses; yellow/green, strain-specific bias. (**E**,**F**) Maternal fraction (top) and log(TPM + 1) (bottom) across developmental stages for gene groups defined by the stage at which they first show unbiased transcript abundance, in *Dictyota* **(E)** and *Undaria* **(F)**. The number of genes per group is indicated. Smoothed trends (loess) are shown in red. Statistical differences between groups in overall expression level and expression trajectory were assessed by linear mixed modelling; see Results section. Developmental stages marked with an asterisk showed statistically significant enrichment of GO terms, summarized in **Figure S10**.

Genes showing significant maternal bias in both crosses were classified as maternal (red, **Figure 4D**), those with paternal bias in both crosses as paternal (blue), and genes consistently biased toward one strain, irrespective of parental origin, were classified separately (yellow and green; full gene lists in **Data S1to 3**).

In *Dictyota*, the early embryonic transcriptome was strongly dominated by maternal transcripts (**Figure 4, B and C**). Consistent with this observation, the earliest developmental samples clustered closer to egg cells than to later embryonic stages in principal component analyses on genetic variants (**Figure S6**). As development proceeded, the maternal fraction gradually decreased and the majority of genes approached unbiased expression (**Figure 4, B and C**). Interestingly, this transition toward balanced parental expression occurred earlier in *Undaria* than in *Dictyota* (**Figure 4, B and C**). This difference may reflect the slower developmental pace of *Undaria* or differences in egg size and maternal provisioning. The earlier loss of maternal bias in *Undaria* relative to developmental stage may therefore partly reflect this difference in developmental tempo, providing support for a correlation with developmental speed ^3^. However, tempo alone cannot fully account for the patterns observed, since both oogamous species still share rapid ZGA onset relative to the partially delayed, biphasic activation in *Scytosiphon*, which develops the most slowly of the three. Importantly, the same temporal trend was observed in the inter-strain and in an intra-strain cross in *Dictyota* indicating that the observed dynamics are unlikely to result from differences in genetic divergence between parental strains (**Figure S9**).

In contrast, the nearly isogamous species *Scytosiphon* showed evidence of both parental alleles contributing to the transcriptome already at the zygote stage (**Figure 4, B and C**). This pattern could arise from early zygotic transcription but may simply reflect the near-equal cytoplasmic contributions of the two gametes. To distinguish between these possibilities, we compared the observed maternal fraction at the zygote stage with the fraction expected under a simple dilution model assuming that the male gamete contributes 80% of the volume of the female gamete^23^ (**Figure S9**). Under this model, genes should fall along the diagonal if dilution alone explains the observed maternal fractions. Instead, substantial dispersion from the expected relationship was observed, with Pearson correlations of 0.28 and 0.24 for the cross and reciprocal cross, respectively (r^2^ = 0.079 and 0.058). Thus, dilution alone explains only ∼6–8% of the observed variance in maternal fraction at the zygote stage, suggesting that additional processes such as transcriptional activation or differential transcript stability contribute to the observed patterns, consistent with the differential expression observed between zygotes and gametes (**Figure 2C**).

We next quantified the number of genes displaying a bias in the abundance of maternal or paternal transcripts at each stage (**Figure 4, B and C)**. In *Dictyota* and *Undaria*, the number of maternally biased genes decreased progressively during development as unbiased expression became increasingly prevalent. In *Scytosiphon*, by contrast, the majority of genes showed balanced parental transcript abundance from the earliest stages, consistent with the allele-specific patterns observed in the maternal fraction analysis.

To further characterise the dynamics of maternal transcript inheritance, genes in the two oogamous species were grouped according to the developmental stage at which they lost maternal bias in both reciprocal crosses (**Figure 4D**). Genes retaining maternal bias until late developmental stages generally displayed the highest transcript abundance throughout development, whereas genes that lost maternal bias early showed lower average expression levels and often increased expression over time. To formally test these observations, we fitted a linear mixed model with log-transformed expression as a function of developmental stage, gene group, their interaction, and cross direction as a covariate, with gene identity as a random intercept. In both species, genes losing maternal bias later showed progressively higher average expression levels across all developmental stages (linear trend across ordered groups — *Dictyota*: estimate = 20.97, z = 24.2, p < 0.0001; *Undaria*: estimate = 6.24, z = 6.7, p < 0.0001). Similarly, the interaction between developmental stage and gene group was significant in both species, with a monotonic shift from positive to negative expression slopes across groups (linear × linear contrast — *Dictyota*: estimate = -2.22, z = -26.7, p < 0.0001; *Undaria*: estimate = -0.72, z = -6.4, p < 0.0001). Thus, the timing of maternal bias loss is associated with distinct developmental expression trajectories: late-transitioning genes generally decline in expression across development, whereas early-transitioning genes tend to become more highly expressed over time. These patterns suggest that long-lasting maternal bias primarily reflects the persistence of large transcript pools deposited in the egg cell, which are gradually diluted or replaced as embryonic transcription increases.

Functional enrichment analysis of these gene groups revealed that genes with sustained maternal bias and high transcript abundance were enriched in housekeeping functions, including ribosomal proteins and photosynthesis-related genes (**Figure S10**). This enrichment was observed in both species and suggests that egg cells provide embryos with a substantial “starter pack” of transcripts required for early cellular metabolism.

Finally, we compared the timing of maternal bias loss for single-copy orthologous genes between *Dictyota* and *Undaria* (**Figure 4, E and F**). A contingency analysis revealed a significant shift in the timing of this transition, with orthologous genes tending to lose maternal bias earlier in *Undaria* than in *Dictyota*. These results indicate that although the overall structure of maternal transcript inheritance is conserved between the two oogamous species, the timing of the transition toward balanced parental expression differs between them. Together, these analyses reveal therefore maternal dominance in the early embryonic transcriptomes of the oogamous species, followed by a progressive transition toward balanced parental expression, whereas the nearly isogamous *Scytosiphon* exhibits substantial biparental contributions from the very earliest stages.

## DISCUSSION

Fertilisation initiates the MZT, where maternal transcripts are degraded and replaced by zygotic transcription. We have shown that, in brown algae, the onset of this process is conserved but its dynamics vary across species, with gamete dimorphism playing a key role in shaping how the transition unfolds.

### Zygotic genome activation initiates early and independently of the N/C ratio

In animals, the N/C ratio has long been considered a central timer of ZGA ^35–37^: as rapid early cleavages progressively subdivide the egg cytoplasm, the rising N/C ratio titrates out maternal repressors and allows for large-scale transcription of the zygotic genome ^38^. Under this framework, embryos that begin with cytoplasm-poor cells and a high initial N/C ratio should activate their genomes earliest, while heavily provisioned, cytoplasm-rich eggs should delay ZGA until successive divisions have sufficiently raised nuclear content. A comparative study across 61 animal species proposed that ZGA timing can be quantitatively predicted from genome size and egg volume ^4^. This mechanism is likely ancestral to the holozoan lineage, as it also controls developmental timing in unicellular animal relatives like *Sphaeroforma arctica* ^37^.

Our results question the generality of the N/C ratio as a universal timer of ZGA. Despite substantial variation in gamete size and initial N/C ratio across the three brown algal species examined, all initiate zygotic genome activation very early after fertilisation and before the first cell division. This timing is inconsistent with predictions from the canonical animal N/C-ratio model ^36,38^ (**Figure S11**), indicating that alternative mechanisms permit early transcription in brown algae. Land plants display a qualitatively similar pattern of pre-cleavage activation, although their narrower N/C range currently limits formal comparisons.

Therefore, it appears that the N/C-ratio-dependent delay of ZGA is not a universal feature of embryogenesis, but rather a derived feature of the holozoan lineage. Although many fundamental aspects of embryonic development are conserved across multicellular eukaryotes ^32^, the regulatory logic underlying the MZT appears to have evolved independently in animals and other complex multicellular lineages.

### Oogamy shapes the dynamics, but not the onset, of zygotic genome activation

While the onset of ZGA is similarly early across species, its subsequent dynamics differ in ways that correlate with the degree of gamete dimorphism. In the two oogamous species, the number of newly transcribed zygotic genes increases approximately linearly across early development. In *Scytosiphon*, by contrast, the initial wave of ZGA, comparable in timing and gene numbers to that observed in the oogamous species, is followed by a second, larger wave at a later stage, affecting a greater number of genes.

The significance of this two-step pattern is not fully clear, but one possibility is that in an isogamous zygote, where both gametes contribute roughly equally to the cytoplasm, the first wave may partly fulfill a role analogous to maternal provisioning in oogamous species, sustaining the earliest developmental steps, while the more extensive second wave drives the bulk of zygotic reprogramming. Whether this reflects a functional partitioning between the two waves, or simply a difference in the tempo of genome activation linked to the isogamous starting condition, remains to be determined. One possibility is that chromatin configuration inherited from the gametes plays a role: in oogamous species, the egg nucleus enters fertilisation in a transcriptionally permissive state, potentially enabling rapid activation of the maternal genome, while in isogamous species both gametic nuclei may be slightly transcription permissive, but require more important decondensation before large-scale transcription can proceed.

### Gamete dimorphism determines the dynamics of parental transcript clearance

By quantifying transcript clearance and changes in parent-of-origin expression across development, we uncovered patterns of parental transcript persistence and turnover during early embryogenesis. Despite differences in oogamy and N/C ratio, parental transcript clearance reaches 50% between the 2- and 4-cell stages in all species. In contrast, the persistence of parentally biased transcripts varies with gamete dimorphism: both oogamous species show strong maternal bias during early development, but the transition toward balanced parental transcript abundance occurs at different rates. *Undaria* reaches near-balanced expression earlier than *Dictyota*, potentially reflecting differences in developmental timing and the extent of maternal provisioning. In *Scytosiphon*, transcripts from both parental alleles are detectable from the earliest stages examined. This pattern cannot be explained by cytoplasmic dilution alone and likely reflects both early transcriptional activation and more equal parental cytoplasmic contributions.

Taken together, these results indicate that gamete dimorphism is a key determinant of parental asymmetry during early embryogenesis, and suggest that the parental transcriptome functions as a “starter pack” comprising two functional categories: transcripts that disappear early and are likely required only for the first post-fertilisation developmental steps, and abundantly provisioned housekeeping transcripts that are slowly replaced as zygotic transcription ramps up.

In *Dictyota*, substantial transcriptome remodelling is already evident at the zygote stage, yet most genes remain strongly maternally biased. As these samples correspond to the estimated timing of karyogamy (∼1 h post-fertilisation ^25^), this suggests that early transcription is driven mainly by the maternal genome before full incorporation of the paternal genome, similar to pre-karyogamy activation of the female nucleus in *Oryza* ^11^. A previous study in Arabidopsis showed that some genes retain asymmetric transcription from one parental genome for several divisions after zygotic genome activation ^16^. Similarly, maternal transcript bias persists for several divisions in *Dictyota* and *Undaria*, but because these genes are mainly highly expressed housekeeping genes, this likely reflects incomplete transcript turnover rather than biased *de novo* transcription.

### No evidence of long-term parental imprinting in brown algae

In the latest developmental stages examined across all three species, we observe very little or no persistent parental bias in transcript abundance, suggesting that stable parental imprinting is unlikely in these species (although transient early maternal dominance could fall under broader definitions). Parental imprinting has been shown in species where the sporophyte is metabolically dependent on the gametophyte, as in *Marchantia polymorpha* ^6^. In *Undaria*, early sporophytes remain physically attached to the maternal gametophyte ^26^ which might be expected to promote prolonged maternal control over the embryonic transcriptome. Yet we detect no persistent maternal bias, nor any enhanced early maternal dominance relative to *Dictyota*, where sporophytes and gametophytes develop autonomously. This absence may be explained by the fact that while the parent gametophyte provides developmental cues to the developing sporophyte^26^, there is no evidence of metabolic exchange with its sporophytic offspring.

Finally, the temporal dynamics of maternal bias loss were consistent across crosses involving both closely and distantly related parental strains, in contrast to the variability reported in systems such as *Arabidopsis* ^16^. This suggests that parental contribution patterns are robust to genetic background in brown algae.

### A conserved developmental trajectory built on ancient regulatory genes

Despite the divergent MZT dynamics described above, and the substantial evolutionary distances separating the three species^39^, orthogroup-based and network-level analyses reveal a broadly conserved transcriptional trajectory across early development. Co-expression modules associated with developmental progression show partial conservation across species, and a common transcriptional trajectory emerges at the systems level. Critically, these conserved components are enriched in evolutionarily ancient genes, with deeply conserved gene families disproportionately represented among network hub genes, suggesting that an ancestral regulatory backbone underlies early development across the brown algal lineage.

This pattern is consistent with previous transcriptomic studies in brown algae ^32,40^ and is reminiscent of observations in animals, where comparative transcriptomics has shown that embryonic stages can align along conserved developmental axes even when gene-level expression and network composition diverge substantially, a phenomenon sometimes described as developmental system drift ^41,42^. Our results extend this principle to an independently evolved multicellular lineage, suggesting that the coupling between conserved developmental outcomes and flexible regulatory programs is a general feature of complex multicellular life.

Interestingly, we did not observe increased network similarity between the two oogamous species, *Dictyota* and *Undaria*. Their shared phenotypic characteristics may thus arise not from conserved co-expression modules, but rather from convergent evolution of distinct transcriptional programs associated with oogamy itself. This interpretation is consistent with previous phylogenetic analyses indicating independent origins of oogamy in Laminariales and Dictyotales ^34^.

## RESOURCE AVAILABILITY

Lead contactFurther information and requests for resources and reagents should be directed to and will be fulfilled by the lead contact, Susana M. Coelho.

### Data, code, and materials availability

BioProject ID PRJNA144506. Raw sequencing data have been deposited in the NCBI Sequence Read Archive under BioProject accession PRJNA144506. Sample-level accession numbers (SRR identifiers) are provided in **Table S3**. VCF files containing the parental SNPs used for allele-specific expression analysis are available in **Data S4**. Orthogroups and PC3 loadings are available in **Data S5** and **S6**.

## ACKNOWLEDGMENTS

We thank Jaruwatana Sodai Lotharukpong for helpful discussions Masakazu Hoshino for the sampling of *Scytosiphon* and *Undaria* Kobe strains, and Olivier De Clerck for the sampling of Dictyota Wimereux strains. We thank Katharina Hipp from the Electron Microscopy Facility at the Max Planck Institute for Biology Tubingen for assistance in the preparation of the TEM images. We thank Agnes Henschen for technical support, Andrea Belkacemi for assistance with algal cultures, and Dafne Ibarra for assistance with library preparation.

This work was supported by the Max Planck Gesellschaft, the ERC grant n. 638240 (SMC), the Moore Foundation grant GBMF11489 (SMC) and the Bettencourt-Schuller Foundation.

## AUTHOR CONTRIBUTIONS

S.M.C. conceived and designed the experiments, supervised the project and acquired funding. K.A.B. and M.Z. performed experiments. K.A.B. performed algae culture and sample collection (genetic crosses and micro-dissections for tissue collection). K.A.B., P.R., M.Z., R.A.B., F.H., S.M.C. analysed and interpreted data; K.A.B., P.R. and F.B.H. curated the data. K.A.B. and P.R. performed data analysis and visualization with support from S.M.C. K.A.B., P.R. and S.M.C. wrote the original draft. K.A.B., P.R., R.A.B. and S.M.C. reviewed and edited the manuscript.

## DECLARATION OF INTERESTS

Authors declare that they have no competing interests.

### Supplementary Information

Materials and Methods

Figures. S1 to S11

Tables S1 to S8

Data S1 to S6

## Notes

### Competing Interest Statement

The authors have declared no competing interest.

### Summary of Updates

We added a new supplemental figure

## References

1. Tadros, W., and Lipshitz, H.D. (2009). The maternal-to-zygotic transition: a play in two acts. Development 136, 3033–3042. 10.1242/dev.033183.

2. Schulz, K.N., and Harrison, M.M. (2019). Mechanisms regulating zygotic genome activation. Nat. Rev. Genet. 20, 221–234. 10.1038/s41576-018-0087-x.

3. Kojima, M.L., Hoppe, C., and Giraldez, A.J. (2025). The maternal-to-zygotic transition: reprogramming of the cytoplasm and nucleus. Nat. Rev. Genet. 26, 245–267. 10.1038/s41576-024-00792-0.

4. Campo-Bes, I., Mantica, F., Permanyer, J., Rodriguez-Marin, C., Guynes, K., Senar-Serra, T., Quiroga-Artigas, G., Liang, Y., Carrillo-Baltodano, A.M., Cruz, J., et al. (2026). Evolutionary landscapes of zygotic genome activation across animals. Preprint at Evolutionary Biology, http://doi.org/10.64898/2026.04.16.718233 10.64898/2026.04.16.718233.

5. Guo, Q., Xu, F., Song, S., Kong, S., Zhai, F., Xiu, Y., Liu, D., Li, M., Lian, Y., Ding, L., et al. (2025). Allelic transcriptomic profiling identifies the role of PRD-like homeobox genes in human embryonic-cleavage-stage arrest. Dev. Cell 60, 1290–1303.e6. 10.1016/j.devcel.2024.12.031.

6. Montgomery, S.A., Hisanaga, T., Wang, N., Axelsson, E., Akimcheva, S., Sramek, M., Liu, C., and Berger, F. (2022). Polycomb-mediated repression of paternal chromosomes maintains haploid dosage in diploid embryos of Marchantia. eLife 11, e79258. 10.7554/eLife.79258.

7. Weijers, D., Geldner, N., Offringa, R., and Jürgens, G. (2001). Early paternal gene activity in Arabidopsis. Nature 414, 709–710. 10.1038/414709a.

8. Autran, D., Baroux, C., Raissig, M.T., Lenormand, T., Wittig, M., Grob, S., Steimer, A., Barann, M., Klostermeier, U.C., Leblanc, O., et al. (2011). Maternal Epigenetic Pathways Control Parental Contributions to Arabidopsis Early Embryogenesis. Cell 145, 707–719. 10.1016/j.cell.2011.04.014.

9. Vielle-Calzada, J.-P., Baskar, R., and Grossniklaus, U. (2000). Delayed activation of the paternal genome during seed development. Nature 404, 91–94. 10.1038/35003595.

10. Bayer, M., Nawy, T., Giglione, C., Galli, M., Meinnel, T., and Lukowitz, W. (2009). Paternal Control of Embryonic Patterning in Arabidopsis thaliana. Science 323, 1485–1488. 10.1126/science.1167784.

11. Toda, E., Koshimizu, S., Kinoshita, A., Higashiyama, T., Izawa, T., Yano, K., and Okamoto, T. (2025). Transcriptional dynamics during karyogamy in rice zygotes. Development 152, DEV204497. 10.1242/dev.204497.

12. Zhao, P., Zhou, X., Shen, K., Liu, Z., Cheng, T., Liu, D., Cheng, Y., Peng, X., and Sun, M.-X. (2019). Two-Step Maternal-to-Zygotic Transition with Two-Phase Parental Genome Contributions. Dev. Cell 49, 882–893.e5. 10.1016/j.devcel.2019.04.016.

13. Chen, J., Strieder, N., Krohn, N.G., Cyprys, P., Sprunck, S., Engelmann, J.C., and Dresselhaus, T. (2017). Zygotic Genome Activation Occurs Shortly after Fertilization in Maize. Plant Cell 29, 2106–2125. 10.1105/tpc.17.00099.

14. Anderson, S.N., Johnson, C.S., Chesnut, J., Jones, D.S., Khanday, I., Woodhouse, M., Li, C., Conrad, L.J., Russell, S.D., and Sundaresan, V. (2017). The Zygotic Transition Is Initiated in Unicellular Plant Zygotes with Asymmetric Activation of Parental Genomes. Dev. Cell 43, 349–358.e4. 10.1016/j.devcel.2017.10.005.

15. Toda, E., Koshimizu, S., Kinoshita, A., Higashiyama, T., Izawa, T., Yano, K., and Okamoto, T. (2025). Transcriptional dynamics during karyogamy in rice zygotes. Development 152, DEV204497. 10.1242/dev.204497.

16. Alaniz-Fabián, J., Xiang, D., Del Toro-De León, G., Gao, P., Abreu-Goodger, C., Datla, R., and Gillmor, C.S. (2025). A maternal transcriptome bias in early Arabidopsis embryogenesis. Development 152, dev204449. 10.1242/dev.204449.

17. Gueno, J., Borg, M., Bourdareau, S., Cossard, G., Godfroy, O., Lipinska, A., Tirichine, L., Cock, J.M., and Coelho, S.M. (2022). Chromatin landscape associated with sexual differentiation in a UV sex determination system. Nucleic Acids Res. 50, 3307–3322. 10.1093/nar/gkac145.

18. Liu, P., Vigneau, J., Craig, R.J., Barrera-Redondo, J., Avdievich, E., Martinho, C., Borg, M., Haas, F.B., Liu, C., and Coelho, S.M. (2024). 3D chromatin maps of a brown alga reveal U/V sex chromosome spatial organization. Nat. Commun. 15, 9590. 10.1038/s41467-024-53453-5.

19. Vigneau, J., Lotharukpong, J.S., Liu, P., Luthringer, R., Lombard, B., Loew, D., Haas, F.B., Borg, M., and Coelho, S.M. (2025). Rewiring of chromatin regulation underlies the evolution of brown algal multicellularity. Preprint at Evolutionary Biology, http://doi.org/10.1101/2025.09.16.676480 10.1101/2025.09.16.676480.

20. Bukhanets, V., Batista, R.A., Haas, F.B., Luthringer, R., Kushkush, J., Zheng, M., Hipp, K., Alva, V., Martinho, C., and Coelho, S.M. (2026). Germline fate determination by a single ARGONAUTE protein in Ectocarpus. Proc. Natl. Acad. Sci. 123, e2518712123. 10.1073/pnas.2518712123.

21. Heesch, S., Serrano-Serrano, M., Luthringer, R., Peters, A.F., Destombe, C., Cock, J.M., Valero, M., Roze, D., Salamin, N., and coelho, susanam (2019). Evolution of life cycles and reproductive traits: insights from the brown algae. bioRxiv. 10.1101/530477.

22. Bringloe, T.T., Starko, S., Wade, R.M., Vieira, C., Kawai, H., Clerck, O.D., Cock, J.M., Coelho, S.M., Destombe, C., Valero, M., et al. (2020). Phylogeny and evolution of the brown algae. Crit. Rev. Plant Sci. 39, 281–321. 10.1080/07352689.2020.1787679.

23. Luthringer, R., Cormier, A., Peters, A.F., Cock, J.M., and Coelho, S.M. (2015). Sexual dimorphism in the brown algae. Perspectives in Phycology 1, 11–25.

24. Bogaert, K.A., Zakka, E.E., Coelho, S.M., and De Clerck, O. (2022). Polarization of brown algal zygotes. Semin. Cell Dev. Biol., S1084-9521(22)00075-1. 10.1016/j.semcdb.2022.03.008.

25. Phillips, J.A., Clayton, M., Maier, I., Boland, W., and Muller, D. (1990). Sexual Reproduction in Dictyota-Diemensis (dictyotales, Phaeophyta). PHYCOLOGIA 29, 367–379. 10.2216/i0031-8884-29-3-367.1.

26. Dries, E., Meyers, Y., Liesner, D., Gonzaga, F.M., Becker, J.F.M., Zakka, E.E., Beeckman, T., Coelho, S.M., De Clerck, O., and Bogaert, K.A. (2024). Cell wall-mediated maternal control of apical–basal patterning of the kelp Undaria pinnatifida. New Phytol. 243, 1887–1898. 10.1111/nph.19953.

27. Bogaert, K.A., Beeckman, T., and De Clerck, O. (2017). Two-step cell polarization in algal zygotes. Nat. Plants 3, 16221. 10.1038/nplants.2016.221.

28. Klochkova, T.A., Motomura, T., Nagasato, C., Klimova, A.V., and Kim, G.H. (2019). The role of egg flagella in the settlement and development of zygotes in two Saccharina species. Phycologia 58, 145–153. 10.1080/00318884.2018.1528804.

29. Wyatt, C.D.R., Pernaute, B., Gohr, A., Miret-Cuesta, M., Goyeneche, L., Rovira, Q., Salzer, M.C., Boke, E., Bogdanovic, O., Bonnal, S., et al. (2022). A developmentally programmed splicing failure contributes to DNA damage response attenuation during mammalian zygotic genome activation. Sci. Adv. 8, eabn4935. 10.1126/sciadv.abn4935.

30. Barrera-Redondo, J., Lotharukpong, J.S., Drost, H.-G., and Coelho, S.M. (2023). Uncovering gene-family founder events during major evolutionary transitions in animals, plants and fungi using GenEra. Genome Biol. 24, 54. 10.1186/s13059-023-02895-z.

31. Barrera-Redondo, J., Lipinska, A.P., Liu, P., Dinatale, E., Cossard, G., Bogaert, K., Hoshino, M., Craig, R.J., Avia, K., Leiria, G., et al. (2025). Origin and evolutionary trajectories of brown algal sex chromosomes. Nat. Ecol. Evol. 10.1038/s41559-025-02838-w.

32. Lotharukpong, J.S., Zheng, M., Luthringer, R., Liesner, D., Drost, H.-G., and Coelho, S.M. (2024). A transcriptomic hourglass in brown algae. Nature 635, 129–135. 10.1038/s41586-024-08059-8.

33. Silberfeld, T., Leigh, J.W., Verbruggen, H., Cruaud, C., de Reviers, B., and Rousseau, F. (2010). A multi-locus time-calibrated phylogeny of the brown algae (Heterokonta, Ochrophyta, Phaeophyceae): Investigating the evolutionary nature of the “brown algal crown radiation”. Mol. Phylogenet. Evol. 56, 659–674. 10.1016/j.ympev.2010.04.020.

34. Heesch, S., Serrano-Serrano, M., Barrera-Redondo, J., Luthringer, R., Peters, A.F., Destombe, C., Cock, J.M., Valero, M., Roze, D., Salamin, N., et al. (2021). Evolution of life cycles and reproductive traits: insights from the brown algae. J. Evol. Biol. n/a. 10.1111/jeb.13880.

35. Newport, J., and Kirschner, M. (1982). A major developmental transition in early xenopus embryos: II. control of the onset of transcription. Cell 30, 687–696. 10.1016/0092-8674(82)90273-2.

36. Jukam, D., Kapoor, R.R., Straight, A.F., and Skotheim, J.M. (2021). The DNA-to-cytoplasm ratio broadly activates zygotic gene expression in Xenopus. Curr. Biol. 31, 4269–4281.e8. 10.1016/j.cub.2021.07.035.

37. Olivetta, M., and Dudin, O. (2023). The nuclear-to-cytoplasmic ratio drives cellularization in the close animal relative Sphaeroforma arctica. Curr. Biol. 33, 1597–1605.e3. 10.1016/j.cub.2023.03.019.

38. Amodeo, A.A., Jukam, D., Straight, A.F., and Skotheim, J.M. (2015). Histone titration against the genome sets the DNA-to-cytoplasm threshold for the Xenopus midblastula transition. Proc. Natl. Acad. Sci. 112. 10.1073/pnas.1413990112.

39. Choi, S.-W., Graf, L., Choi, J.W., Jo, J., Boo, G.H., Kawai, H., Choi, C.G., Xiao, S., Knoll, A.H., Andersen, R.A., et al. (2024). Ordovician origin and subsequent diversification of the brown algae. Curr. Biol. 34, 740–754.e4. 10.1016/j.cub.2023.12.069.

40. Lipinska, Cormier A., Luthringer, R., Peters, A.F., Corre, E., Gachon, C.M.M., Cock, J.M., and Coelho, S.M. (2015). Sexual dimorphism and the evolution of sex-biased gene expression in the brown alga Ectocarpus. Mol. Biol. Evol. 32, 1581–1597. 10.1093/molbev/msv049.

41. True, J.R., and Haag, E.S. (2001). Developmental system drift and flexibility in evolutionary trajectories. Evol. Dev. 3, 109–119. 10.1046/j.1525-142x.2001.003002109.x.

42. McColgan, Á., and DiFrisco, J. (2024). Understanding developmental system drift. Development 151, dev203054. 10.1242/dev.203054.

