## Supplemental Materials for "Evolutionary flexibility in the maternal-to-zygotic transition reveals alternative routes to embryogenesis across eukaryotes"

MATERIAL & METHODS

### Biological material

Three brown algal species were used in this study: *Dictyota dichotoma*, *Undaria pinnatifida*, and *Scytosiphon promiscuus*. Strain identities and origins are provided in **Table S2**. *Dictyota dichotoma* strains were maintained in PES-enriched seawater under white light at 20 °C with a 12h:12h (light:dark) photoperiod. Gamete release and fertilisation were induced by exposure to light at the start of the photoperiod. Gamete release commenced approximately 20 min after the onset of light and continued for ~15 min. Eggs and sperm were released simultaneously, enabling immediate fertilisation in the culture medium. Eggs, zygotes and embryos were staged and collected by manual picking with a pipette immediately prior to sampling.

*Undaria pinnatifida* strains were maintained in PES medium under white light at 20°C with a 18h:8h (light:dark) dark photoperiod. Male and female gametophytes were fragmented and allowed to develop for approximately 6–7 days, after which the fertile period begins. Gametes were released spontaneously at the onset of the dark period. Fertilisation and zygote isolation by microsurgical blade were performed as described in^25^.

*Scytosiphon promiscuus* strains were maintained in PES medium under white light at 12°C with a 12h light : 12h dark photoperiod. A fertilisation protocol was developed to obtain large synchronised populations of zygotes. Male and female gametes were released by refreshing the culture medium. Fertilisation was initiated by combining small droplets of male and female gamete suspensions under unilateral light. Motile gametes accumulated at the illuminated edge of the droplet, where fertilisation occurred; upon fertilisation, zygotes settled and formed dense patches. Regions of high zygote density were selected, and any remaining unfertilised gametes - identified by two eyespots, and smaller cell size - were removed carefully by micromanipulation of a hair. Fertilised zygotes were allowed to develop under the culture conditions described above.  The patterning-stage sample of the *Scytosiphon* cross was excluded from all downstream analyses. This sample, harvested from a separate gamete-release period, displayed atypically slow development, raising suspicion of accidental inclusion of female parthenotes. Strain-specific read sorting subsequently confirmed this contamination (**Figure** **S9b**). The corresponding reciprocal-cross sample showed no such signature and was retained.

For all species, individual gametes embryos or zygotes were transferred into 4 µl lysis buffer, squashed using a 6 mm coverslip, and the coverslip was washed with an additional 4 µl lysis buffer. The resulting lysate was directly processed for low-input RNA library preparation (see RNA extraction, library preparation and sequencing.

### RNA extraction, library preparation and sequencing

Algal material from early stages with low cell count per individual were extracted using a low-input approach (referred to as Direct Low Input in **Table S3**). Briefly, embryos were sorted manually to ensure developmental synchronicity, and the sorted material was lysed directly by pressing between glass slides in the lysis buffer provided with the NEBNext Single Cell/Low Input RNA Library Prep Kit for Illumina (NEB, #6420), which was also used for library preparation according to the manufacturer's instructions.

Adult multicellular algal material was extracted using a CTAB-based RNA extraction protocol (referred to as Bulk in **Table S3**). Adult multicellular algal material was quickly brushed and rinsed with filtered autoclaved natural seawater and transferred into 1.5 ml low-bind Eppendorf tubes. Snap-frozen material was dry-ground with a pestle in liquid N₂/dry ice and mixed with 750 µl of freshly prepared RNA extraction buffer (100 mM Tris-HCl pH 8.0, 1.4 M NaCl, 2% CTAB, 20 mM EDTA pH 8.0, 1% β-mercaptoethanol, 2% polyvinylpyrrolidone) preheated to 65°C, followed by addition of 250 µl 5 M NaCl. An equal volume of chloroform:isoamylalcohol (24:1) was added, mixed and centrifuged at 10,000g for 15 min at 4°C. The aqueous phase was re-extracted with 250 µl ethanol and an equal volume of chloroform:isoamylalcohol (24:1). RNA was precipitated overnight at −20°C with LiCl to a final concentration of 4 M and 1% β-mercaptoethanol. The pellet was collected by centrifugation at >18,000g for 45–60 min at 4°C, washed with cold 70% ethanol, air dried and dissolved in 30 µl RNase-free H₂O. Residual DNA was removed using the TURBO DNase Kit (Thermo Fisher, AM1907) according to the manufacturer's instructions. RNA concentration and quality were assessed using a Qubit RNA BR Assay Kit (Invitrogen, Q10210) and an RNA Nano Bioanalyzer chip (Agilent, 5067-1511). Libraries were prepared using the NEBNext Ultra II Directional RNA Library Prep Kit with Sample Purification Beads (NEB, #7765) according to the manufacturer's instructions.
Libraries were sequenced on an Illumina platform as 150 bp paired-end reads, either in-house or by Azenta Life Sciences. Sequencing platform details per sample are provided in **Table S3**. Data is available under BioProject ID PRJNA144506.

### N/C ratio calculation

N/C ratios were calculated as two times the haploid genome size (Mb) divided by zygote cytoplasmic volume (µm³), expressed as Mb DNA per µm³, reflecting the diploid DNA content of the zygote. Haploid genome sizes were obtained directly from published genome assemblies: *Scytosiphon promiscuus* and *Dictyota dichotoma* from^29,30^ *Undaria pinnatifida* from^43^ and animal and plant genomes from their respective published assemblies (**Table S2**). Zygote volumes for brown algae were obtained from published measurements: *Scytosiphon promiscuus* from^44^ *Undaria pinnatifida* from^25^ and *Dictyota dichotoma* from^26^. Zygote volumes for comparative animal and plant species were obtained from the literature as indicated in **Table S1**. All N/C values are plotted on a log scale.

### TEM (Transmission electron microscopy)

For TEM, gamete samples were high-pressure frozen (HPF Compact 03, Engineering Office M. Wohlwend GmbH) as in^19,45^, freeze-substituted (AFS2, Leica Microsystems) with 0.2% OsO_4_ and 0.1% uranyl acetate in acetone containing 1.5% H_2_O as substitution medium and embedded in Epon. For TEM, ultrathin sections were stained with uranyl acetate and lead citrate and analyzed with a JEM-2100Plus (Jeol) operated at 200 kV.

### RNA-seq

Paired-end RNA-seq reads were processed using a custom analysis pipeline. Read quality was assessed before and after preprocessing with FastQC. Adapter trimming and quality filtering were performed using Cutadapt, applying a minimum Phred quality score of 25 and discarding reads shorter than 40 bp. Adapter sequences from TruSeq, Nextera, and custom libraries were screened. Trimmed reads were aligned to the reference genome using STAR, using either local or end-to-end alignement as specified. Gene annotations (GTF) were supplied during alignment, allowing introns up to 50 kb. Coordinate-sorted BAM files and STAR-derived gene-level counts were generated. Gene expression was quantified using featureCounts (Subread v2.0.5) in paired-end mode, assigning reads to mRNA features defined in GFF annotations. PCR duplicates were identified and removed using samtools markdup^46^, and quantification was repeated on duplicate removed alignments. Alignment quality and read distribution across genomic features were assessed using Qualimap (v2.3) in RNA-seq mode. Reads mapping to predefined organellar or rRNA contigs were identified from duplicate-removed BAM files and removed at the BAM level. Cleaned alignments were used for downstream analyses.

### Differential gene expression analysis

Raw read counts from featureCounts were imported into R (v4.2) for differential expression analysis using DESeq2^47^. Analyses were performed independantly for each of the three brown algal species: *Dictyota*, *Scytosiphon*, and *Undaria*. Samples corresponding to female gametes, zygote, polarisation, 2-celled, 4-celled, and patterning. Genes with a rowsummed count of ≤10 across all sam-ples were removed prior to model fitting. DESeq2 was run with a single-factor design (~time_point), with female gametes set as the refernce level. For each species, pairwise contrasts were extracted compar-ing each developmental stage with gametes set as the reference level (female gametes for the oogamous *Dictyota* and *Undaria*; mixed gametes for the isogamous *Scytosiphon*). Genes were classified as differentially expressed at an adjusted p-value (Benjamini-Hochberg) < 0.05 and log₂ fold change ≥ 1. To characterise the temporal dynamics of differential expression, genes were assigned to the earliest developmental stage at which they first apeared as a DEG. Mean VST-transformed expression values across all DEGs at each stage were partitioned into low, medium and high, to assess whether early- or late-appearing DEGs differed in overal expression level. Transcriptome-wide VST expression tertile boundaries were (6.44 , 8.57) (*Dictyota*), (5.01, 8.25) (*Scytosiphon*), and (5.33, 7.57) (*Undaria*).

Principal component analysis was performed on VST-normalised counts (DESeq2, blind = TRUE) using the 500 most variably expressed genes, selected by variance across all samples. PCA was computed using DESeq2's plotPCA function^47^.

### Transcript clearance

Transcript clearance was quantified by identifying genes expressed in the female gamete (VST-normalised counts ≥ 2 and DESeq2-normalised counts ≥ 10) that were significantly downregulated at any subsequent developmental stage (adjusted p-value < 0.05, log₂FC ≤ −1). Genes were retained if their mean VST expression decreased monotonically across stages (allowing no increase > 0.4 VST units between consecutive stage means). For each retained gene, the clearance stage was defined as the earliest stage at which log₂(normalised counts + 1) dropped below 10% of the reference level by the last stage. Across the three species, the number of genes meeting all clearance criteria (expressed in the female gamete, significantly downregulated, and showing a monotonically decreasing trajectory) was 421 (*Dictyota*), 437 (*Undaria*), and 509 (*Scytosiphon*).

### Orthogroup identification

Orthologous groups were identified across the three brown algal species using OrthoFinder v2.5.5^48^ with default parameters, using the predicted proteomes of *Dictyota dichotoma*^30^*, Undaria pinnatifida*^43^*,* and *Scytosiphon promiscuus*^30,49^ as input. The resulting single-copy orthogroups (**Data S5**) were used for cross-species PCA of conserved transcriptional dynamics, assessment of co-expression module conservation across species.

### co-expression network analysis

Weighted gene co-expression network analysis. For each species, signed co-expression networks were constructed independently using WGCNA^50^. Variance-stabilized expression matrices (DESeq2) were used as input, restricted to genes passing the expression threshold described above. Soft-thresholding powers were selected per species based on the scale-free topology criterion (R² > 0.80). Modules were identified using the blockwiseModules function with default parameters, and module eigengenes were computed as the first principal component of each module's expression matrix. The grey module, containing unassigned genes, was excluded from downstream analyses.

### Co-expression network conservation across species

To assess whether co-expression modules are conserved across the three brown algal species, we determined gene-set overlap and conservation of temporal expression dynamics during MZT using the orthology group geneset.

*Gene-set overlaps.* For each pair of species, orthologs were mapped to their respective module assignments. Enrichment of module overlap was tested with one-sided Fisher's exact tests on module-by-module contingency tables, with p-values corrected for multiple testing (Benjamiini–Hochberg). The Jaccard index and odds ratio were computed for each module pair. Only pairs passing statistical significance (adjusted p < 0.05), a minimum overlap of 50 orthologous genes, and a minimum, odds ratio ≥ 1.5 were retained.

*Eigengene trajectory conservation.* For module pairs identified as significantly overlapping at eigengene expression trajectories were compared across the shared developmental axis. Per-stage mean eigengene values were computed for each module, and trajectory similarity was quantified by Pearson correlation. Module pairs with |ρ| > 0.6 and p < 0.05 were considered to show conserved temporal expression dynamics.

### Phylostratigraphic analysis

Gene ages for each of the three brown algal species were obtained from published phylostratigraphic datasets. Ages for *Dictyota dichotoma* and *Scytosiphon promiscuus* were taken from^30^, and ages for *Undaria pinnatifida* were taken from^51^. Each gene was assigned a phylostratum (age rank), and phylostrata were collapsed into three simplified age classes - Old, Mid, and Young - using species-specific cut-offs adapted to the depth of each phylostratigraphic tree. For *Dictyota* (11 phylostrata), Old = PS 1–2, Mid = PS 3–10, Young = PS 11. For *Scytosiphon* (14 phylostrata), Old = PS 1–2, Mid = PS 3–13, Young = PS 14. For *Undaria* (13 phylostrata), Old = PS 1–2, Mid = PS 3–11, Young = PS 12–13.

Enrichment of age classes within WGCNA co-expression modules was tested using two-sided Fisher's exact tests against the background distribution of age classes across all age-annotated expressed genes, with Benjamini–Hochberg correction for multiple testing (**Table S4**). The same test was applied to the top 25% of genes by absolute module membership (|MM|) to assess age-class enrichment among hub genes, using each module's own age composition as the background (**Table S5**). Genes without an assigned age were excluded from enrichment tests. The background transcriptome was defined as genes with ≥100 raw counts in at least two samples.

### Annotations

GO annotation and functional enrichment along the PC3 developmental axis. Per-species gene-level GO annotations were obtained by combining InterProScan v-5.74-105.0^52^ and eggNOG-mapper outputs, expanded to include ancestor terms (GO.db), and mapped to the generic GO slim (goslim_generic.obo, GSEABase). Each single-copy orthogroup inherited the union of GO slim terms from its three member genes; KEGG pathway annotations were derived analogously from eggNOG-mapper-v2^53^ KO identifiers.

Rank-based functional enrichment along the PC3 axis. To exploit the full distribution of PC3 loadings rather than a fixed cut-off, a rank-based competitive enrichment test was performed by comparing the PC3 loadings of orthogroups annotated to each term with the background distribution of loadings across all PCA orthogroups using a two-sided Wilcoxon rank-sum test (minimum term size: 15 for BP and MF, 5 for CC). P-values were adjusted for multiple testing using the Benjamini–Hochberg (BH) procedure within each ontology. Tests were performed independently for GO Biological Process (BP), Molecular Function (MF) and Cellular Component (CC) ontologies, and for KEGG pathways. Cleaned full and slim GO-term datasets can be found in **Datasets 1-2**.

### Cross-species transcriptome distance metrics

For each species pair, pairwise distances between developmental stages were computed on shared single-copy orthogroups (Figure S5). The TPM values were log₂(x + 1)-transformed, replicate samples aggregated to stage-level profiles by per-orthogroup median, and orthogroups with SD ≤ 0.1 across all samples removed. Jensen–Shannon divergence was computed on stage profiles after sample-wise probability normalisation, which removes absolute expression magnitude by construction and captures differences in the shape of the global expression distribution. Pearson correlations and Euclidean and Manhattan distances were computed after per-orthogroup mean-centring within each species, removing lineage-specific baseline offsets and isolating relative developmental dynamics. Pearson values are reported as r.

### Identification of newly-expressed genes

For each species (*Scytosiphon*, *Undaria*, and *Dictyota*), normalized counts were summarized as the median across biological replicates for each developmental stage. Genes were considered expressed at a given stage when their median normalised count exceeded the normalized count threshold. The earliest developmental stage at which expression exceeded the threshold (5 normalized counts) was identified. Genes already expressed at the reference stage (female gametes for *Undaria* and *Dictyota*, gametes for *Scytosiphon*) were excluded, and the remaining genes were classified as first expressed after fertilization. These genes were further grouped by developmental stage and binned according to their median normalized abundance (low: 5–10, moderate: 10–100, high: ≥100 normalized counts).

### Strain genotyping and verification

Strain identity was verified for each of the RNA-seq samples used for strain-specific SNP determination or parental expression analysis in the crosses (samples listed either in **Table S3** or **Table S7**).  RNA-seq reads were trimmed using Trimmomatic version 0.39^54^ standard Illumina TruSeq paired-end adapters. Low-input samples were trimmed with the parameters: ILLUMINACLIP:2:5:5 SLIDINGWINDOW:4:15 HEADCROP:7 MINLEN:30, and bulk samples with: ILLUMINACLIP:2:30:10:2:keepBothReads SLIDINGWINDOW:4:20 MINLEN:50.

Trimmed reads were mapped to the reference genomes using STAR version 2.7.11b^55^ in two-pass mode. Genome indices were built with --runMode genomeGenerate prior to mapping. Mapping parameters included --outSAMtype BAM SortedByCoordinate, --outSAMunmapped Within, --outSAMattributes All, --outSAMmapqUnique 60, and read group information added via --outSAMattrRGline.

Variant calling was performed using GATK version 4.6.1.0^56^ following the pipeline described in ^57^ modified to produce GVCFs. Briefly, an initial round of variant calling was performed with HaplotypeCaller at the sample's true ploidy. High-quality variants were selected retaining variants within the 90th percentile of quality scores. VCFs were indexed at each step using IndexFeatureFile. Base quality score recalibration was then performed by sequentially applying BaseRecalibrator and ApplyBQSR using these high-quality variants as a known sites resource. Recalibration was assessed with AnalyzeCovariates, and recalibrated reads were exported with PrintReads.

Recalibrated BAM files from all samples of the same species were used for final GVCF calling with HaplotypeCaller in GVCF mode at ploidy 2. For each species, GVCFs were combined with CombineGVCFs and jointly genotyped with GenotypeGVCFs at ploidy 2.

Variants were filtered in two steps. First, VariantFiltration from GATK was applied with the following expression filters: FS > 60.0 (strand bias), QD < 2.0 (quality by depth), MQ < 40.0 (mapping quality), QUAL < 30.0 (variant quality), and DP < 10n where n is the number of samples (minimum total depth), with missing values treated as failing, and variants overlapping the SDR as defined in^30^ were additionally masked. Second, BCFtools version 1.22^58^ was used to retain only biallelic SNPs passing all filters (-v snps -m2 -M2), with species-specific thresholds for missing data, minor allele count, and number of heterozygous genotypes per site: for *Scytosiphon* *promiscuus*, no missing genotypes allowed (-i 'N_MISSING == 0'), MAC ≥ 6, and between 6 and 24 heterozygous genotypes per site; for *Undaria pinnatifida*, a maximum of 5 missing genotypes (-i 'N_MISSING <= 5'), MAC ≥ 10, and between 10 and 26 heterozygous genotypes; for *Dictyota dichotoma*, a maximum of 10 missing genotypes (-i 'N_MISSING <= 10'), MAC ≥ 15, and between 20 and 45 heterozygous genotypes. The heterozygosity bounds were set to retain sites consistent with the expected proportion of diploid hybrid samples in each dataset while excluding potentially paralogous sites showing excess heterozygosity.

The resulting VCF was processed in R using vcfR version 1.15.0^59^. Genotypes were encoded numerically (0, 1, 2 for homozygous reference, heterozygous, and homozygous alternative respectively), and PCA was performed in R using prcomp with scaling and centering to verify strain identity across all available samples, including both haploid parental samples and diploid hybrid samples from the crosses (**Figure S6, S7, and S8**).

### Calling of strain-specific SNPs

For each strain, haploid RNA-seq samples were used to perform the variant calling; when available, multiple haploid life stages were included (**Table S7**). Trimmed, mapped, and recalibrated reads from the previous section were merged for each haploid strain using SAMtools version 1.7^46,60^ with the merge function, and a common read group was assigned using addreplacerg. Merged BAMs were used for final GVCF calling with HaplotypeCaller in GVCF mode at ploidy 2. For each pair of strains, GVCFs were combined with CombineGVCFs and jointly genotyped with GenotypeGVCFs at ploidy 2.

Variants were filtered in two steps. First, VariantFiltration from GATK was applied with the following expression filters: FS > 60.0 (strand bias), QD < 2.0 (quality by depth), MQ < 40.0 (mapping quality), QUAL < 30.0 (variant quality), and DP < 20 (read depth), with missing values treated as failing. Variants overlapping the sex-determining region (SDR) as defined in ^30^ were additionally removed. Second, BCFtools version 1.22^60^ was used to retain only biallelic SNPs (-v snps -m2 -M2) with no missing genotypes (-i 'N_MISSING == 0'), a minimum read depth of 10 in each strain (-i 'MIN(FORMAT/DP) >= 10'), and where the two strains carried different alleles (combination of -i 'MAC >= 2' and -i 'N_PASS(GT="het") = 0').

The resulting VCF was processed in R using vcfR version 1.15.0^59^ to produce a dual-hybrid VCF for use with SNPsplit. Sites where both strains were homozygous and carried different alleles were retained, the INFO field was cleared, and a filter flag (FI=1) was appended to the genotype fields. Processed cross specific VCFs can be found in supplementary datasets.

### Sorting of strain-specific reads

Strain-specific read sorting was performed using SNPSplit version 0.6.0^61^. Masked versions of each reference genome were generated using the SNPSplit_genome_preparation function in dual-hybrid mode, which masks polymorphic sites for both strains simultaneously, using the VCF files generated at the previous step. Masked genomes were indexed with STAR using the splice junctions identified during the 2-pass mapping described above. Trimmed reads were mapped to these masked genomes in end-to-end mode with the following parameters: --outSAMtype BAM SortedByCoordinate --outSAMattributes All --outSAMunmapped Within --outFilterMultimapNmax 1 --outFilterMismatchNmax 999 --outFilterMismatchNoverLmax 0.04 --alignIntronMin 20 --alignIntronMax 50000 --alignMatesGapMax 50000 --alignEndsType EndToEnd. Resulting BAM files were indexed with SAMtools version 1.21^46,58^ and reads were assigned to maternal or paternal genomes using SNPSplit. Read counts per gene were obtained using featureCounts from the Subread package version 2.0.1^62^ counting reads mapping to exons with options -s 2 -p. Count tables were imported into R for downstream analysis.

### Analysis of maternal bias

In each individual sample, transcript per million (TPM) values were computed for each gene as read counts normalised by gene length and sequencing depth. In addition, maternal and paternal sorted read counts were normalised by the total number of sorted reads, which was provided to edgeR version 4.6.3^63^ as a custom library size. For each gene, the maternal fraction was computed as maternal reads / (maternal + paternal reads). For each developmental stage, genome bias in transcript abundance was assessed using edgeR: genes with insufficient counts were removed using filterByExpr with default parameters, dispersion was estimated with estimateDisp, and differences in transcript abundance between maternal and paternal genomes were tested using exactTest. Based on the exact test results, genes were classified into the following categories: genes with FDR < 0.05 and logFC < -1 in the paternal versus maternal comparison were classified as maternally biased; genes with FDR < 0.05 and logFC > 1 as paternally biased; genes with no test result (filtered out by filterByExpr) and mean TPM < 5 were classified as not expressed; genes with FDR ≥ 0.05 or |logFC| ≤ 1 were classified as unbiased; and genes with no test result but mean TPM ≥ 5 were classified as undetermined. The computed values can be found in supplementary datasets.

To investigate the relationship between parental bias dynamics and transcript abundance across development in the two oogamous species, genes were grouped based on the stage at which they first showed unbiased transcript abundance across both reciprocal crosses. For each gene and developmental stage, a consensus bias category was derived from the two reciprocal crosses: genes classified as maternally biased in both crosses were assigned as maternally biased; genes maternally biased in only one cross were classified as maternally oriented; and genes not maternally biased in either cross were classified as unbiased. Note that the egg cell stage was set as maternally biased for all genes by definition, as no paternal contribution is expected at this stage. Based on these consensus categories, each gene was assigned to a group corresponding to the first developmental stage at which it transitioned to unbiased expression; genes that remained maternally biased throughout all stages were assigned to a "never" group. Gene Ontology enrichment analysis was performed for each group defined in this way using the enricher function from clusterProfiler version 4.16.0^64^, with the set of genes having both SNP information and GO annotations as the background universe.

To compare parental bias dynamics between the two oogamous species, the first unbiased stage assignments of one-to-one orthologous gene pairs were compared. Due to the small number of one-to-one orthologous gene pairs containing a SNP in each parent (n = 136), the “patterning” and “never” categories were pooled. A contingency table was built counting orthologous pairs sharing each combination of first unbiased stages across the two species, and a one-sided Fisher's exact test was applied to each cell individually, comparing the observed count against the marginal totals of the full matrix. To test whether genes losing maternal bias at different developmental stages differed in their expression dynamics, we fitted a linear mixed model using the lme4 package^65^ in R, with p-values obtained using lmerTest^66^. Log-transformed expression (log1p TPM) was modelled as a function of developmental stage (coded numerically 1–6), gene group (the stage at which each gene first lost maternal bias, coded as an ordered factor from egg cell to never), their interaction, and cross direction (cross or reciprocal) as a covariate. Gene identity was included as a random intercept to account for the repeated-measures structure of the data. The ordered factor coding of gene groups allowed decomposition of group effects into polynomial contrasts, of which the linear component was the primary focus. To test whether genes in later-losing groups had globally higher expression levels, estimated marginal means were computed per group averaged across all developmental stages using the emmeans package (https://rvlenth.github.io/emmeans/) and a linear trend contrast was applied across the ordered groups. The monotonic shift in expression trajectories across groups was assessed via the linear × linear component of the stage × group interaction term.

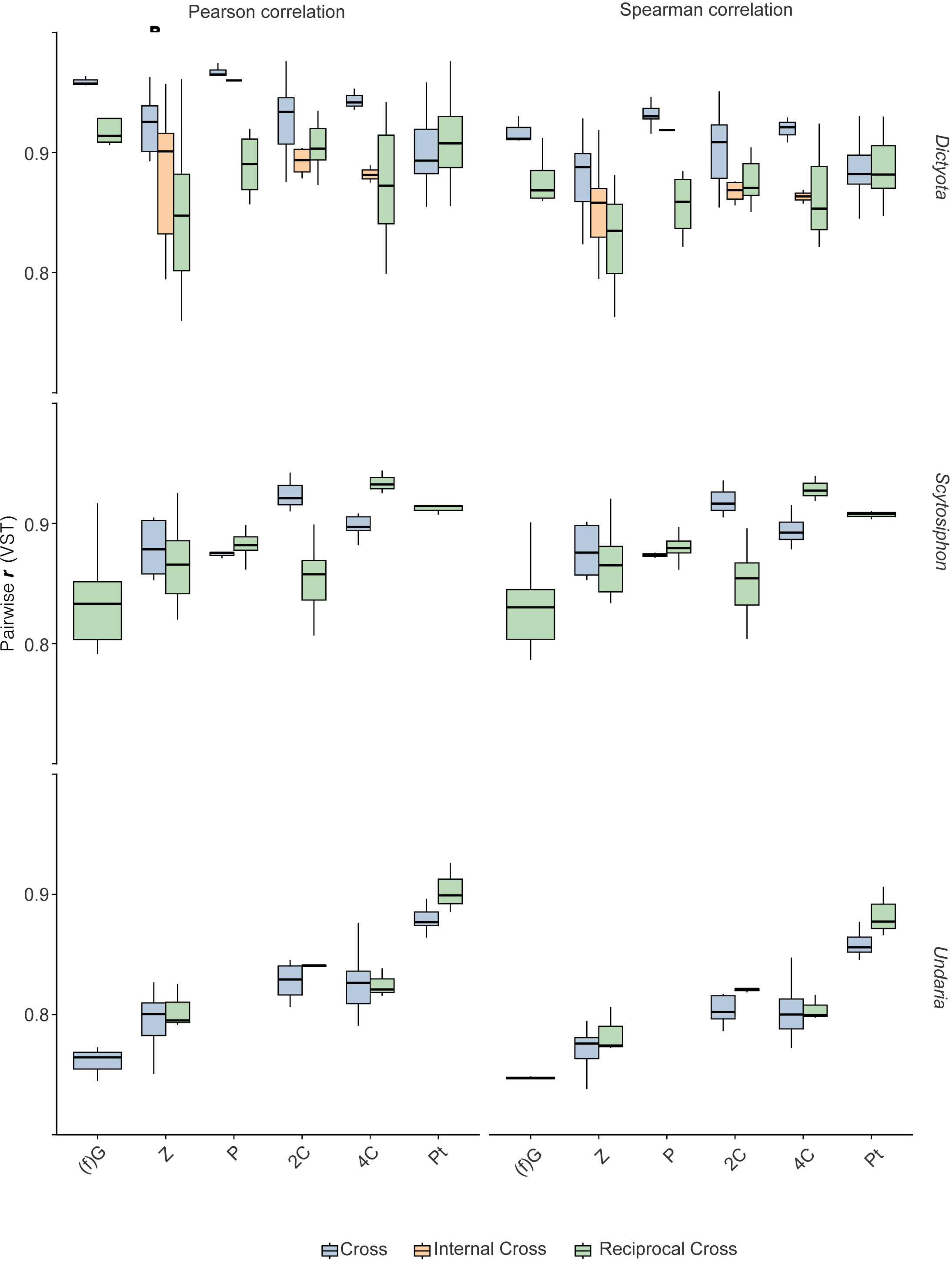

**Figure S1. Reproducibility of transcriptomes across biological replicates.** Pairwise correlations (Pearson and Spearman) of variance-stabilized (VST) gene expression values were computed between samples belonging to the same developmental stage and cross/genotype.

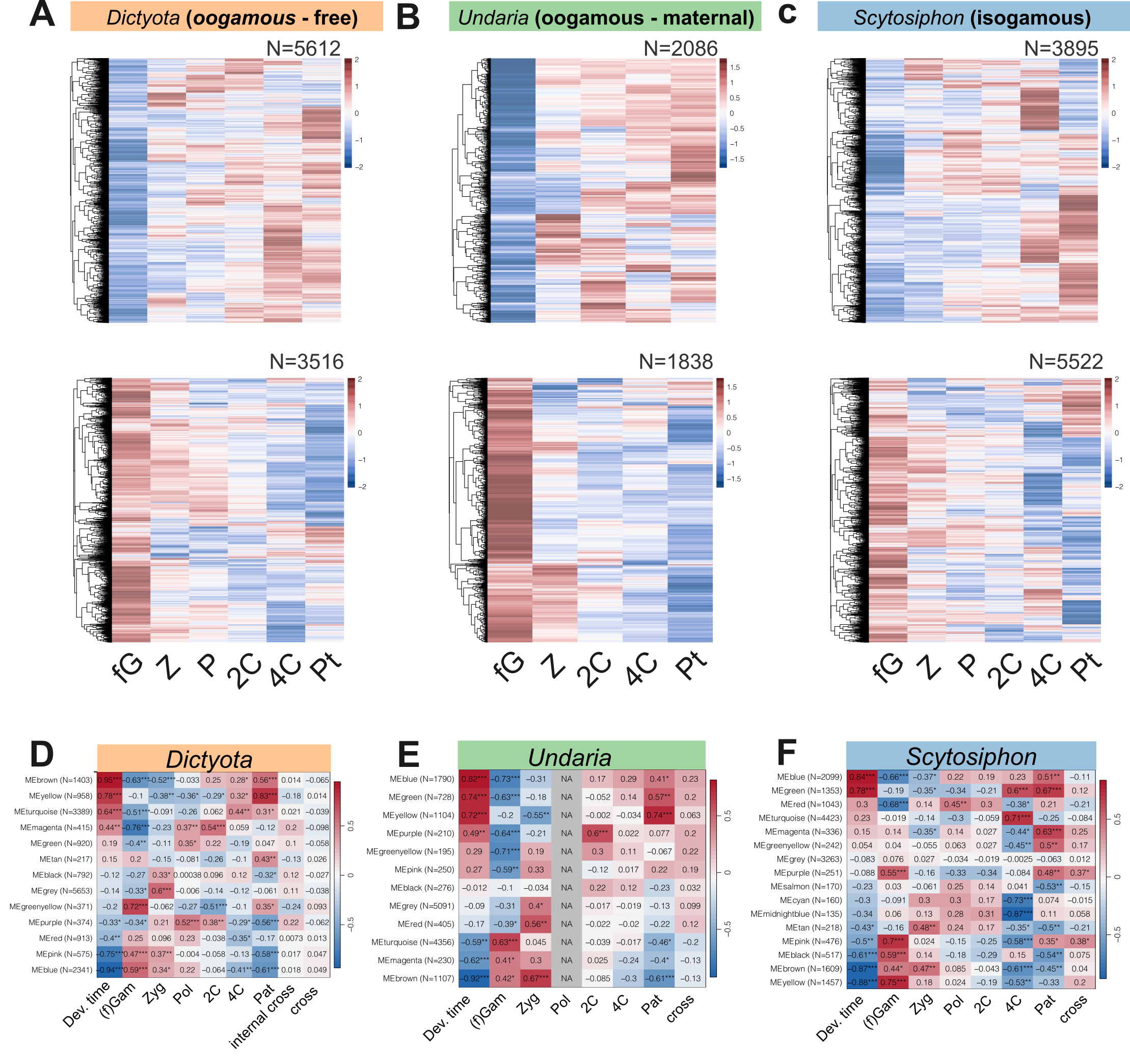

**Figure S2. Heatmaps of differentially expressed genes across early development in three brown algal species.** Scaled expression (z-score of VST-normalized counts) of differentially expressed genes (DEGs) identified by DESeq2 relative to female gametes (fG) across developmental stages: zygote (Z), polarised zygote (P), two-cell (2C), four-cell (4C), and patterning stage (Pt). Upper panels show upregulated DEGs and lower panels show downregulated DEGs for *Dictyota dichotoma* (oogamous, free-developing; **(A)**, *Undaria pinnatifida* (oogamous, maternally retained; **(B)**, and *Scytosiphon promiscuus* (isogamous; **(C)**. Number of differentially expressed genes per heatmap is indicated (N). Rows represent individual genes clustered by hierarchical clustering; columns represent developmental stages. Red indicates relatively high expression, blue indicates relatively low expression.(**D–F**) Module–trait correlation heatmaps for each species, showing Pearson correlation coefficients between WGCNA module eigengenes and developmental traits. Asterisks indicate statistical significance (* p<0.05, ** p<0.01, *** p<0.001).

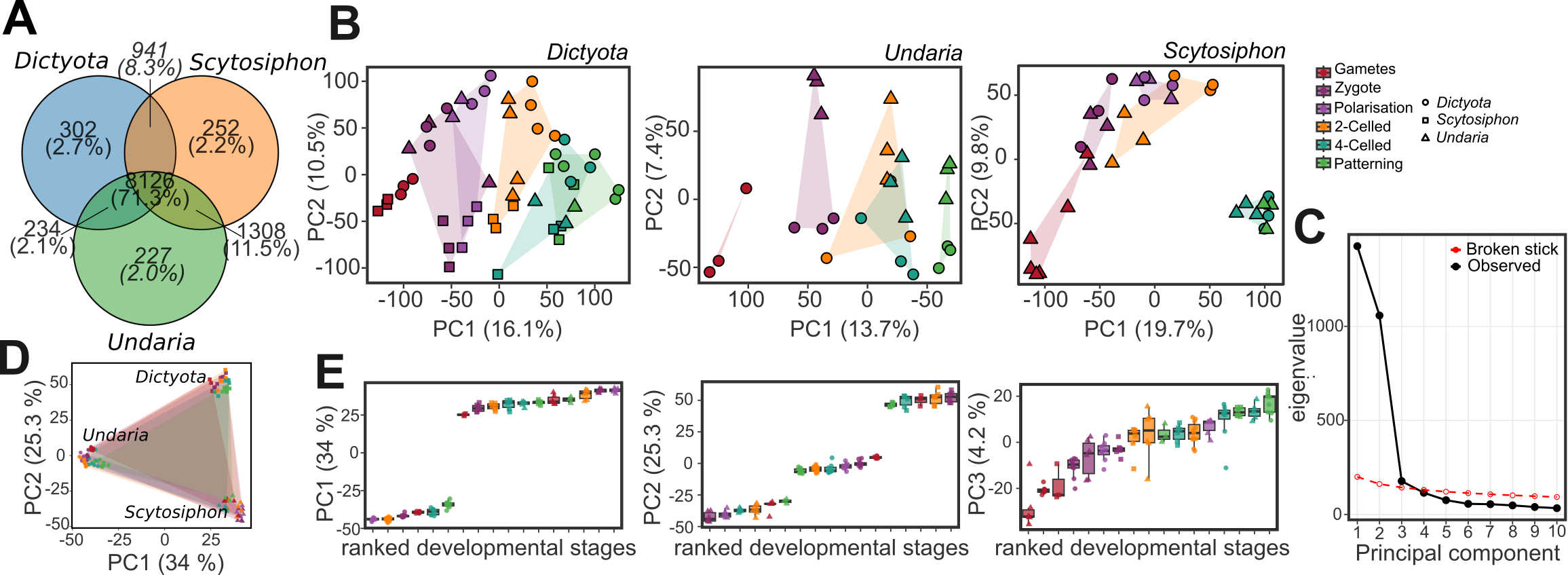

**Figure S3. Cross-species comparison of developmental transcriptomes using orthogroup expression.**  **(A)** Venn diagram showing the number of orthogroups shared among *Dictyota*, *Scytosiphon*, and *Undaria*, as identified by OrthoFinder. The 8,126 three-way orthogroups (71.3%) (single-copy orthologs) were used for all downstream analyses. **(B)** Per-species PCA on log_2_(TPM+1)-transformed expression of all single copy orthogroups, showing species-specific developmental trajectories across six stages (female gametes -> patterning). Points are coloured by developmental stage; convex hulls highlight stage clusters. **(C)** Scree plot of eigenvalues from the joint three-species PCA, compared against broken stick, indicating that at least three principal components capture non-random variance**. (D)** Joint three-species PCA (PC1 vs PC2) on log₂(TPM + 1)-transformed orthogroup expression. Samples cluster primarily by species rather than by developmental stage, with *Dictyota*, *Undaria*, and *Scytosiphon* occupying distinct regions of the PC1–PC2 space. Together with (B), this indicates that the two largest axes of variation in the joint dataset capture lineage-specific expression differences, whereas developmental progression conserved across species is captured by PC3 (see Figure 3c,d)**. (E)** Cross species PCA on log_2_(TPM+1) Boxplots showing the distribution of PC1, PC2, and PC3 sample scores across developmental stages of all organisms ranked on average principal component score, illustrating that PC3 captures developmental progression whereas PC1 and PC2 reflect species-specific variation.

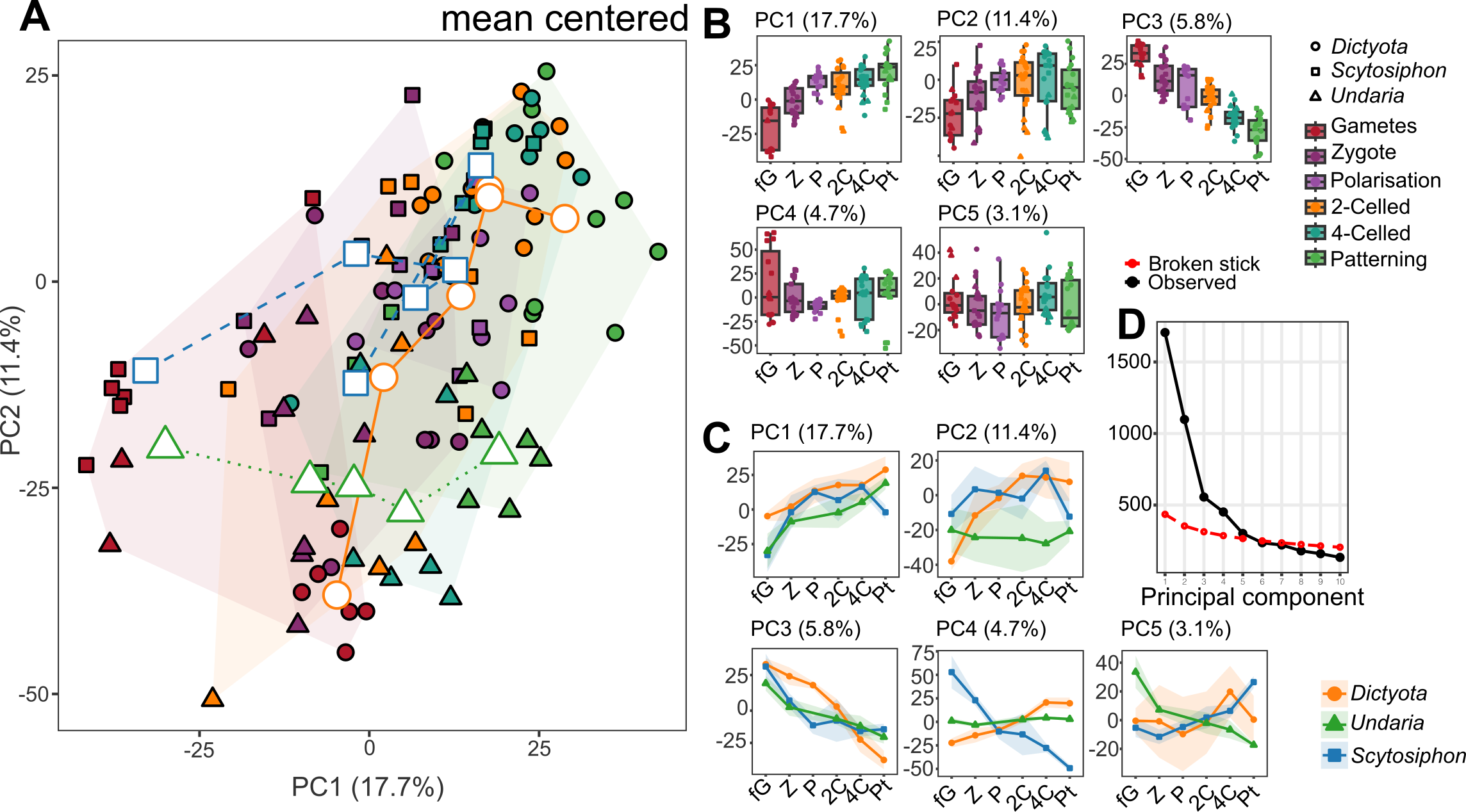

**Figure S4. Cross-species comparison of developmental transcriptomes using orthogroup expression and mean centered data.** **(A)** Joint PCA (PC1 vs PC2) of all three species based on mean-centered log₂(TPM + 1) orthogroup expression. Convex hulls are drawn per species; points are coloured by developmental stage and shaped by species. Mean-centering removes species-specific expression offsets, enhancing the visibility of shared developmental dynamics. **(B)** Boxplots showing the distribution of PC1–PC5 sample scores across developmental stages, illustrating that developmental progression is captured across multiple principal components when species-specific baselines are removed. **(C)** Line plots of mean PC scores across developmental stages for each species, showing species-specific trajectories along PC1–PC5, with correlated trajectories in PC1 and PC3. **(D)** Scree plot of eigenvalues from the mean-centered PCA, compared against broken stick, indicating that up to five principal components capture non-random variance according to the broken stick criterion.

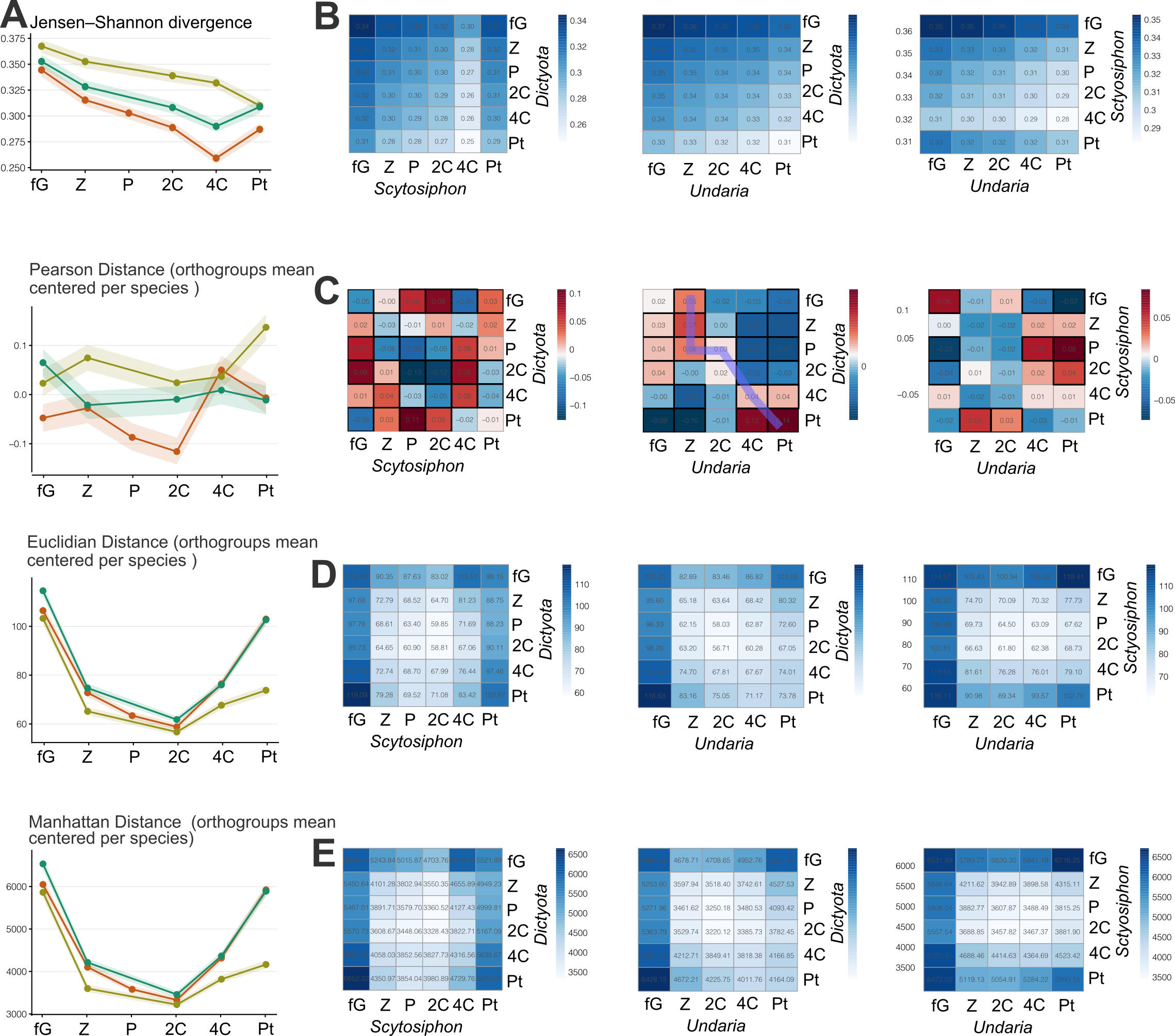

**Figure S5. Cross-species divergence of developmental transcriptomes across complementary distance metrics.** Pairwise transcriptome distances were computed between all developmental stages (female gametes, fG; zygote, Z; polarisation, P; 2-celled, 2C; 4-celled, 4C; patterning, Pt) across *Dictyota, Undaria and Scytosiphon,* using four complementary metrics applied to single-copy orthogroup expression. Each row shows one metric: line plots (left) summarise the distance between matched developmental stages across species pairs, while the three heatmaps (right) display the full pairwise distance matrix for each species pair (*Dictyota* vs *Scytosiphon*, *Dictyota* vs *Undaria*, *Scytosiphon* vs *Undaria*). Line colours denote species pairs and are consistent across all panels: orange = *Dictyota* vs *Scytosiphon*, olive = *Dictyota* vs *Undaria*, teal = *Scytosiphon* vs *Undaria*. Shaded ribbons denote bootstrap 95% confidence intervals. **(A)** Jensen–Shannon divergence computed on TPM-normalised orthogroup expression distributions, quantifying differences in the shape of the global expression distribution between stages. JSD decreases across development in all species pairs, indicating increasing similarity in global expression profiles of conserved orthogroups as embryogenesis progresses. **(B–E)** Pearson **(C)**, Euclidean **(D)**, and Manhattan **(E)** distances computed on log₂(TPM + 1)-transformed orthogroup expression after mean-centring per species to remove lineage-specific expression offsets. Mean-centring isolates the relative developmental dynamics of each orthogroup from its absolute expression baseline, allowing detection of conserved temporal trajectories independent of species-specific expression levels. All three metrics reveal a consistent mid-developmental convergence, with minimum inter-species distance around the 2C–4C stages and greater divergence at the earliest (fG, Z) and latest (Pt) stages.

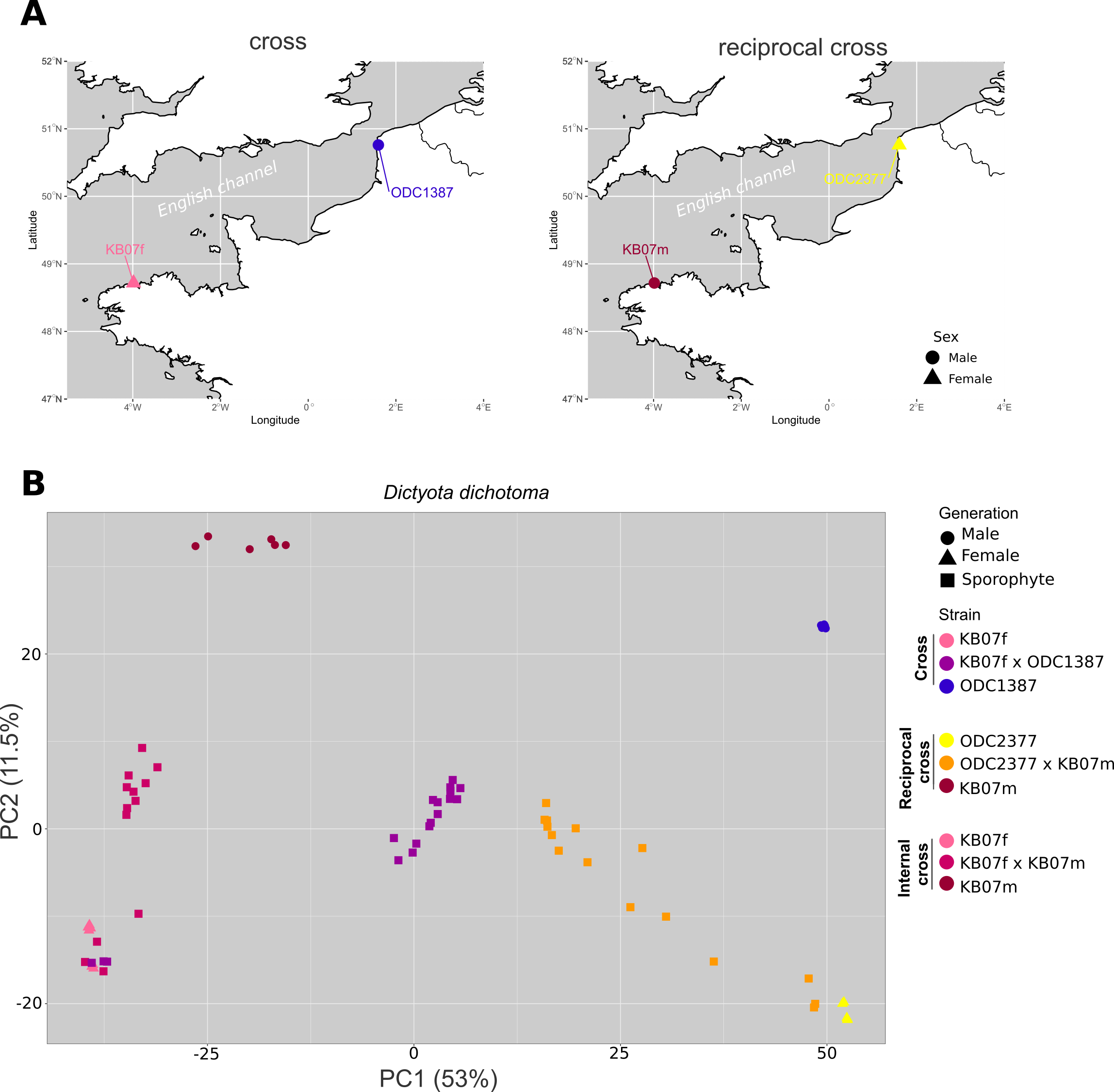

**Figure S6. Geographical origin and genetic identity of *Dictyota dichotoma* strains. (A)** Geographical origins of the strains used in the reciprocal crosses. Collection sites are indicated by points, with circles representing female strains and triangles representing male strains. Maps were generated in R using the rnaturalearth and sf packages. **(B)** Principal component analysis of SNP genotypes across all available samples, including haploid parental samples and diploid hybrid samples from the crosses. Each point represents a sample, coloured by strain and shaped by generation. The PCA was used to verify the genetic identity of parental strains prior to strain-specific SNP calling.

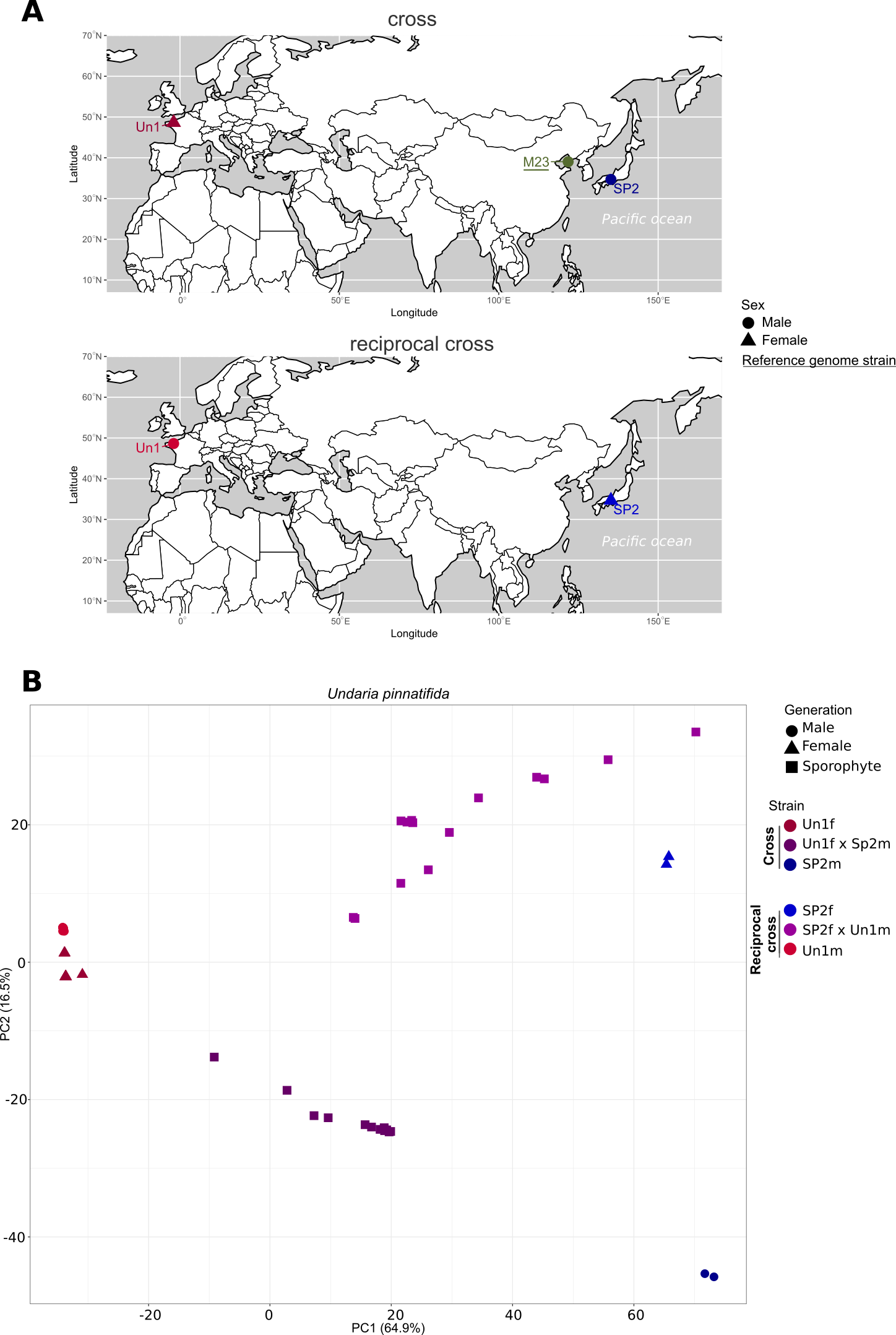

**Figure S7. Geographical origin and genetic identity of *Undaria pinnatifida* strains.** **(A)** Geographical origins of the strains used in the reciprocal crosses. Collection sites are indicated by points, with circles representing female strains and triangles representing male strains. Maps were generated in R using the rnaturalearth and sf packages. **(B)** Principal component analysis of SNP genotypes across all available samples, including haploid parental samples and diploid hybrid samples from the crosses. Each point represents a sample, coloured by strain and shaped by generation. The PCA was used to verify the genetic identity of parental strains prior to strain-specific SNP calling.

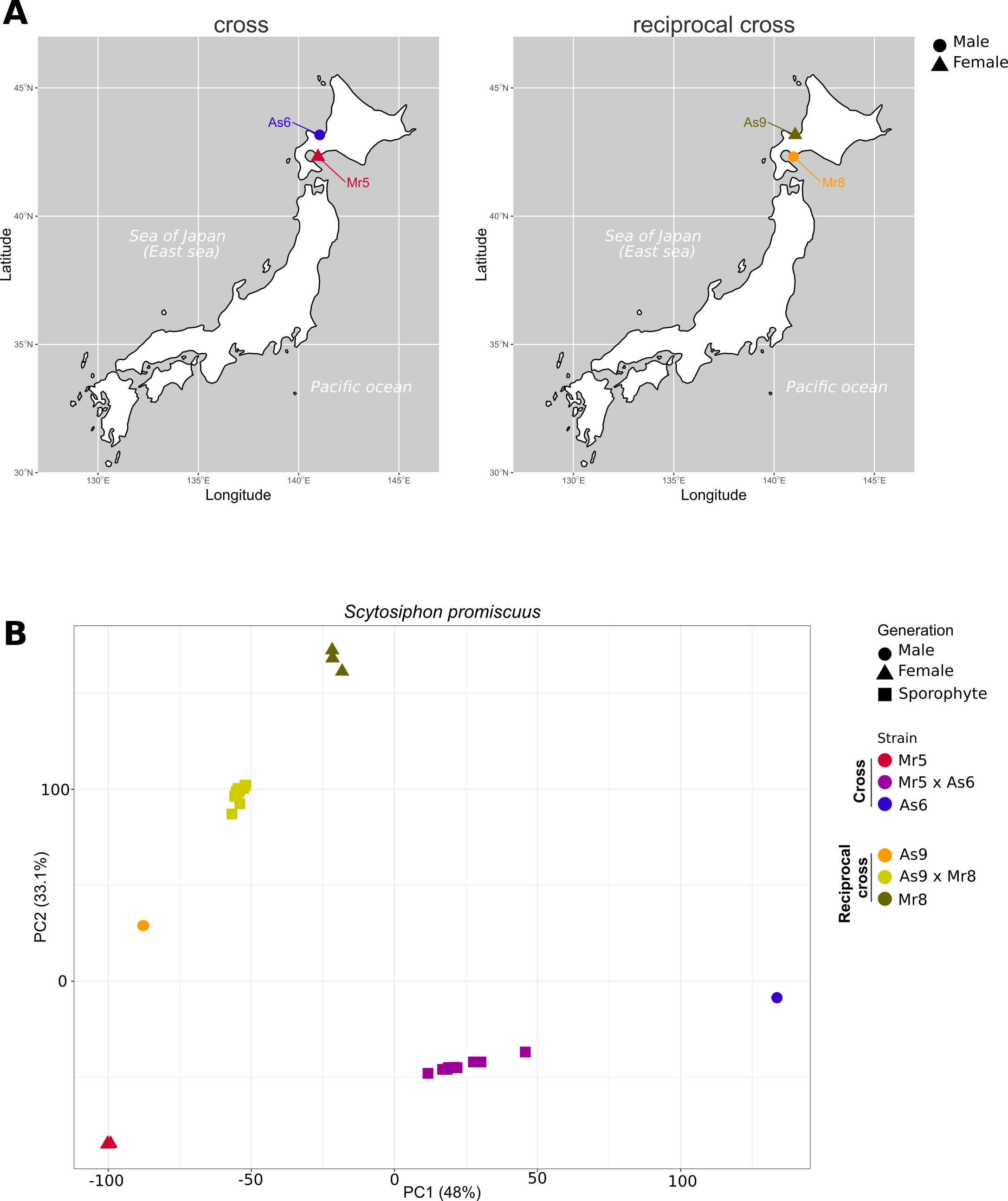

**Figure S8. Geographical origin and genetic identity of *Scytosiphon promiscuus* strains** **(A)** Geographical origins of the strains used in the reciprocal crosses. Collection sites are indicated by points, with circles representing female strains and triangles representing male strains. Maps were generated in R using the rnaturalearth and sf packages. **(B)** Principal component analysis of SNP genotypes across all available samples, including haploid parental samples and diploid hybrid samples from the crosses. Each point represents a sample, coloured by strain and shaped by generation. The PCA was used to verify the genetic identity of parental strains prior to strain-specific SNP calling.

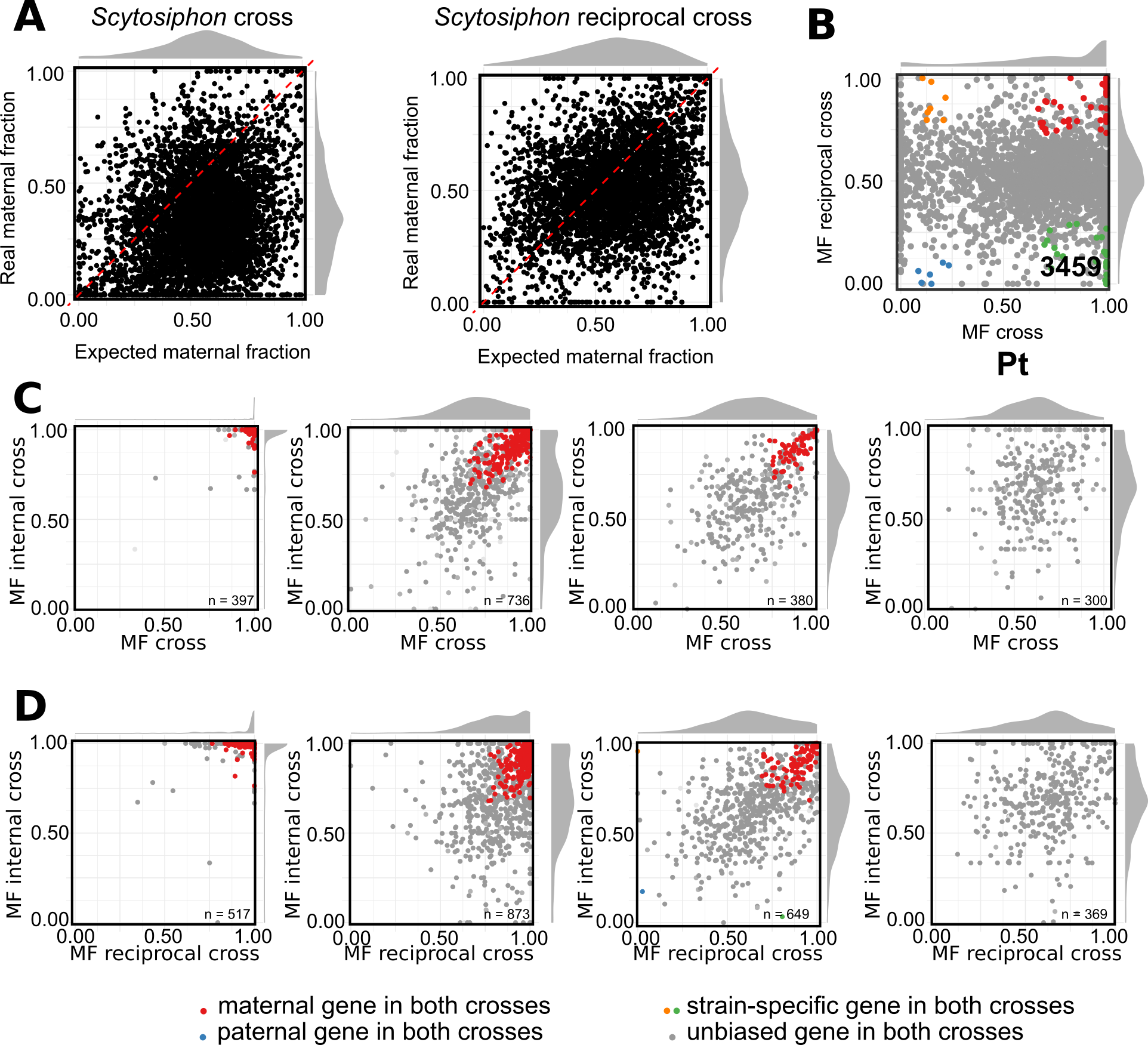

**Figure S9. Mean maternal fraction in the internal cross versus cross** s**hows that maternal bias dynamics are not driven by parental genetic divergence, with supporting *Scytosiphon* controls. (A)** Comparison of observed maternal fraction at the zygote stage with the maternal fraction expected under a dilution model assuming the male gamete contributes 80% of the volume of the female gamete, in *Scytosiphon promiscuus*. The diagonal represents the expectation under pure cytoplasmic dilution. **(B)** Mean maternal fraction in the cross versus reciprocal cross at the patterning (Pt) stage in *Scytosiphon*. The pronounced strain-specific clusters (orange, upper-left; green, lower-right) indicate that the cross sample was contaminated with female parthenotes, in which only the maternal-strain genome is present and transcripts are scored as "maternal" in the cross direction. This sample, harvested from a separate release period, had shown atypically slow development, raising prior suspicion of parthenote inclusion that the allelic pattern confirmed; it was therefore excluded from downstream analyses. **(C**) Internal cross versus reciprocal cross **(D)** for individual genes in *Dictyota dichotoma*, across developmental stages (n genes indicated per panel). Points are coloured by bias category: red, maternally biased in both crosses; blue, paternally biased in both crosses; yellow/green, strain-specific bias.

**Figure S10. GO enrichment and cross-species developmental stage comparison in brown algae.** **(A)** Comparison of first unbiased stage assignments between one-to-one orthologous gene pairs in *Dictyota* and *Undaria*. Cell colour indicates −log10(p-value) from a one-sided Fisher's exact test; significance levels: ** p < 0.01, *** p < 0.001. **(B-D)** Enriched GO terms for gene groups highlighted with asterisk in **Figure 4 (D and E)** in *Undaria* (**B**), and in *Dictyota* (**C** and **D**).

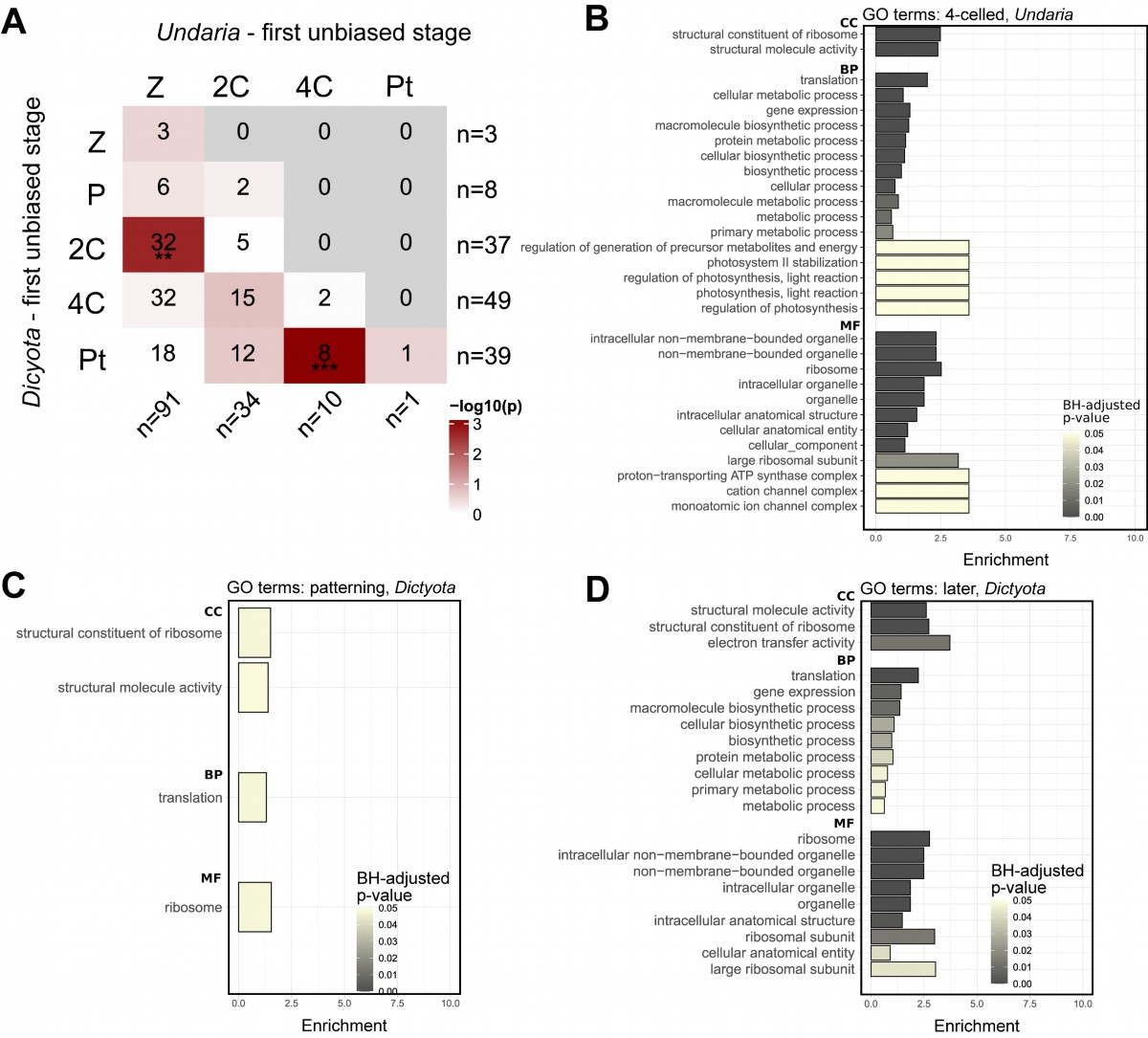

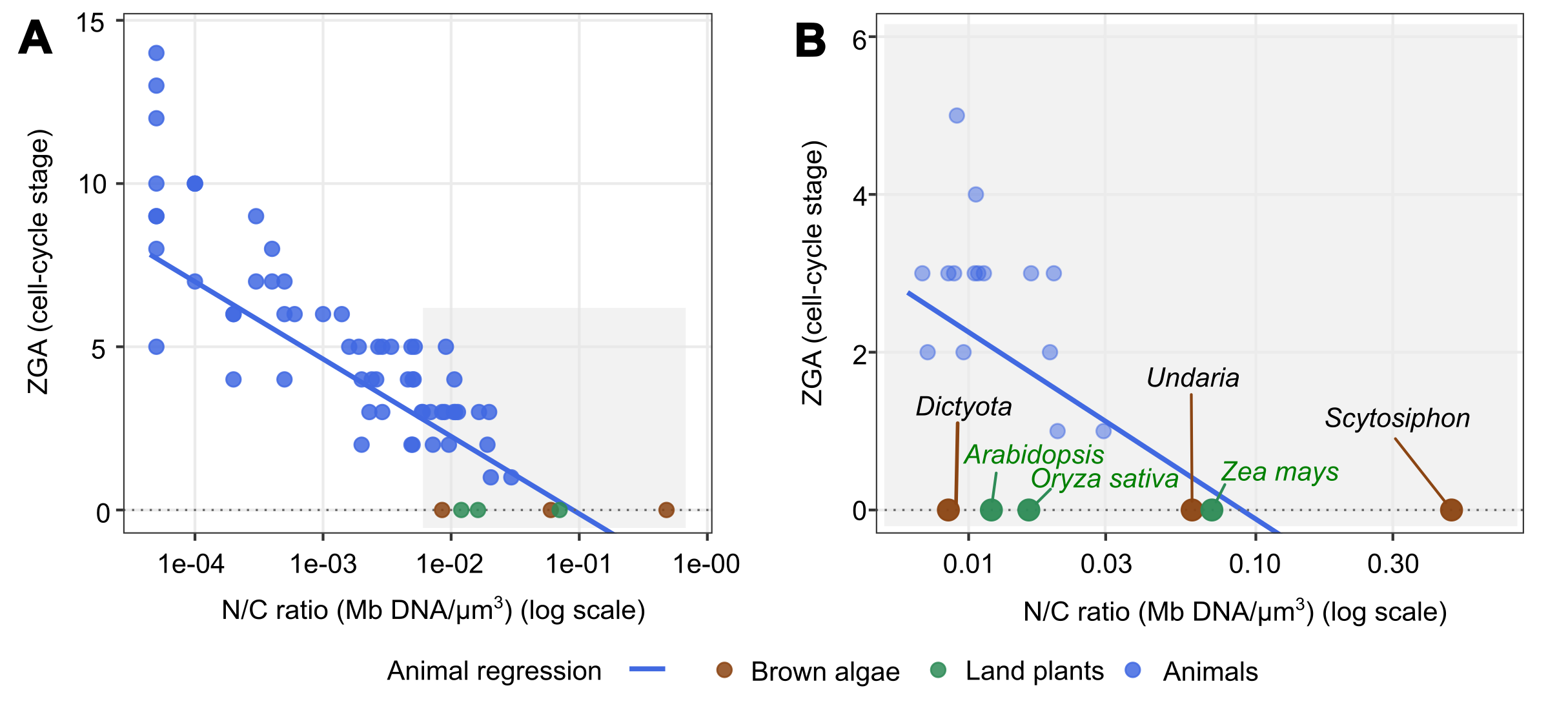

**Figure S11. Brown algae deviate from the animal nuclear-to-cytoplasmic (N/C) scaling of ZGA timing.** ZGA timing (cell-cycle stage) plotted against the N/C ratio (Mb DNA / µm³, log scale) for animals (blue), brown algae (brown) and land plants (green). The blue line shows the published animal regression, ZGA = -2.370 × log₁₀(N/C) + 11.734 (Campo-Bes et al., 2026). Animal ZGA values are taken from (Campo-Bes et al., 2026) see **Table S1**. **(A)** Full N/C range. The grey shaded box indicates the zoom region shown in (b). **(B)** Zoom into the N/C range occupied by non-animal species; animal points within this range are shown faded for context. All three brown algae (*Scytosiphon*, *Undaria*, *Dictyota*) and all three land plants (*Arabidopsis*, *Oryza sativa*, *Zea mays*) initiate ZGA at cell-cycle stage 0, i.e. before the first cleavage, regardless of their N/C ratio. An ANCOVA of ZGA against log₁₀(N/C) with clade as a factor rejects the hypothesis that non-animal species follow the animal slope (clade × log₁₀(N/C) interaction, F₂,₆₀ = 3.38, p = 0.040; full model adjusted R² = 0.78), with the brown algae interaction term significant individually (p = 0.021) and the plants term non-significant (p = 0.27).

**Table S1. Nuclear-to-cytoplasmic (N/C) ratios and zygotic genome activation (ZGA) timing used for comparative analysis** (see Figure 1). N/C ratio = haploid genome size (Mb) / zygote cytoplasmic volume (µm³). For animal species, haploid genome sizes, ZGA timings and egg volumes were derived from Campo-Bes et al. (2026) and their summarized sources. ZGA timing is expressed as the cell cycle number at which ZGA initiates, as reported in the Source ZGA column; predicted ZGA values for animals are from Campo-Bes et al. (2026; https://doi.org/10.64898/2026.04.16.718233). For the isogamous *Scytosiphon promiscuus*, the zygote volume was estimated as twice the gamete volume. Predicted ZGA values reported are based on diploid genome content (2× haploid), following the formula of Campo-Bes et al. (2026): y = −2.370 × log₁₀(X*106) + 11.734, with the X = N/C ratio.

| **Species** | **Clade** | **Haploid genome (Mb)** | **Genome source** | **Volume (µm³)** | **Volume source** | **N/C ratio (Mb DNA µm⁻³)** | **ZGA (cycles)** | **Source ZGA** | **predicted ZGA (Campo-Bes et al. 2026 BioRxiv)** |
| --- | --- | --- | --- | --- | --- | --- | --- | --- | --- |
| ***Scytosiphon promiscuus*** | Brown algae | 193 | Barrera-Redondo et al. (2025) | 804 | Hoshino, M., & Kogame, K. (2019). Journal of phycology, 55(2), 466-472. | 0.4801 | 0 | *this study* | -1.73 |
| ***Undaria pinnatifida*** | Brown algae | 511 | Shan et al. (2020) | 17,031 | Dries, E., Meyers, Y., Liesner, D., Gonzaga, F. M., Becker, J. F., Zakka, E. E., Beeckman, T., Coelho, S.M., De Clerck, O., & Bogaert, K. A. (2024). New Phytologist, 243(5), 1887-1898. | 0.06 | 0 | *this study* | 0.41 |
| ***Dictyota dichotoma*** | Brown algae | 854 | Barrera-Redondo et al. (2025) | 201,491 | Bogaert, K. A., Beeckman, T., & De Clerck, O. (2017). Annals of Botany, 120(4), 529-538.; | 0.0085 | 0 | *this study* | 2.42 |
|  |  |  |  |  | Manton, I. (1959). Journal of Experimental Botany, 10(3), 448-461. |  |  |  |  |
| ***Arabidopsis thaliana*** | Land plants | 119.1 | <https://www.ncbi.nlm.nih.gov/datasets/genome/GCF_000001735.4/> | 19,900 | Mansfield, S. G., & Briarty, L. G. (1991). The developing embryo. Canadian Journal of Botany, 69(3), 461-476. | 0.012 | 0 | <https://doi.org/10.1038/nature10756;https://doi.org/10.1016/j.devcel.2019.04.016;https://doi.org/10.1242/dev.204449> | 2.07 |
| ***Oryza sativa (rice)*** | Land plants | 385.7 | <https://www.ncbi.nlm.nih.gov/datasets/genome/GCF_001433935.1/> | 47,713 | Uchiumi, T., Uemura, I. & Okamoto, T. Establishment of an in vitro fertilization system in rice (Oryza sativa L.). Planta 226, 581-589 (2007). https://doi.org/10.1007/s00425-007-0506-2 | 0.0162 | 0 | <https://doi.org/10.1242/dev.204497> | 1.76 |
| ***Zea mays*** | Land plants | 2200 | <https://www.ncbi.nlm.nih.gov/datasets/genome/GCF_902167145.1/> | 62,628 | Faure, J. E., Mogensen, H. L., Kranz, E., Digonnet, C., & Dumas, C. (1992). Protoplasma, 171(3), 97-103. | 0.0703 | 0 | <https://doi.org/10.1105/tpc.17.00099> | 0.25 |
| ***Homo sapiens*** | Animals | 3,100 | https://doi.org/10.64898/2026.04.16.718233 | 696,910 | <https://doi.org/10.1007/s10815-022-02602-0;https://doi.org/10.1007/s10815-011-9555-3;https://doi.org/10.1055/a-1876-2231> | 0.0089 | 3 | <https://doi.org/10.1007/s10815-022-02602-0> | 2.37 |
| ***Macaca mulatta*** | Animals | 2,971 | https://doi.org/10.64898/2026.04.16.718233 | 523,599 | <https://doi.org/10.1038/srep06598;> | 0.0113 | 3 | <https://doi.org/10.1038/srep06598> | 2.12 |
| ***Callithrix jacchus*** | Animals | 2,830 | https://doi.org/10.64898/2026.04.16.718233 | 523,599 | <https://doi.org/10.1186/s13048-024-01441-0;https://doi.org/10.1038/s41598-023-45224-x;> | 0.0108 | 3 | <https://doi.org/10.1186/s13048-024-01441-0> | 2.17 |
| ***Mus musculus*** | Animals | 2,728 | https://doi.org/10.64898/2026.04.16.718233 | 268,083 | <https://doi.org/10.1262/jrd.2022-101;https://doi.org/10.1242/dev.039487;> | 0.0204 | 1 | <https://doi.org/10.1262/jrd.2022-101> | 1.52 |
| ***Rattus norvegicus*** | Animals | 2,648 | https://doi.org/10.64898/2026.04.16.718233 | 179,594 | <https://doi.org/10.1126/science.1088313;> | 0.0295 | 1 | <https://doi.org/10.1126/science.1088313> | 1.14 |
| ***Oryctolagus cuniculus*** | Animals | 2,738 | https://doi.org/10.64898/2026.04.16.718233 | 523,599 | <https://doi.org/10.1038/258719a0;> | 0.0105 | 3 | <https://doi.org/10.1038/258719a0> | 2.21 |
| ***Bos taurus*** | Animals | 2,711 | https://doi.org/10.64898/2026.04.16.718233 | 904,779 | <https://doi.org/10.1016/S0093-691X(97)00300-2;https://doi.org/10.1002/mrd.20639;> | 0.006 | 3 | <https://doi.org/10.1016/S0093-691X(97)00300-2> | 2.78 |
| ***Bos grunniens*** | Animals | 2,898 | https://doi.org/10.64898/2026.04.16.718233 | 1,150,347 | <https://doi.org/10.6000/1927-520X.2020.09.16;> | 0.005 | 4 | <https://doi.org/10.6000/1927-520X.2020.09.16> | 2.96 |
| ***Bubalus bubalis*** | Animals | 2,923 | https://doi.org/10.64898/2026.04.16.718233 | 1,150,347 | <https://doi.org/10.3389/fgene.2019.01040;> | 0.0051 | 4 | <https://doi.org/10.3389/fgene.2019.01040> | 2.95 |
| ***Ovis aries*** | Animals | 2,619 | https://doi.org/10.64898/2026.04.16.718233 | 1,150,347 | <https://doi.org/10.3390/ani14050807;> | 0.0046 | 4 | <https://doi.org/10.3390/ani14050807> | 3.06 |
| ***Sus scrofa*** | Animals | 2,502 | https://doi.org/10.64898/2026.04.16.718233 | 696,910 | <https://doi.org/10.3906/vet-1502-16;> | 0.0072 | 2 | <https://doi.org/10.3906/vet-1502-16> | 2.59 |
| ***Equus caballus*** | Animals | 2,507 | https://doi.org/10.64898/2026.04.16.718233 | 1,022,654 | <https://doi.org/10.1093/biolre/ioac172;> | 0.0049 | 2 | <https://doi.org/10.1093/biolre/ioac172> | 2.99 |
| ***Monodelphis domestica*** | Animals | 3,601 | https://doi.org/10.64898/2026.04.16.718233 | 1,441,991 | <https://doi.org/10.1038/s41586-025-08992-2;> | 0.005 | 2 | <https://doi.org/10.1038/s41586-025-08992-2> | 2.97 |
| ***Gallus gallus*** | Animals | 1,053 | https://doi.org/10.64898/2026.04.16.718233 | 418,879,021 | <https://doi.org/10.1371/journal.pone.0080631;> | 0 | 13 | <https://doi.org/10.1371/journal.pone.0080631> | 10.07 |
| ***Xenopus tropicalis*** | Animals | 1,451 | https://doi.org/10.64898/2026.04.16.718233 | 220,893,234 | <https://doi.org/10.1002/dvdy.10178;> | 0 | 9 | <https://doi.org/10.1002/dvdy.10178> | 9.08 |
| ***Ambystoma mexicanum*** | Animals | 29,100 | https://doi.org/10.64898/2026.04.16.718233 | 4,188,790,205 | <https://doi.org/10.1016/j.ydbio.2016.05.024;> | 0 | 9 | <https://doi.org/10.1016/j.ydbio.2016.05.024> | 9.03 |
| ***Danio rerio*** | Animals | 1,373 | https://doi.org/10.64898/2026.04.16.718233 | 22,061,834 | https://doi.org/10.64898/2026.04.16.718233 | 0.0001 | 10 | https://doi.org/10.64898/2026.04.16.718233 | 6.77 |
| ***Oryzias latipes*** | Animals | 734 | https://doi.org/10.64898/2026.04.16.718233 | 14,980,685 | https://doi.org/10.64898/2026.04.16.718233 | 0.0001 | 10 | https://doi.org/10.64898/2026.04.16.718233 | 7.01 |
| ***Lethenteron camtschaticum*** | Animals | 1,063 | https://doi.org/10.64898/2026.04.16.718233 | 150,796,447 | https://doi.org/10.64898/2026.04.16.718233 | 0 | 8 | https://doi.org/10.64898/2026.04.16.718233 | 9.01 |
| ***Ciona savignyi*** | Animals | 177 | https://doi.org/10.64898/2026.04.16.718233 | 606,131 | [https://doi.org/10.1038/s41467-024-46780-0;doi: 10.3391/ai.2010.5.4.05;](https://doi.org/10.1038/s41467-024-46780-0;doi:%2010.3391/ai.2010.5.4.05;) | 0.0006 | 6 | <https://doi.org/10.1038/s41467-024-46780-0> | 5.18 |
| ***Phallusia mammillata*** | Animals | 234 | https://doi.org/10.64898/2026.04.16.718233 | 904,779 | <https://doi.org/10.1016/j.cub.2008.05.039;> | 0.0005 | 4 | <https://doi.org/10.1016/j.cub.2008.05.039> | 5.3 |
| ***Oikopleura dioica*** | Animals | 64 | https://doi.org/10.64898/2026.04.16.718233 | 268,083 | <https://doi.org/10.1111/dgd.12769;> | 0.0005 | 7 | <https://doi.org/10.1111/dgd.12769> | 5.38 |
| ***Branchiostoma floridae*** | Animals | 513 | https://doi.org/10.64898/2026.04.16.718233 | 2,806,162 | <https://doi.org/10.3389/fcell.2021.668006;> | 0.0004 | 7 | <https://doi.org/10.3389/fcell.2021.668006> | 5.66 |
| ***Branchiostoma lanceolatum*** | Animals | 475 | https://doi.org/10.64898/2026.04.16.718233 | 904,779 | <https://doi.org/10.3389/fcell.2021.668006;> | 0.001 | 6 | <https://doi.org/10.3389/fcell.2021.668006> | 4.57 |
| ***Strongylocentrotus purpuratus*** | Animals | 922 | https://doi.org/10.64898/2026.04.16.718233 | 268,083 | https://doi.org/10.64898/2026.04.16.718233 | 0.0069 | 3 | https://doi.org/10.64898/2026.04.16.718233 | 2.64 |
| ***Paracentrotus lividus*** | Animals | 928 | https://doi.org/10.64898/2026.04.16.718233 | 381,704 | https://doi.org/10.64898/2026.04.16.718233 | 0.0049 | 5 | https://doi.org/10.64898/2026.04.16.718233 | 3 |
| ***Lytechinus pictus*** | Animals | 999 | https://doi.org/10.64898/2026.04.16.718233 | 696,910 | <https://doi.org/10.1002/dvdy.223;> | 0.0029 | 5 | <https://doi.org/10.1002/dvdy.223> | 3.54 |
| ***Holothuria leucospilota*** | Animals | 1,391 | https://doi.org/10.64898/2026.04.16.718233 | 1,436,755 | <https://doi.org/10.1016/j.aquaculture.2018.01.013;> | 0.0019 | 5 | <https://doi.org/10.1016/j.aquaculture.2018.01.013> | 3.94 |
| ***Patiria miniata*** | Animals | 608 | https://doi.org/10.64898/2026.04.16.718233 | 3,053,628 | <https://doi.org/10.1101/2025.03.24.644836;> | 0.0004 | 8 | <https://doi.org/10.1101/2025.03.24.644836> | 5.57 |
| ***Ptychodera flava*** | Animals | 1,162 | https://doi.org/10.64898/2026.04.16.718233 | 904,779 | <https://doi.org/10.2108/zsj.15.85;> | 0.0026 | 4 | <https://doi.org/10.2108/zsj.15.85> | 3.65 |
| ***Symsagittifera roscoffensis*** | Animals | 754 | https://doi.org/10.64898/2026.04.16.718233 | 523,599 | https://doi.org/10.64898/2026.04.16.718233 | 0.0029 | 3 | https://doi.org/10.64898/2026.04.16.718233 | 3.54 |
| ***Caenorhabditis elegans*** | Animals | 100 | https://doi.org/10.64898/2026.04.16.718233 | 23,562 | <https://www.ncbi.nlm.nih.gov/books/NBK20121/;> | 0.0085 | 3 | <https://www.ncbi.nlm.nih.gov/books/NBK20121/> | 2.42 |
| ***Caenorhabditis angaria*** | Animals | 107 | https://doi.org/10.64898/2026.04.16.718233 | 23,562 | <https://doi.org/10.1242/dev.200984;> | 0.0091 | 5 | <https://doi.org/10.1242/dev.200984> | 2.35 |
| ***Caenorhabditis plicata*** | Animals | 132 | https://doi.org/10.64898/2026.04.16.718233 | 50,265 | https://doi.org/10.64898/2026.04.16.718233 | 0.0052 | 5 | https://doi.org/10.64898/2026.04.16.718233 | 2.92 |
| ***Oscheius tipulae*** | Animals | 61 | https://doi.org/10.64898/2026.04.16.718233 | 11,454 | <https://doi.org/10.1016/j.cub.2022.10.043;> | 0.0106 | 4 | <https://doi.org/10.1016/j.cub.2022.10.043> | 2.19 |
| ***Steinernema carpocapsae*** | Animals | 85 | https://doi.org/10.64898/2026.04.16.718233 | 28,731 | <https://doi.org/10.1093/gbe/evx195;> | 0.0059 | 3 | <https://doi.org/10.1093/gbe/evx195> | 2.8 |
| ***Steinernema feltiae*** | Animals | 122 | https://doi.org/10.64898/2026.04.16.718233 | 14,726 | <https://doi.org/10.1093/gbe/evx195;> | 0.0165 | 3 | <https://doi.org/10.1093/gbe/evx195> | 1.74 |
| ***Ditylenchus destructor*** | Animals | 139 | https://doi.org/10.64898/2026.04.16.718233 | 137,445 | <https://doi.org/10.1038/s41597-024-03542-3;> | 0.002 | 2 | <https://doi.org/10.1038/s41597-024-03542-3> | 3.9 |
| ***Heterodera glycines*** | Animals | 158 | https://doi.org/10.64898/2026.04.16.718233 | 137,445 | <https://doi.org/10.1038/s41597-024-03542-3;> | 0.0023 | 3 | <https://doi.org/10.1038/s41597-024-03542-3> | 3.77 |
| ***Meloidogyne incognita*** | Animals | 184 | https://doi.org/10.64898/2026.04.16.718233 | 137,445 | <https://doi.org/10.1038/s41597-024-03542-3;> | 0.0027 | 5 | <https://doi.org/10.1038/s41597-024-03542-3> | 3.61 |
| ***Ascaris suum*** | Animals | 298 | https://doi.org/10.64898/2026.04.16.718233 | 31,059 | <http://dx.doi.org/10.3347/kjp.2012.50.3.239;> | 0.0192 | 2 | <http://dx.doi.org/10.3347/kjp.2012.50.3.239> | 1.58 |
| ***Drosophila melanogaster*** | Animals | 144 | https://doi.org/10.64898/2026.04.16.718233 | 12,300,000 | <https://doi.org/10.2307/4099;https://doi.org/10.1111/j.1420-9101.2008.01649.x;> | 0 | 14 | <https://doi.org/10.2307/4099> | 8.49 |
| ***Tribolium castaneum*** | Animals | 166 | https://doi.org/10.64898/2026.04.16.718233 | 28,274,334 | <https://doi.org/10.1016/j.cois.2016.08.002;> | 0 | 12 | <https://doi.org/10.1016/j.cois.2016.08.002> | 9.2 |
| ***Blattella germanica*** | Animals | 1,781 | https://doi.org/10.64898/2026.04.16.718233 | 589,048,623 | <https://doi.org/10.1186/s13227-024-00234-2;> | 0 | 10 | <https://doi.org/10.1186/s13227-024-00234-2> | 9.88 |
| ***Cloeon dipterum*** | Animals | 180 | https://doi.org/10.64898/2026.04.16.718233 | 3,046,690 | https://doi.org/10.64898/2026.04.16.718233 | 0.0001 | 7 | https://doi.org/10.64898/2026.04.16.718233 | 6.82 |
| ***Parhyale hawaiensis*** | Animals | 2,757 | https://doi.org/10.64898/2026.04.16.718233 | 19,293,568 | <https://doi.org/10.1242/dev.00155;> | 0.0003 | 7 | <https://doi.org/10.1242/dev.00155> | 5.91 |
| ***Parasteatoda tepidariorum*** | Animals | 1,230 | https://doi.org/10.64898/2026.04.16.718233 | 65,449,847 | <https://doi.org/10.1186/s12915-022-01421-0;https://doi.org/10.1007/s00427-019-00631-x;https://doi.org/10.1242/dev.078204> | 0 | 5 | <https://doi.org/10.1186/s12915-022-01421-0> | 8 |
| ***Hypsibius exemplaris*** | Animals | 104 | https://doi.org/10.64898/2026.04.16.718233 | 87,114 | https://doi.org/10.64898/2026.04.16.718233 | 0.0024 | 4 | https://doi.org/10.64898/2026.04.16.718233 | 3.73 |
| ***Owenia fusiformis*** | Animals | 519 | https://doi.org/10.64898/2026.04.16.718233 | 523,599 | <https://doi.org/10.1017/S0016756803278341;> | 0.002 | 4 | <https://doi.org/10.1017/S0016756803278341> | 3.92 |
| ***Platynereis dumerilii*** | Animals | 1,470 | https://doi.org/10.64898/2026.04.16.718233 | 2,144,661 | <https://doi.org/10.1371/journal.pone.0295290;> | 0.0014 | 6 | <https://doi.org/10.1371/journal.pone.0295290> | 4.3 |
| ***Capitella teleta*** | Animals | 334 | https://doi.org/10.64898/2026.04.16.718233 | 4,188,790 | https://doi.org/10.64898/2026.04.16.718233 | 0.0002 | 4 | https://doi.org/10.64898/2026.04.16.718233 | 6.51 |
| ***Urechis unicinctus*** | Animals | 1,195 | https://doi.org/10.64898/2026.04.16.718233 | 696,910 | <https://doi.org/10.3390/ijms21072306;> | 0.0034 | 5 | <https://doi.org/10.3390/ijms21072306> | 3.36 |
| ***Crassostrea gigas*** | Animals | 648 | https://doi.org/10.64898/2026.04.16.718233 | 65,450 | https://doi.org/10.64898/2026.04.16.718233 | 0.0198 | 3 | https://doi.org/10.64898/2026.04.16.718233 | 1.55 |
| ***Mytilus galloprovincialis*** | Animals | 1,282 | https://doi.org/10.64898/2026.04.16.718233 | 268,083 | https://doi.org/10.64898/2026.04.16.718233 | 0.0096 | 2 | https://doi.org/10.64898/2026.04.16.718233 | 2.3 |
| ***Lingula anatina*** | Animals | 406 | https://doi.org/10.64898/2026.04.16.718233 | 523,599 | <https://doi.org/10.1038/ncomms9301;> | 0.0016 | 5 | <https://doi.org/10.1038/ncomms9301> | 4.17 |
| ***Brachionus plicatilis*** | Animals | 109 | https://doi.org/10.64898/2026.04.16.718233 | 448,921 | <https://doi.org/10.1371/journal.pone.0029365;> | 0.0005 | 6 | <https://doi.org/10.1371/journal.pone.0029365> | 5.37 |
| ***Nematostella vectensis*** | Animals | 273 | https://doi.org/10.64898/2026.04.16.718233 | 6,370,626 | <https://doi.org/10.1016/j.ydbio.2007.07.029;> | 0.0001 | 10 | <https://doi.org/10.1016/j.ydbio.2007.07.029> | 7.15 |
| ***Hydractinia symbiolongicarpus*** | Animals | 481 | https://doi.org/10.64898/2026.04.16.718233 | 4,849,048 | <https://doi.org/10.15252/embj.2022112934;> | 0.0002 | 6 | <https://doi.org/10.15252/embj.2022112934> | 6.29 |
| ***Clytia hemisphaerica*** | Animals | 421 | https://doi.org/10.64898/2026.04.16.718233 | 3,053,628 | https://doi.org/10.64898/2026.04.16.718233 | 0.0003 | 9 | https://doi.org/10.64898/2026.04.16.718233 | 5.95 |
| ***Mnemiopsis leidyi*** | Animals | 208 | https://doi.org/10.64898/2026.04.16.718233 | 1,949,816 | <https://doi.org/10.1038/s41596-022-00702-w;> | 0.0002 | 6 | <https://doi.org/10.1038/s41596-022-00702-w> | 6.21 |

**Table S2.** **Biological strains and genomes used in this study**. *Left*: strains sampled for RNA-seq, with species, strain identifier, geographical origin, and sex of the initially collected individual. *Right*: reference genomes used for read mapping, with the corresponding species, strain, publication, and DOI.

| **Samples** | | | | **Genomes** | | | |
| --- | --- | --- | --- | --- | --- | --- | --- |
| **Species** | **Strain** | **Geographical origin** | **Sex of the initially collected individual** | **species** | **strain** | **publication** | **DOI** |
| ***Dictyota dichotoma*** | ODC2377 | Wimereux, France | sporophyte | *Dictyota dichotoma* | KB07 | Barrera-Redondo et al. 2025 | 10.1038/s41559-025-02838-w |
| ***Dictyota dichotoma*** | ODC1387 | Wimereux, France | sporophyte | *Undaria pinnatifida* | M23 | Shan et al. 2020 | 10.3389/fgene.2020.00140 |
| ***Dictyota dichotoma*** | KB07V | Roscoff, France | sporophyte | *Scytosiphon promiscuus* | As6 | Barrera-Redondo et al. 2025 | 10.1038/s41559-025-02838-w |
| ***Undaria pinnatifida*** | UN1 (Phaeo216 and 218) | Saint-Malo, France | sporophyte |  |  |  |  |
| ***Undaria pinnatifida*** | UpinK2 | Kobe, Japan | sporophyte |  |  |  |  |
| ***Scytosiphon promiscuus*** | As6 | Asari, Japan | male |  |  |  |  |
| ***Scytosiphon promiscuus*** | As9 | Asari, Japan | female |  |  |  |  |
| ***Scytosiphon promiscuus*** | Mr5 (sxs103) | Muroran, Japan | female |  |  |  |  |
| ***Scytosiphon promiscuus*** | Mr8 | Muroran, Japan | male |  |  |  |  |

**Table S3. RNA-seq samples generated for the developmental time course.** For each library, the species, cross type (direct, reciprocal, or internal), parental strains, life cycle stage, developmental stage, time point (hours or days after fertilization, hAF/dAF), sequencing method, number of PCR cycles, and data source are indicated, along with SRA/BioSample/BioProject accessions for publicly released data.

| **species** | **cross** | **female strain** | **male strain** | **life cycle stage** | **stage** | **time** | **Extraction protocol** | **PCR cycles** | **Reference** | **SRA** | **BioSample** | **BioProject** |
| --- | --- | --- | --- | --- | --- | --- | --- | --- | --- | --- | --- | --- |
| ***S. promiscuus*** | NA | As9 | x | f-gam | female gamete | 0hAF | Direct Low input | 20 | this study | SRR37846298 | SAMN56777093 | PRJNA1445065 |
| ***S. promiscuus*** | NA | As9 | x | f-gam | female gamete | 0hAF | Direct Low input | 20 | this study | SRR37846297 | SAMN56777093 | PRJNA1445065 |
| ***S. promiscuus*** | NA | As9 | x | f-gam | female gamete | 0hAF | Direct Low input | 20 | this study | SRR37846251 | SAMN56777093 | PRJNA1445065 |
| ***S. promiscuus*** | NA | sxs103 | x | f-gam | female gamete | 0hAF | Direct Low input | 15 | Lotharukpong et al. in prep. |  |  |  |
| ***S. promiscuus*** | NA | sxs103 | x | f-gam | female gamete | 0hAF | Direct Low input | 15 | Lotharukpong et al. in prep. |  |  |  |
| ***S. promiscuus*** | NA | sxs103 | x | f-gam | female gamete | 0hAF | Direct Low input | 15 | Lotharukpong et al. in prep. |  |  |  |
| ***S. promiscuus*** | NA | x | As6 | m-gam | male gametes | 0hAF | Direct Low input | 15 | Lotharukpong et al. in prep. |  |  |  |
| ***S. promiscuus*** | NA | x | As6 | m-gam | male gametes | 0hAF | Direct Low input | 15 | Lotharukpong et al. in prep. |  |  |  |
| ***S. promiscuus*** | NA | x | As6 | m-gam | male gametes | 0hAF | Direct Low input | 15 | Lotharukpong et al. in prep. |  |  |  |
| ***S. promiscuus*** | NA | x | Mr8 | m-gam | male gamete | 0hAF | Direct Low input | 21 | this study | SRR37846215 | SAMN56777104 | PRJNA1445065 |
| ***S. promiscuus*** | NA | x | Mr8 | m-gam | male gamete | 0hAF | Direct Low input | 21 | this study | SRR37846204 | SAMN56777104 | PRJNA1445065 |
| ***S. promiscuus*** | NA | x | Mr8 | m-gam | male gamete | 0hAF | Direct Low input | 21 | this study | SRR37846193 | SAMN56777104 | PRJNA1445065 |
| ***S. promiscuus*** | cross | Mr5 | As6 | SP | zygote | 6hAF | Direct Low input | 20 | this study | SRR37846182 | SAMN56777103 | PRJNA1445065 |
| ***S. promiscuus*** | cross | Mr5 | As6 | SP | zygote | 6hAF | Direct Low input | 20 | this study | SRR37846171 | SAMN56777103 | PRJNA1445065 |
| ***S. promiscuus*** | cross | Mr5 | As6 | SP | zygote | 6hAF | Direct Low input | 20 | this study | SRR37846241 | SAMN56777103 | PRJNA1445065 |
| ***S. promiscuus*** | reciprocal cross | As9 | Mr8 | SP | zygote | 8hAF | Direct Low input | 21 | this study | SRR37846230 | SAMN56777098 | PRJNA1445065 |
| ***S. promiscuus*** | reciprocal cross | As9 | Mr8 | SP | zygote | 8hAF | Direct Low input | 21 | this study | SRR37846296 | SAMN56777098 | PRJNA1445065 |
| ***S. promiscuus*** | reciprocal cross | As9 | Mr8 | SP | zygote | 8hAF | Direct Low input | 21 | this study | SRR37846285 | SAMN56777098 | PRJNA1445065 |
| ***S. promiscuus*** | cross | Mr5 | As6 | SP | polarisation | 1dAF | Direct Low input | 20 | this study | SRR37846274 | SAMN56777102 | PRJNA1445065 |
| ***S. promiscuus*** | cross | Mr5 | As6 | SP | polarisation | 1dAF | Direct Low input | 20 | this study | SRR37846263 | SAMN56777102 | PRJNA1445065 |
| ***S. promiscuus*** | cross | Mr5 | As6 | SP | polarisation | 1dAF | Direct Low input | 20 | this study | SRR37846254 | SAMN56777097 | PRJNA1445065 |
| ***S. promiscuus*** | reciprocal cross | As9 | Mr8 | SP | polarisation | 1dAF | Direct Low input | 20 | this study | SRR37846256 | SAMN56777097 | PRJNA1445065 |
| ***S. promiscuus*** | reciprocal cross | As9 | Mr8 | SP | polarisation | 1dAF | Direct Low input | 20 | this study | SRR37846255 | SAMN56777097 | PRJNA1445065 |
| ***S. promiscuus*** | reciprocal cross | As9 | Mr8 | SP | polarisation | 1dAF | Direct Low input | 20 | this study | SRR37846257 | SAMN56777102 | PRJNA1445065 |
| ***S. promiscuus*** | cross | Mr5 | As6 | SP | 2-celled | 2dAF | Direct Low input | 20 | this study | SRR37846253 | SAMN56777099 | PRJNA1445065 |
| ***S. promiscuus*** | cross | Mr5 | As6 | SP | 2-celled | 2dAF | Direct Low input | 20 | this study | SRR37846252 | SAMN56777099 | PRJNA1445065 |
| ***S. promiscuus*** | cross | Mr5 | As6 | SP | 2-celled | 2dAF | Direct Low input | 20 | this study | SRR37846250 | SAMN56777099 | PRJNA1445065 |
| ***S. promiscuus*** | reciprocal cross | As9 | Mr8 | SP | 2-celled | 2dAF | Direct Low input | 18 | this study | SRR37846249 | SAMN56777094 | PRJNA1445065 |
| ***S. promiscuus*** | reciprocal cross | As9 | Mr8 | SP | 2-celled | 2dAF | Direct Low input | 20 | this study | SRR37846248 | SAMN56777094 | PRJNA1445065 |
| ***S. promiscuus*** | reciprocal cross | As9 | Mr8 | SP | 2-celled | 2dAF | Direct Low input | 20 | this study | SRR37846247 | SAMN56777094 | PRJNA1445065 |
| ***S. promiscuus*** | cross | Mr5 | As6 | SP | 4-celled | 8dAF | Direct Low input | 20 | this study | SRR37846246 | SAMN56777100 | PRJNA1445065 |
| ***S. promiscuus*** | cross | Mr5 | As6 | SP | 4-celled | 8dAF | Direct Low input | 20 | this study | SRR37846245 | SAMN56777100 | PRJNA1445065 |
| ***S. promiscuus*** | cross | Mr5 | As6 | SP | 4-celled | 8dAF | Direct Low input | 20 | this study | SRR37846220 | SAMN56777100 | PRJNA1445065 |
| ***S. promiscuus*** | reciprocal cross | As9 | Mr8 | SP | 4-celled | 8dAF | Direct Low input | 20 | this study | SRR37846218 | SAMN56777095 | PRJNA1445065 |
| ***S. promiscuus*** | reciprocal cross | As9 | Mr8 | SP | 4-celled | 8dAF | Direct Low input | 20 | this study | SRR37846217 | SAMN56777095 | PRJNA1445065 |
| ***S. promiscuus*** | reciprocal cross | As9 | Mr8 | SP | 4-celled | 8dAF | Direct Low input | 20 | this study | SRR37846216 | SAMN56777095 | PRJNA1445065 |
| ***S. promiscuus*** | cross | Mr5 | As6 | SP | patterning | 2wAF | Direct Low input | 20 | this study | SRR37846214 | SAMN56777101 | PRJNA1445065 |
| ***S. promiscuus*** | cross | Mr5 | As6 | SP | patterning | 2wAF | Direct Low input | 20 | this study | SRR37846213 | SAMN56777101 | PRJNA1445065 |
| ***S. promiscuus*** | reciprocal cross | As9 | Mr8 | SP | patterning | 2wAF | Direct Low input | 20 | this study | SRR37846212 | SAMN56777096 | PRJNA1445065 |
| ***S. promiscuus*** | reciprocal cross | As9 | Mr8 | SP | patterning | 2wAF | Direct Low input | 20 | this study | SRR37846211 | SAMN56777096 | PRJNA1445065 |
| ***S. promiscuus*** | reciprocal cross | As9 | Mr8 | SP | patterning | 2wAF | Direct Low input | 20 | this study | SRR37846210 | SAMN56777096 | PRJNA1445065 |
| ***U. pinnatifida*** | NA | UN1f | x | f-gam | female gamete | 0hAF | Direct Low input | 20 | this study | SRR37846209 | SAMN56777105 | PRJNA1445065 |
| ***U. pinnatifida*** | NA | UN1f | x | f-gam | female gamete | 0hAF | Direct Low input | 20 | this study | SRR37846208 | SAMN56777105 | PRJNA1445065 |
| ***U. pinnatifida*** | NA | UN1f | x | f-gam | female gamete | 0hAF | Direct Low input | 20 | this study | SRR37846207 | SAMN56777105 | PRJNA1445065 |
| ***U. pinnatifida*** | NA | x | UN1m | m-gam | male gamete | 0hAF | Direct Low input | 20 | this study | SRR37846206 | SAMN56777108 | PRJNA1445065 |
| ***U. pinnatifida*** | NA | x | UN1m | m-gam | male gamete | 0hAF | Direct Low input | 20 | this study | SRR37846205 | SAMN56777108 | PRJNA1445065 |
| ***U. pinnatifida*** | NA | x | UN1m | m-gam | male gamete | 0hAF | Direct Low input | 20 | this study | SRR37846203 | SAMN56777108 | PRJNA1445065 |
| ***U. pinnatifida*** | NA | x | UN1m | m-gam | male gamete | 0hAF | Direct Low input | 20 | this study | SRR37846202 | SAMN56777108 | PRJNA1445065 |
| ***U. pinnatifida*** | NA | x | UN1m | m-gam | male gamete | 0hAF | Direct Low input | 20 | this study | SRR37846201 | SAMN56777108 | PRJNA1445065 |
| ***U. pinnatifida*** | cross | UN1f | UnpinK2m | SP | zygote | 1hAF | Direct Low input | 20 | this study | SRR37846200 | SAMN56777114 | PRJNA1445065 |
| ***U. pinnatifida*** | cross | UN1f | UnpinK2m | SP | zygote | 1hAF | Direct Low input | 20 | this study | SRR37846199 | SAMN56777114 | PRJNA1445065 |
| ***U. pinnatifida*** | cross | UN1f | UnpinK2m | SP | zygote | 1hAF | Direct Low input | 20 | this study | SRR37846198 | SAMN56777114 | PRJNA1445065 |
| ***U. pinnatifida*** | reciprocal cross | UnpinK2f | UN1m | SP | zygote | 1hAF | Direct Low input | 20 | this study | SRR37846197 | SAMN56777120 | PRJNA1445065 |
| ***U. pinnatifida*** | reciprocal cross | UnpinK2f | UN1m | SP | zygote | 1hAF | Direct Low input | 20 | this study | SRR37846196 | SAMN56777120 | PRJNA1445065 |
| ***U. pinnatifida*** | reciprocal cross | UnpinK2f | UN1m | SP | zygote | 1hAF | Direct Low input | 20 | this study | SRR37846195 | SAMN56777120 | PRJNA1445065 |
| ***U. pinnatifida*** | cross | UN1f | UnpinK2m | SP | 2-celled | 10hAF | Direct Low input | 20 | this study | SRR37846194 | SAMN56777111 | PRJNA1445065 |
| ***U. pinnatifida*** | cross | UN1f | UnpinK2m | SP | 2-celled | 10hAF | Direct Low input | 20 | this study | SRR37846192 | SAMN56777111 | PRJNA1445065 |
| ***U. pinnatifida*** | cross | UN1f | UnpinK2m | SP | 2-celled | 10hAF | Direct Low input | 20 | this study | SRR37846191 | SAMN56777111 | PRJNA1445065 |
| ***U. pinnatifida*** | reciprocal cross | UnpinK2f | UN1m | SP | 2-celled | 10hAF | Direct Low input | 20 | this study | SRR37846190 | SAMN56777117 | PRJNA1445065 |
| ***U. pinnatifida*** | reciprocal cross | UnpinK2f | UN1m | SP | 2-celled | 10hAF | Direct Low input | 20 | this study | SRR37846189 | SAMN56777117 | PRJNA1445065 |
| ***U. pinnatifida*** | reciprocal cross | UnpinK2f | UN1m | SP | 2-celled | 10hAF | Direct Low input | 20 | this study | SRR37846188 | SAMN56777117 | PRJNA1445065 |
| ***U. pinnatifida*** | cross | UN1f | UnpinK2m | SP | 4-celled | 1dAF | Direct Low input | 20 | this study | SRR37846187 | SAMN56777112 | PRJNA1445065 |
| ***U. pinnatifida*** | cross | UN1f | UnpinK2m | SP | 4-celled | 1dAF | Direct Low input | 20 | this study | SRR37846186 | SAMN56777112 | PRJNA1445065 |
| ***U. pinnatifida*** | cross | UN1f | UnpinK2m | SP | 4-celled | 1dAF | Direct Low input | 20 | this study | SRR37846185 | SAMN56777112 | PRJNA1445065 |
| ***U. pinnatifida*** | reciprocal cross | UnpinK2f | UN1m | SP | 4-celled | 1dAF | Direct Low input | 20 | this study | SRR37846184 | SAMN56777118 | PRJNA1445065 |
| ***U. pinnatifida*** | reciprocal cross | UnpinK2f | UN1m | SP | 4-celled | 1dAF | Direct Low input | 20 | this study | SRR37846183 | SAMN56777118 | PRJNA1445065 |
| ***U. pinnatifida*** | reciprocal cross | UnpinK2f | UN1m | SP | 4-celled | 1dAF | Direct Low input | 20 | this study | SRR37846181 | SAMN56777118 | PRJNA1445065 |
| ***U. pinnatifida*** | cross | UN1f | UnpinK2m | SP | patterning | 2dAF | Direct Low input | 20 | this study | SRR37846180 | SAMN56777113 | PRJNA1445065 |
| ***U. pinnatifida*** | cross | UN1f | UnpinK2m | SP | patterning | 2dAF | Direct Low input | 20 | this study | SRR37846179 | SAMN56777113 | PRJNA1445065 |
| ***U. pinnatifida*** | cross | UN1f | UnpinK2m | SP | patterning | 2dAF | Direct Low input | 20 | this study | SRR37846178 | SAMN56777113 | PRJNA1445065 |
| ***U. pinnatifida*** | reciprocal cross | UnpinK2f | UN1m | SP | patterning | 2dAF | Direct Low input | 20 | this study | SRR37846177 | SAMN56777119 | PRJNA1445065 |
| ***U. pinnatifida*** | reciprocal cross | UnpinK2f | UN1m | SP | patterning | 2dAF | Direct Low input | 20 | this study | SRR37846176 | SAMN56777119 | PRJNA1445065 |
| ***U. pinnatifida*** | reciprocal cross | UnpinK2f | UN1m | SP | patterning | 2dAF | Direct Low input | 20 | this study | SRR37846175 | SAMN56777119 | PRJNA1445065 |
| ***D. dichotoma*** | NA | KB07f | x | f-gam | female gamete | 0hAF | Direct Low input | 18 | Lotharukpong et al. in prep. |  |  |  |
| ***D. dichotoma*** | NA | KB07f | x | f-gam | female gamete | 0hAF | Direct Low input | 18 | Lotharukpong et al. in prep. |  |  |  |
| ***D. dichotoma*** | NA | KB07f | x | f-gam | female gamete | 0hAF | Direct Low input | 18 | Lotharukpong et al. in prep. |  |  |  |
| ***D. dichotoma*** | NA | ODC2377 | x | f-gam | female gamete | 0hAF | Direct Low input | 21 | this study | SRR37846174 | SAMN56777075 | PRJNA1445065 |
| ***D. dichotoma*** | NA | ODC2377 | x | f-gam | female gamete | 0hAF | Direct Low input | 21 | this study | SRR37846173 | SAMN56777075 | PRJNA1445065 |
| ***D. dichotoma*** | NA | ODC2377 | x | f-gam | female gamete | 0hAF | Direct Low input | 21 | this study | SRR37846172 | SAMN56777075 | PRJNA1445065 |
| ***D. dichotoma*** | NA | x | KB07m | m-gam | male gamete | 0hAF | Direct Low input | 16 | Lotharukpong et al. in prep. |  |  |  |
| ***D. dichotoma*** | NA | x | KB07m | m-gam | male gamete | 0hAF | Direct Low input | 16 | Lotharukpong et al. in prep. |  |  |  |
| ***D. dichotoma*** | NA | x | KB07m | m-gam | male gamete | 0hAF | Direct Low input | 16 | Lotharukpong et al. in prep. |  |  |  |
| ***D. dichotoma*** | NA | x | KB07m | m-gam | male gamete | 0hAF | Direct Low input | 16 | Lotharukpong et al. in prep. |  |  |  |
| ***D. dichotoma*** | cross | KB07 | ODC1387 | SP | zygote | 1hAF | Direct Low input | 17 | this study | SRR37846170 | SAMN56777085 | PRJNA1445065 |
| ***D. dichotoma*** | cross | KB07 | ODC1387 | SP | zygote | 1hAF | Direct Low input | 17 | this study | SRR37846169 | SAMN56777085 | PRJNA1445065 |
| ***D. dichotoma*** | cross | KB07 | ODC1387 | SP | zygote | 1hAF | Direct Low input | 17 | this study | SRR37846168 | SAMN56777085 | PRJNA1445065 |
| ***D.dichotoma*** | reciprocal cross | ODC2377 | KB07 | SP | zygote | 1hAF | Direct Low input | 21 | this study | SRR37846167 | SAMN56777092 | PRJNA1445065 |
| ***D.dichotoma*** | reciprocal cross | ODC2377 | KB07 | SP | zygote | 1hAF | Direct Low input | 21 | this study | SRR37846166 | SAMN56777092 | PRJNA1445065 |
| ***D.dichotoma*** | reciprocal cross | ODC2377 | KB07 | SP | zygote | 1hAF | Direct Low input | 21 | this study | SRR37846165 | SAMN56777092 | PRJNA1445065 |
| ***D. dichotoma*** | internal cross | ODC1387 | KB07 | SP | zygote | 1hAF | Direct Low input | 17 | this study | SRR37846164 | SAMN56777081 | PRJNA1445065 |
| ***D. dichotoma*** | internal cross | ODC1387 | KB07 | SP | zygote | 1hAF | Direct Low input | 17 | this study | SRR37846244 | SAMN56777081 | PRJNA1445065 |
| ***D. dichotoma*** | internal cross | ODC1387 | KB07 | SP | zygote | 1hAF | Direct Low input | 17 | this study | SRR37846243 | SAMN56777081 | PRJNA1445065 |
| ***D. dichotoma*** | cross | KB07 | ODC1387 | SP | polarisation | 5.5hAF | Direct Low input | 17 | this study | SRR37846242 | SAMN56777084 | PRJNA1445065 |
| ***D. dichotoma*** | cross | KB07 | ODC1387 | SP | polarisation | 5.5hAF | Direct Low input | 17 | this study | SRR37846240 | SAMN56777084 | PRJNA1445065 |
| ***D. dichotoma*** | cross | KB07 | ODC1387 | SP | polarisation | 5.5hAF | Direct Low input | 17 | this study | SRR37846239 | SAMN56777084 | PRJNA1445065 |
| ***D.dichotoma*** | reciprocal cross | ODC2377 | KB07 | SP | polarisation | 5.5hAF | Direct Low input | 21 | this study | SRR37846238 | SAMN56777091 | PRJNA1445065 |
| ***D.dichotoma*** | reciprocal cross | ODC2377 | KB07 | SP | polarisation | 5.5hAF | Direct Low input | 21 | this study | SRR37846237 | SAMN56777091 | PRJNA1445065 |
| ***D.dichotoma*** | reciprocal cross | ODC2377 | KB07 | SP | polarisation | 5.5hAF | Direct Low input | 21 | this study | SRR37846236 | SAMN56777091 | PRJNA1445065 |
| ***D. dichotoma*** | internal cross | ODC1387 | KB07 | SP | polarisation | 5.5hAF | Direct Low input | 17 | this study | SRR37846235 | SAMN56777086 | PRJNA1445065 |
| ***D. dichotoma*** | internal cross | ODC1387 | KB07 | SP | polarisation | 5.5hAF | Direct Low input | 17 | this study | SRR37846234 | SAMN56777086 | PRJNA1445065 |
| ***D. dichotoma*** | cross | KB07 | ODC1387 | SP | 2-celled | 8hAF | Direct Low input | 17 | this study | SRR37846233 | SAMN56777082 | PRJNA1445065 |
| ***D. dichotoma*** | cross | KB07 | ODC1387 | SP | 2-celled | 8hAF | Direct Low input | 17 | this study | SRR37846232 | SAMN56777082 | PRJNA1445065 |
| ***D. dichotoma*** | cross | KB07 | ODC1387 | SP | 2-celled | 8hAF | Direct Low input | 17 | this study | SRR37846231 | SAMN56777082 | PRJNA1445065 |
| ***D. dichotoma*** | cross | KB07 | ODC1387 | SP | 2-celled | 8hAF | Direct Low input | 17 | this study | SRR37846229 | SAMN56777082 | PRJNA1445065 |
| ***D.dichotoma*** | reciprocal cross | ODC2377 | KB07 | SP | 2-celled | 8hAF | Direct Low input | 21 | this study | SRR37846228 | SAMN56777088 | PRJNA1445065 |
| ***D.dichotoma*** | reciprocal cross | ODC2377 | KB07 | SP | 2-celled | 8hAF | Direct Low input | 21 | this study | SRR37846227 | SAMN56777088 | PRJNA1445065 |
| ***D.dichotoma*** | reciprocal cross | ODC2377 | KB07 | SP | 2-celled | 8hAF | Direct Low input | 21 | this study | SRR37846226 | SAMN56777088 | PRJNA1445065 |
| ***D. dichotoma*** | internal cross | ODC1387 | KB07 | SP | 2-celled | 8hAF | Direct Low input | 17 | this study | SRR37846225 | SAMN56777078 | PRJNA1445065 |
| ***D. dichotoma*** | internal cross | ODC1387 | KB07 | SP | 2-celled | 8hAF | Direct Low input | 17 | this study | SRR37846224 | SAMN56777078 | PRJNA1445065 |
| ***D. dichotoma*** | internal cross | ODC1387 | KB07 | SP | 2-celled | 8hAF | Direct Low input | 17 | this study | SRR37846223 | SAMN56777078 | PRJNA1445065 |
| ***D. dichotoma*** | internal cross | ODC1388 | KB08 | SP | 2-celled | 8hAF | Direct Low input | 17 | this study | SRR37846222 | SAMN56777078 | PRJNA1445065 |
| ***D. dichotoma*** | cross | KB07 | ODC1387 | SP | 4-celled | 14hAF | Direct Low input | 18 | Lotharukpong et al. in prep. |  |  |  |
| ***D. dichotoma*** | cross | KB07 | ODC1387 | SP | 4-celled | 14hAF | Direct Low input | 18 | Lotharukpong et al. in prep. |  |  |  |
| ***D. dichotoma*** | cross | KB07 | ODC1387 | SP | 4-celled | 14hAF | Direct Low input | 18 | Lotharukpong et al. in prep. |  |  |  |
| ***D.dichotoma*** | reciprocal cross | ODC2377 | KB07 | SP | 4-celled | 14hAF | Direct Low input | 21 | this study | SRR37846221 | SAMN56777089 | PRJNA1445065 |
| ***D.dichotoma*** | reciprocal cross | ODC2377 | KB07 | SP | 4-celled | 14hAF | Direct Low input | 21 | this study | SRR37846219 | SAMN56777089 | PRJNA1445065 |
| ***D.dichotoma*** | reciprocal cross | ODC2377 | KB07 | SP | 4-celled | 14hAF | Direct Low input | 21 | this study | SRR37846295 | SAMN56777089 | PRJNA1445065 |
| ***D. dichotoma*** | internal cross | ODC1387 | KB07 | SP | 4-celled | 14hAF | Direct Low input | 17 | this study | SRR37846294 | SAMN56777079 | PRJNA1445065 |
| ***D. dichotoma*** | internal cross | ODC1387 | KB07 | SP | 4-celled | 14hAF | Direct Low input | 17 | this study | SRR37846293 | SAMN56777079 | PRJNA1445065 |
| ***D. dichotoma*** | internal cross | ODC1387 | KB07 | SP | 4-celled | 14hAF | Direct Low input | 17 | this study | SRR37846292 | SAMN56777079 | PRJNA1445065 |
| ***D. dichotoma*** | Cross | KB07 | ODC1387 | SP | patterning | 28hAF | Direct Low input | 17 | this study | SRR37846291 | SAMN56777083 | PRJNA1445065 |
| ***D. dichotoma*** | Cross | KB07 | ODC1387 | SP | patterning | 28hAF | Direct Low input | 17 | this study | SRR37846290 | SAMN56777083 | PRJNA1445065 |
| ***D. dichotoma*** | Cross | KB07 | ODC1387 | SP | patterning | 28hAF | Direct Low input | 17 | this study | SRR37846289 | SAMN56777083 | PRJNA1445065 |
| ***D. dichotoma*** | Cross | KB07 | ODC1387 | SP | patterning | 28hAF | Direct Low input | 17 | this study | SRR37846288 | SAMN56777083 | PRJNA1445065 |
| ***D. dichotoma*** | Cross | KB07 | ODC1387 | SP | patterning | 28hAF | Direct Low input | 17 | this study | SRR37846287 | SAMN56777083 | PRJNA1445065 |
| ***D.dichotoma*** | reciprocal cross | ODC2377 | KB07 | SP | patterning | 28hAF | Direct Low input | 21 | this study | SRR37846286 | SAMN56777090 | PRJNA1445065 |
| ***D.dichotoma*** | reciprocal cross | ODC2377 | KB07 | SP | patterning | 28hAF | Direct Low input | 21 | this study | SRR37846284 | SAMN56777090 | PRJNA1445065 |
| ***D.dichotoma*** | reciprocal cross | ODC2377 | KB07 | SP | patterning | 28hAF | Direct Low input | 21 | this study | SRR37846283 | SAMN56777090 | PRJNA1445065 |
| **D. dichotoma** | internal cross | KB07 | ODC1387 | SP | patterning | 28hAF | Direct Low input | 17 | this study | SRR37846282 | SAMN56777080 | PRJNA1445065 |

**Table S4. Phylostratigraphic age class enrichment within WGCNA co-expression modules.** For each species, the table reports the enrichment of evolutionarily old, mid-age, and young genes within each co-expression module. Columns indicate: module colour, age class (Old/Mid/Young), number of genes in the module (k), total number of expressed genes (m), observed and expected percentage of genes per age class, the deviation from the genomic background (delta percent), odds ratio, nominal p-value, and Benjamini–Hochberg adjusted p-value.

| ***Dictyota*** | | | | | | | | | |
| --- | --- | --- | --- | --- | --- | --- | --- | --- | --- |
| **module** | AgeClass | k | m | observed_percent | expected_percent | delta_percent | odds_ratio | pval | padj |
| **black** | Old | 390 | 792 | 49.24 | 54.09 | -4.85 | 0.816179258 | 0.005573097 | 0.007245027 |
| **black** | Mid | 271 | 792 | 34.22 | 27.12 | 7.09 | 1.420436806 | 8.05E-06 | 1.50E-05 |
| **black** | Young | 131 | 792 | 16.54 | 18.78 | -2.24 | 0.851170195 | 0.103514286 | 0.112140477 |
| **blue** | Old | 1571 | 2338 | 67.19 | 54.09 | 13.1 | 1.877804147 | 4.52E-43 | 5.88E-42 |
| **blue** | Mid | 516 | 2338 | 22.07 | 27.12 | -5.05 | 0.733133049 | 2.19E-09 | 5.69E-09 |
| **blue** | Young | 251 | 2338 | 10.74 | 18.78 | -8.05 | 0.482147044 | 1.88E-29 | 1.83E-28 |
| **brown** | Old | 888 | 1398 | 63.52 | 54.09 | 9.43 | 1.524914553 | 1.44E-13 | 5.63E-13 |
| **brown** | Mid | 338 | 1398 | 24.18 | 27.12 | -2.95 | 0.846236982 | 0.010244437 | 0.012888163 |
| **brown** | Young | 172 | 1398 | 12.3 | 18.78 | -6.48 | 0.585845843 | 1.78E-11 | 5.34E-11 |
| **green** | Old | 438 | 919 | 47.66 | 54.09 | -6.43 | 0.762368477 | 6.94E-05 | 0.000112731 |
| **green** | Mid | 255 | 919 | 27.75 | 27.12 | 0.62 | 1.033509814 | 0.675418412 | 0.675418412 |
| **green** | Young | 226 | 919 | 24.59 | 18.78 | 5.81 | 1.438790212 | 7.68E-06 | 1.50E-05 |
| **greenyellow** | Old | 302 | 371 | 81.4 | 54.09 | 27.31 | 3.799934195 | 1.33E-28 | 1.04E-27 |
| **greenyellow** | Mid | 57 | 371 | 15.36 | 27.12 | -11.76 | 0.481772334 | 6.58E-08 | 1.43E-07 |
| **greenyellow** | Young | 12 | 371 | 3.23 | 18.78 | -15.55 | 0.141524665 | 8.15E-20 | 5.30E-19 |
| **grey** | Old | 2246 | 5637 | 39.84 | 54.09 | -14.25 | 0.43355177 | 4.48E-147 | 1.75E-145 |
| **grey** | Mid | 1714 | 5637 | 30.41 | 27.12 | 3.28 | 1.265591629 | 3.76E-11 | 1.05E-10 |
| **grey** | Young | 1677 | 5637 | 29.75 | 18.78 | 10.97 | 2.623379951 | 6.18E-134 | 1.20E-132 |
| **magenta** | Old | 236 | 415 | 56.87 | 54.09 | 2.78 | 1.121908884 | 0.252388323 | 0.266030935 |
| **magenta** | Mid | 140 | 415 | 33.73 | 27.12 | 6.61 | 1.378427381 | 0.003020558 | 0.004062129 |
| **magenta** | Young | 39 | 415 | 9.4 | 18.78 | -9.39 | 0.442132329 | 1.47E-07 | 3.03E-07 |
| **pink** | Old | 344 | 571 | 60.25 | 54.09 | 6.15 | 1.296506217 | 0.002801879 | 0.003902617 |
| **pink** | Mid | 149 | 571 | 26.09 | 27.12 | -1.03 | 0.947031194 | 0.599056994 | 0.614821652 |
| **pink** | Young | 78 | 571 | 13.66 | 18.78 | -5.12 | 0.676709549 | 0.001079637 | 0.001619455 |
| **purple** | Old | 161 | 374 | 43.05 | 54.09 | -11.04 | 0.635636964 | 1.62E-05 | 2.88E-05 |
| **purple** | Mid | 123 | 374 | 32.89 | 27.12 | 5.76 | 1.324616099 | 0.013469753 | 0.015918799 |
| **purple** | Young | 90 | 374 | 24.06 | 18.78 | 5.28 | 1.380032818 | 0.010857816 | 0.013232963 |
| **red** | Old | 616 | 911 | 67.62 | 54.09 | 13.53 | 1.823535413 | 1.81E-17 | 1.01E-16 |
| **red** | Mid | 202 | 911 | 22.17 | 27.12 | -4.95 | 0.755537081 | 0.000497833 | 0.000776619 |
| **red** | Young | 93 | 911 | 10.21 | 18.78 | -8.58 | 0.477422402 | 4.91E-13 | 1.74E-12 |
| **tan** | Old | 170 | 216 | 78.7 | 54.09 | 24.61 | 3.173721036 | 5.89E-14 | 2.55E-13 |
| **tan** | Mid | 39 | 216 | 18.06 | 27.12 | -9.07 | 0.588747302 | 0.002030166 | 0.002932462 |
| **tan** | Young | 7 | 216 | 3.24 | 18.78 | -15.54 | 0.143065579 | 7.54E-12 | 2.45E-11 |
| **turquoise** | Old | 1888 | 3386 | 55.76 | 54.09 | 1.67 | 1.086141211 | 0.030905442 | 0.034437492 |
| **turquoise** | Mid | 973 | 3386 | 28.74 | 27.12 | 1.61 | 1.103694688 | 0.020746489 | 0.023797444 |
| **turquoise** | Young | 525 | 3386 | 15.51 | 18.78 | -3.28 | 0.756120933 | 4.05E-08 | 9.28E-08 |
| **yellow** | Old | 641 | 958 | 66.91 | 54.09 | 12.82 | 1.765817977 | 1.38E-16 | 6.71E-16 |
| **yellow** | Mid | 183 | 958 | 19.1 | 27.12 | -8.02 | 0.620415583 | 3.34E-09 | 8.14E-09 |
| **yellow** | Young | 134 | 958 | 13.99 | 18.78 | -4.8 | 0.691050177 | 6.32E-05 | 0.000107235 |
| ***Undaria*** | | | | | | | | | |
| **module** | AgeClass | k | m | observed_percent | expected_percent | delta_percent | odds_ratio | pval | padj |
| **black** | Old | 144 | 170 | 84.71 | 65.14 | 19.57 | 3.002524004 | 1.32E-08 | 6.78E-08 |
| **black** | Mid | 2.30E+01 | 1.70E+02 | 13.53 | 28.79 | -15.26 | 0.382770965 | 3.16E-06 | 1.14E-05 |
| **black** | Young | 3 | 170 | 1.76 | 6.07 | -4.31 | 0.274693088 | 0.013901772 | 0.023831609 |
| **blue** | Old | 9.67E+02 | 1.24E+03 | 77.98 | 65.14 | 12.85 | 2.028365042 | 3.16E-25 | 1.14E-23 |
| **blue** | Mid | 2.56E+02 | 1.24E+03 | 20.65 | 28.79 | -8.14 | 0.613708235 | 6.86E-12 | 6.17E-11 |
| **blue** | Young | 1.70E+01 | 1.24E+03 | 1.37 | 6.07 | -4.7 | 0.195393487 | 1.38E-17 | 1.65E-16 |
| **brown** | Old | 5.19E+02 | 7.99E+02 | 64.96 | 65.14 | -0.18 | 0.991474777 | 0.908151976 | 0.908151976 |
| **brown** | Mid | 248 | 799 | 31.04 | 28.79 | 2.25 | 1.122489677 | 0.145062945 | 0.193417261 |
| **brown** | Young | 3.20E+01 | 7.99E+02 | 4.01 | 6.07 | -2.07 | 0.628150901 | 0.00899637 | 0.016193465 |
| **green** | Old | 381 | 534 | 71.35 | 65.14 | 6.21 | 1.350586911 | 0.002119474 | 0.004768817 |
| **green** | Mid | 145 | 534 | 27.15 | 28.79 | -1.64 | 0.918471287 | 0.405805546 | 0.456531239 |
| **green** | Young | 8.00E+00 | 5.34E+02 | 1.5 | 6.07 | -4.58 | 0.226348583 | 2.17E-07 | 8.69E-07 |
| **greenyellow** | Old | 9.60E+01 | 1.24E+02 | 77.42 | 65.14 | 12.28 | 1.845779568 | 0.003270162 | 0.006540325 |
| **greenyellow** | Mid | 2.60E+01 | 1.24E+02 | 20.97 | 28.79 | -7.82 | 0.653543829 | 0.057842668 | 0.086764002 |
| **greenyellow** | Young | 2.00E+00 | 1.24E+02 | 1.61 | 6.07 | -4.46 | 0.251375548 | 0.034979785 | 0.054750968 |
| **grey** | Old | 2.25E+03 | 3.61E+03 | 62.44 | 65.14 | -2.7 | 0.842513258 | 4.68E-05 | 0.000153297 |
| **grey** | Mid | 1.07E+03 | 3.61E+03 | 29.74 | 28.79 | 0.95 | 1.069263248 | 0.131275261 | 0.181765746 |
| **grey** | Young | 2.82E+02 | 3.61E+03 | 7.82 | 6.07 | 1.75 | 1.523418173 | 2.10E-07 | 8.69E-07 |
| **magenta** | Old | 114 | 144 | 79.17 | 65.14 | 14.03 | 2.049682964 | 0.000282508 | 0.000782331 |
| **magenta** | Mid | 23 | 144 | 15.97 | 28.79 | -12.82 | 0.46650822 | 0.000401123 | 0.00103146 |
| **magenta** | Young | 7.00E+00 | 1.44E+02 | 4.86 | 6.07 | -1.21 | 0.787988803 | 0.724130186 | 0.74481962 |
| **pink** | Old | 127 | 167 | 76.05 | 65.14 | 10.91 | 1.71129844 | 0.00243965 | 0.005166317 |
| **pink** | Mid | 38 | 167 | 22.75 | 28.79 | -6.03 | 0.7255518 | 0.085486969 | 0.123101235 |
| **pink** | Young | 2 | 167 | 1.2 | 6.07 | -4.88 | 0.185118953 | 0.004643207 | 0.008797656 |
| **purple** | Old | 1.01E+02 | 1.42E+02 | 71.13 | 65.14 | 5.99 | 1.322794891 | 0.155960411 | 0.200520529 |
| **purple** | Mid | 36 | 142 | 25.35 | 28.79 | -3.44 | 0.838349397 | 0.401803238 | 0.456531239 |
| **purple** | Young | 5 | 142 | 3.52 | 6.07 | -2.55 | 0.561223599 | 0.285049524 | 0.353854582 |
| **red** | Old | 2.12E+02 | 2.81E+02 | 75.44 | 65.14 | 10.31 | 1.663128432 | 0.00022405 | 0.000672151 |
| **red** | Mid | 64 | 281 | 22.78 | 28.79 | -6.01 | 0.724198492 | 0.0233817 | 0.038260963 |
| **red** | Young | 5.00E+00 | 2.81E+02 | 1.78 | 6.07 | -4.29 | 0.274968317 | 0.000869646 | 0.002087151 |
| **turquoise** | Old | 2.06E+03 | 3.48E+03 | 59.15 | 65.14 | -5.98 | 0.690099808 | 1.30E-18 | 2.35E-17 |
| **turquoise** | Mid | 1132 | 3479 | 32.54 | 28.79 | 3.75 | 1.293188526 | 6.32E-09 | 3.79E-08 |
| **turquoise** | Young | 2.89E+02 | 3.48E+03 | 8.31 | 6.07 | 2.23 | 1.683448966 | 1.50E-10 | 1.08E-09 |
| **yellow** | Old | 526 | 823 | 63.91 | 65.14 | -1.22 | 0.943969141 | 0.447904804 | 0.488623422 |
| **yellow** | Mid | 250 | 823 | 30.38 | 28.79 | 1.59 | 1.085682399 | 0.299074498 | 0.358889397 |
| **yellow** | Young | 4.70E+01 | 8.23E+02 | 5.71 | 6.07 | -0.36 | 0.932006566 | 0.70513355 | 0.74481962 |
| ***Scytosiphon*** | | | | | | | | | |
| **module** | AgeClass | k | m | observed_percent | expected_percent | delta_percent | odds_ratio | pval | padj |
| **black** | Old | 327 | 516 | 63.37 | 55.72 | 7.66 | 1.38797568 | 0.000374036 | 0.000598458 |
| **black** | Mid | 135 | 516 | 26.16 | 29.8 | -3.63 | 0.830576535 | 0.070660448 | 0.084792538 |
| **black** | Young | 54 | 516 | 10.47 | 14.49 | -4.02 | 0.683188225 | 0.007579985 | 0.010355063 |
| **blue** | Old | 1504 | 2099 | 71.65 | 55.72 | 15.94 | 2.190214734 | 3.04E-57 | 4.86E-56 |
| **blue** | Mid | 417 | 2099 | 19.87 | 29.8 | -9.93 | 0.548537217 | 8.57E-28 | 5.87E-27 |
| **blue** | Young | 178 | 2099 | 8.48 | 14.49 | -6.01 | 0.51317583 | 1.39E-18 | 6.68E-18 |
| **brown** | Old | 937 | 1609 | 58.23 | 55.72 | 2.52 | 1.119568405 | 0.033054581 | 0.042881619 |
| **brown** | Mid | 504 | 1609 | 31.32 | 29.8 | 1.53 | 1.082570254 | 0.161406516 | 0.18446459 |
| **brown** | Young | 168 | 1609 | 10.44 | 14.49 | -4.05 | 0.666272977 | 6.13E-07 | 1.23E-06 |
| **cyan** | Old | 131 | 160 | 81.88 | 55.72 | 26.16 | 3.624886625 | 3.25E-12 | 1.04E-11 |
| **cyan** | Mid | 25 | 160 | 15.62 | 29.8 | -14.17 | 0.433677014 | 3.93E-05 | 6.74E-05 |
| **cyan** | Young | 4 | 160 | 2.5 | 14.49 | -11.99 | 0.150015707 | 6.62E-07 | 1.27E-06 |
| **green** | Old | 660 | 1351 | 48.85 | 55.72 | -6.86 | 0.741938606 | 1.52E-07 | 3.32E-07 |
| **green** | Mid | 446 | 1351 | 33.01 | 29.8 | 3.22 | 1.176027611 | 0.007766297 | 0.010355063 |
| **green** | Young | 245 | 1351 | 18.13 | 14.49 | 3.65 | 1.339764261 | 0.000111481 | 0.00018452 |
| **greenyellow** | Old | 193 | 241 | 80.08 | 55.72 | 24.37 | 3.239289262 | 2.01E-15 | 8.75E-15 |
| **greenyellow** | Mid | 35 | 241 | 14.52 | 29.8 | -15.27 | 0.396349151 | 3.43E-08 | 8.24E-08 |
| **greenyellow** | Young | 13 | 241 | 5.39 | 14.49 | -9.09 | 0.333150838 | 1.17E-05 | 2.08E-05 |
| **grey** | Old | 1448 | 3257 | 44.46 | 55.72 | -11.26 | 0.57375934 | 3.76E-46 | 3.61E-45 |
| **grey** | Mid | 998 | 3257 | 30.64 | 29.8 | 0.85 | 1.050494025 | 0.243616617 | 0.260836837 |
| **grey** | Young | 811 | 3257 | 24.9 | 14.49 | 10.41 | 2.398101404 | 2.00E-69 | 4.81E-68 |
| **magenta** | Old | 244 | 335 | 72.84 | 55.72 | 17.12 | 2.159674892 | 8.06E-11 | 2.15E-10 |
| **magenta** | Mid | 63 | 335 | 18.81 | 29.8 | -10.99 | 0.540280123 | 4.20E-06 | 7.75E-06 |
| **magenta** | Young | 28 | 335 | 8.36 | 14.49 | -6.13 | 0.533214398 | 0.000730831 | 0.001131609 |
| **midnightblue** | Old | 109 | 135 | 80.74 | 55.72 | 25.02 | 3.357850611 | 1.45E-09 | 3.65E-09 |
| **midnightblue** | Mid | 24 | 135 | 17.78 | 29.8 | -12.02 | 0.507234658 | 0.00173526 | 0.00260289 |
| **midnightblue** | Young | 2 | 135 | 1.48 | 14.49 | -13.01 | 0.088045764 | 4.03E-07 | 8.41E-07 |
| **pink** | Old | 339 | 475 | 71.37 | 55.72 | 15.65 | 2.015955878 | 1.45E-12 | 4.97E-12 |
| **pink** | Mid | 112 | 475 | 23.58 | 29.8 | -6.22 | 0.721069716 | 0.002656638 | 0.003750548 |
| **pink** | Young | 24 | 475 | 5.05 | 14.49 | -9.44 | 0.307614541 | 5.20E-11 | 1.56E-10 |
| **purple** | Old | 140 | 249 | 56.22 | 55.72 | 0.51 | 1.02116687 | 0.897864359 | 0.897864359 |
| **purple** | Mid | 89 | 249 | 35.74 | 29.8 | 5.95 | 1.315989575 | 0.042870058 | 0.054151652 |
| **purple** | Young | 20 | 249 | 8.03 | 14.49 | -6.46 | 0.511665887 | 0.002630713 | 0.003750548 |
| **red** | Old | 783 | 1043 | 75.07 | 55.72 | 19.36 | 2.513341283 | 1.98E-40 | 1.59E-39 |
| **red** | Mid | 211 | 1043 | 20.23 | 29.8 | -9.57 | 0.580829335 | 6.87E-13 | 2.54E-12 |
| **red** | Young | 49 | 1043 | 4.7 | 14.49 | -9.79 | 0.277163123 | 2.28E-25 | 1.37E-24 |
| **salmon** | Old | 86 | 170 | 50.59 | 55.72 | -5.13 | 0.812119218 | 0.187428301 | 0.20922229 |
| **salmon** | Mid | 62 | 170 | 36.47 | 29.8 | 6.68 | 1.356836948 | 0.063456556 | 0.078100376 |
| **salmon** | Young | 22 | 170 | 12.94 | 14.49 | -1.55 | 0.876275963 | 0.661255084 | 0.675324341 |
| **tan** | Old | 117 | 218 | 53.67 | 55.72 | -2.05 | 0.919783675 | 0.537984852 | 0.561375498 |
| **tan** | Mid | 76 | 218 | 34.86 | 29.8 | 5.07 | 1.264922502 | 0.101552904 | 0.118891205 |
| **tan** | Young | 25 | 218 | 11.47 | 14.49 | -3.02 | 0.762192679 | 0.244534535 | 0.260836837 |
| **turquoise** | Old | 1880 | 4417 | 42.56 | 55.72 | -13.15 | 0.492399816 | 2.77E-91 | 1.33E-89 |
| **turquoise** | Mid | 1742 | 4417 | 39.44 | 29.8 | 9.64 | 1.797311321 | 6.93E-57 | 8.31E-56 |
| **turquoise** | Young | 795 | 4417 | 18 | 14.49 | 3.51 | 1.427780754 | 6.99E-14 | 2.80E-13 |
| **yellow** | Old | 981 | 1456 | 67.38 | 55.72 | 11.66 | 1.712175346 | 3.18E-21 | 1.70E-20 |
| **yellow** | Mid | 344 | 1456 | 23.63 | 29.8 | -6.17 | 0.710039298 | 4.99E-08 | 1.14E-07 |
| **yellow** | Young | 131 | 1456 | 9 | 14.49 | -5.49 | 0.561154819 | 7.18E-11 | 2.03E-10 |

**Table S5.** **Phylostratigraphic age class enrichment within hub genes of WGCNA co-expression modules**. For each species, the table reports the enrichment of evolutionarily old, mid-age, and young genes among hub genes (top 25% by absolute module membership, |MM|) relative to the age composition of their parent module. Columns indicate: module colour, age class (Old/Mid/Young), number of hub genes in the age class (hub_count), number of module genes in the age class (bg_count), total number of age-assigned hub genes in the module (hub_total), total number of age-assigned genes in the module (bg_total), observed and expected percentage of hub genes per age class, the deviation from the module background (delta percent), percentage enrichment relative to expectation (pct_enrichment), log2 enrichment, nominal p-value (two-sided Fisher's exact test), and Benjamini–Hochberg adjusted p-value (applied within module). Top-fraction threshold (top_frac = 0.25) indicates the |MM| cutoff used to define hub genes.

| ***Dictyota*** | | | | | | | | | | | | | |
| --- | --- | --- | --- | --- | --- | --- | --- | --- | --- | --- | --- | --- | --- |
| **module** | AgeClass | hub_count | bg_count | hub_total | bg_total | observed_percent | expected_percent | delta_percent | pct_enrichment | log2_enrich | p_value | p_adj | top_frac |
| **black** | Mid | 56 | 271 | 198 | 792 | 28.28 | 34.22 | -5.93 | -17.34 | -0.274794119 | 0.046644071 | 0.069966106 | 0.25 |
| **black** | Old | 111 | 390 | 198 | 792 | 56.06 | 49.24 | 6.82 | 13.85 | 0.187085553 | 0.032703878 | 0.069966106 | 0.25 |
| **black** | Young | 31 | 131 | 198 | 792 | 15.66 | 16.54 | -0.88 | -5.34 | -0.079226691 | 0.741467295 | 0.741467295 | 0.25 |
| **blue** | Mid | 87 | 516 | 585 | 2338 | 14.87 | 22.07 | -7.2 | -32.62 | -0.56951736 | 6.84E-07 | 6.84E-07 | 0.25 |
| **blue** | Old | 483 | 1571 | 585 | 2338 | 82.56 | 67.19 | 15.37 | 22.87 | 0.297178314 | 3.47E-21 | 1.04E-20 | 0.25 |
| **blue** | Young | 15 | 251 | 585 | 2338 | 2.56 | 10.74 | -8.17 | -76.12 | -2.065886558 | 1.82E-16 | 2.72E-16 | 0.25 |
| **brown** | Mid | 88 | 338 | 348 | 1398 | 25.29 | 24.18 | 1.11 | 4.59 | 0.064757332 | 0.613134737 | 0.613134737 | 0.25 |
| **brown** | Old | 238 | 888 | 348 | 1398 | 68.39 | 63.52 | 4.87 | 7.67 | 0.106607046 | 0.033885404 | 0.050828106 | 0.25 |
| **brown** | Young | 22 | 172 | 348 | 1398 | 6.32 | 12.3 | -5.98 | -48.62 | -0.960627987 | 4.54E-05 | 0.000136259 | 0.25 |
| **green** | Mid | 58 | 255 | 230 | 919 | 25.22 | 27.75 | -2.53 | -9.12 | -0.137941441 | 0.349993733 | 0.349993733 | 0.25 |
| **green** | Old | 127 | 438 | 230 | 919 | 55.22 | 47.66 | 7.56 | 15.86 | 0.212328628 | 0.009468706 | 0.028406118 | 0.25 |
| **green** | Young | 45 | 226 | 230 | 919 | 19.57 | 24.59 | -5.03 | -20.44 | -0.329894866 | 0.042201287 | 0.063301931 | 0.25 |
| **greenyellow** | Mid | 16 | 57 | 92 | 371 | 17.39 | 15.36 | 2.03 | 13.2 | 0.178823406 | 0.510238496 | 0.759702283 | 0.25 |
| **greenyellow** | Old | 74 | 302 | 92 | 371 | 80.43 | 81.4 | -0.97 | -1.19 | -0.017237953 | 0.759702283 | 0.759702283 | 0.25 |
| **greenyellow** | Young | 2 | 12 | 92 | 371 | 2.17 | 3.23 | -1.06 | -32.79 | -0.57324908 | 0.737638416 | 0.759702283 | 0.25 |
| **grey** | Mid | 427 | 1714 | 1410 | 5637 | 30.28 | 30.41 | -0.12 | -0.4 | -0.00582673 | 0.920147149 | 0.920147149 | 0.25 |
| **grey** | Old | 651 | 2246 | 1410 | 5637 | 46.17 | 39.84 | 6.33 | 15.88 | 0.212603926 | 2.62E-08 | 3.94E-08 | 0.25 |
| **grey** | Young | 332 | 1677 | 1410 | 5637 | 23.55 | 29.75 | -6.2 | -20.85 | -0.337395137 | 2.50E-09 | 7.51E-09 | 0.25 |
| **magenta** | Mid | 38 | 140 | 103 | 415 | 36.89 | 33.73 | 3.16 | 9.36 | 0.129111496 | 0.471350512 | 0.646522075 | 0.25 |
| **magenta** | Old | 61 | 236 | 103 | 415 | 59.22 | 56.87 | 2.36 | 4.14 | 0.058561287 | 0.646522075 | 0.646522075 | 0.25 |
| **magenta** | Young | 4 | 39 | 103 | 415 | 3.88 | 9.4 | -5.51 | -58.68 | -1.27493522 | 0.030789888 | 0.092369665 | 0.25 |
| **pink** | Mid | 27 | 149 | 143 | 571 | 18.88 | 26.09 | -7.21 | -27.64 | -0.46680542 | 0.02751683 | 0.02751683 | 0.25 |
| **pink** | Old | 105 | 344 | 143 | 571 | 73.43 | 60.25 | 13.18 | 21.88 | 0.285456362 | 0.000174493 | 0.000523479 | 0.25 |
| **pink** | Young | 11 | 78 | 143 | 571 | 7.69 | 13.66 | -5.97 | -43.69 | -0.828495002 | 0.016492852 | 0.024739278 | 0.25 |
| **purple** | Mid | 32 | 123 | 93 | 374 | 34.41 | 32.89 | 1.52 | 4.62 | 0.065221143 | 0.7990927 | 0.7990927 | 0.25 |
| **purple** | Old | 46 | 161 | 93 | 374 | 49.46 | 43.05 | 6.41 | 14.9 | 0.200380727 | 0.183714811 | 0.275572216 | 0.25 |
| **purple** | Young | 15 | 90 | 93 | 374 | 16.13 | 24.06 | -7.94 | -32.97 | -0.577226852 | 0.049447797 | 0.148343391 | 0.25 |
| **red** | Mid | 41 | 202 | 227 | 911 | 18.06 | 22.17 | -4.11 | -18.54 | -0.295900722 | 0.096863054 | 0.096863054 | 0.25 |
| **red** | Old | 184 | 616 | 227 | 911 | 81.06 | 67.62 | 13.44 | 19.88 | 0.261534172 | 3.30E-07 | 4.95E-07 | 0.25 |
| **red** | Young | 2 | 93 | 227 | 911 | 0.88 | 10.21 | -9.33 | -91.37 | -3.534400055 | 5.62E-10 | 1.69E-09 | 0.25 |
| **tan** | Mid | 8 | 39 | 54 | 216 | 14.81 | 18.06 | -3.24 | -17.95 | -0.285402219 | 0.545091785 | 0.545091785 | 0.25 |
| **tan** | Old | 46 | 170 | 54 | 216 | 85.19 | 78.7 | 6.48 | 8.24 | 0.11417102 | 0.248977962 | 0.373466943 | 0.25 |
| **tan** | Young | 0 | 7 | 54 | 216 | 0 | 3.24 | -3.24 | -100 | NA | 0.196435819 | 0.373466943 | 0.25 |
| **turquoise** | Mid | 216 | 973 | 847 | 3386 | 25.5 | 28.74 | -3.23 | -11.25 | -0.172260394 | 0.017861615 | 0.017861615 | 0.25 |
| **turquoise** | Old | 526 | 1888 | 847 | 3386 | 62.1 | 55.76 | 6.34 | 11.37 | 0.155424038 | 1.88E-05 | 5.65E-05 | 0.25 |
| **turquoise** | Young | 105 | 525 | 847 | 3386 | 12.4 | 15.51 | -3.11 | -20.05 | -0.322779996 | 0.003650215 | 0.005475323 | 0.25 |
| **yellow** | Mid | 45 | 183 | 239 | 958 | 18.83 | 19.1 | -0.27 | -1.43 | -0.020831704 | 1 | 1 | 0.25 |
| **yellow** | Old | 185 | 641 | 239 | 958 | 77.41 | 66.91 | 10.5 | 15.69 | 0.210215952 | 6.81E-05 | 0.000102205 | 0.25 |
| **yellow** | Young | 9 | 134 | 239 | 958 | 3.77 | 13.99 | -10.22 | -73.08 | -1.893149151 | 1.04E-08 | 3.11E-08 | 0.25 |
| ***Undaria*** | | | | | | | | | | | |  |  |
| **module** | AgeClass | hub_count | bg_count | hub_total | bg_total | observed_percent | expected_percent | delta_percent | pct_enrichment | log2_enrich | p_value | p_adj | top_frac |
| **black** | Mid | 7 | 23 | 48 | 170 | 14.58 | 13.53 | 1.05 | 7.79 | 0.108221401 | 0.806156989 | 1 | 0.25 |
| **black** | Old | 40 | 144 | 48 | 170 | 83.33 | 84.71 | -1.37 | -1.62 | -0.023568471 | 0.813840872 | 1 | 0.25 |
| **black** | Young | 1 | 3 | 48 | 170 | 2.08 | 1.76 | 0.32 | 18.06 | 0.239465935 | 1 | 1 | 0.25 |
| **blue** | Mid | 52 | 256 | 321 | 1240 | 16.2 | 20.65 | -4.45 | -21.53 | -0.349865364 | 0.024745716 | 0.037118574 | 0.25 |
| **blue** | Old | 267 | 967 | 321 | 1240 | 83.18 | 77.98 | 5.19 | 6.66 | 0.09301877 | 0.009717211 | 0.029151632 | 0.25 |
| **blue** | Young | 2 | 17 | 321 | 1240 | 0.62 | 1.37 | -0.75 | -54.55 | -1.137767923 | 0.265621658 | 0.265621658 | 0.25 |
| **brown** | Mid | 69 | 248 | 209 | 799 | 33.01 | 31.04 | 1.98 | 6.36 | 0.089020707 | 0.487061587 | 0.500422063 | 0.25 |
| **brown** | Old | 140 | 519 | 209 | 799 | 66.99 | 64.96 | 2.03 | 3.12 | 0.044384849 | 0.500422063 | 0.500422063 | 0.25 |
| **brown** | Young | 0 | 32 | 209 | 799 | 0 | 4.01 | -4.01 | -100 | NA | 0.0001003 | 0.0003009 | 0.25 |
| **green** | Mid | 38 | 145 | 131 | 534 | 29.01 | 27.15 | 1.85 | 6.83 | 0.095291354 | 0.574030337 | 0.7398412 | 0.25 |
| **green** | Old | 92 | 381 | 131 | 534 | 70.23 | 71.35 | -1.12 | -1.57 | -0.022812301 | 0.7398412 | 0.7398412 | 0.25 |
| **green** | Young | 1 | 8 | 131 | 534 | 0.76 | 1.5 | -0.73 | -49.05 | -0.97272707 | 0.6863008 | 0.7398412 | 0.25 |
| **greenyellow** | Mid | 6 | 26 | 28 | 124 | 21.43 | 20.97 | 0.46 | 2.2 | 0.031364171 | 1 | 1 | 0.25 |
| **greenyellow** | Old | 21 | 96 | 28 | 124 | 75 | 77.42 | -2.42 | -3.12 | -0.04580369 | 0.798283716 | 1 | 0.25 |
| **greenyellow** | Young | 1 | 2 | 28 | 124 | 3.57 | 1.61 | 1.96 | 121.43 | 1.146841388 | 0.402045633 | 1 | 0.25 |
| **grey** | Mid | 274 | 1072 | 939 | 3605 | 29.18 | 29.74 | -0.56 | -1.87 | -0.027254911 | 0.678221369 | 0.678221369 | 0.25 |
| **grey** | Old | 614 | 2251 | 939 | 3605 | 65.39 | 62.44 | 2.95 | 4.72 | 0.0665467 | 0.031174434 | 0.046761651 | 0.25 |
| **grey** | Young | 51 | 282 | 939 | 3605 | 5.43 | 7.82 | -2.39 | -30.57 | -0.526323814 | 0.001421761 | 0.004265282 | 0.25 |
| **magenta** | Mid | 2 | 23 | 40 | 144 | 5 | 15.97 | -10.97 | -68.7 | -1.67556505 | 0.024357598 | 0.036536398 | 0.25 |
| **magenta** | Old | 38 | 114 | 40 | 144 | 95 | 79.17 | 15.83 | 20 | 0.263034406 | 0.002708569 | 0.008125706 | 0.25 |
| **magenta** | Young | 0 | 7 | 40 | 144 | 0 | 4.86 | -4.86 | -100 | NA | 0.190401004 | 0.190401004 | 0.25 |
| **pink** | Mid | 6 | 38 | 40 | 167 | 15 | 22.75 | -7.75 | -34.08 | -0.601188815 | 0.202179179 | 0.42276892 | 0.25 |
| **pink** | Old | 33 | 127 | 40 | 167 | 82.5 | 76.05 | 6.45 | 8.48 | 0.11748563 | 0.395289458 | 0.42276892 | 0.25 |
| **pink** | Young | 1 | 2 | 40 | 167 | 2.5 | 1.2 | 1.3 | 108.75 | 1.061776198 | 0.42276892 | 0.42276892 | 0.25 |
| **purple** | Mid | 11 | 36 | 37 | 142 | 29.73 | 25.35 | 4.38 | 17.27 | 0.229800371 | 0.513018553 | 1 | 0.25 |
| **purple** | Old | 25 | 101 | 37 | 142 | 67.57 | 71.13 | -3.56 | -5 | -0.074061539 | 0.673624566 | 1 | 0.25 |
| **purple** | Young | 1 | 5 | 37 | 142 | 2.7 | 3.52 | -0.82 | -23.24 | -0.381634341 | 1 | 1 | 0.25 |
| **red** | Mid | 14 | 64 | 71 | 281 | 19.72 | 22.78 | -3.06 | -13.42 | -0.207965877 | 0.517128132 | 0.517128132 | 0.25 |
| **red** | Old | 57 | 212 | 71 | 281 | 80.28 | 75.44 | 4.84 | 6.41 | 0.08964876 | 0.3389904 | 0.5084856 | 0.25 |
| **red** | Young | 0 | 5 | 71 | 281 | 0 | 1.78 | -1.78 | -100 | NA | 0.334736611 | 0.5084856 | 0.25 |
| **turquoise** | Mid | 305 | 1132 | 953 | 3479 | 32 | 32.54 | -0.53 | -1.64 | -0.023868251 | 0.685204 | 1 | 0.25 |
| **turquoise** | Old | 569 | 2058 | 953 | 3479 | 59.71 | 59.15 | 0.55 | 0.93 | 0.013382135 | 0.699063446 | 1 | 0.25 |
| **turquoise** | Young | 79 | 289 | 953 | 3479 | 8.29 | 8.31 | -0.02 | -0.21 | -0.003020375 | 1 | 1 | 0.25 |
| **yellow** | Mid | 62 | 250 | 215 | 823 | 28.84 | 30.38 | -1.54 | -5.07 | -0.075032203 | 0.605184577 | 0.907776865 | 0.25 |
| **yellow** | Old | 141 | 526 | 215 | 823 | 65.58 | 63.91 | 1.67 | 2.61 | 0.037188134 | 0.564315678 | 0.907776865 | 0.25 |
| **yellow** | Young | 12 | 47 | 215 | 823 | 5.58 | 5.71 | -0.13 | -2.27 | -0.03307058 | 1 | 1 | 0.25 |
| ***Scytosiphon*** | | | | | | | | | | | | | |
| **module** | AgeClass | hub_count | bg_count | hub_total | bg_total | observed_percent | expected_percent | delta_percent | pct_enrichment | log2_enrich | p_value | p_adj | top_frac |
| **black** | Mid | 23 | 135 | 129 | 516 | 17.83 | 26.16 | -8.33 | -31.85 | -0.553253641 | 0.014862155 | 0.022293233 | 0.25 |
| **black** | Old | 97 | 327 | 129 | 516 | 75.19 | 63.37 | 11.82 | 18.65 | 0.246766017 | 0.001484224 | 0.004452672 | 0.25 |
| **black** | Young | 9 | 54 | 129 | 516 | 6.98 | 10.47 | -3.49 | -33.33 | -0.584962501 | 0.182873645 | 0.182873645 | 0.25 |
| **blue** | Mid | 71 | 417 | 524 | 2099 | 13.55 | 19.87 | -6.32 | -31.8 | -0.552093004 | 2.15E-05 | 2.15E-05 | 0.25 |
| **blue** | Old | 435 | 1504 | 524 | 2099 | 83.02 | 71.65 | 11.36 | 15.86 | 0.212346189 | 6.60E-12 | 1.98E-11 | 0.25 |
| **blue** | Young | 18 | 178 | 524 | 2099 | 3.44 | 8.48 | -5.05 | -59.49 | -1.30374498 | 2.92E-07 | 4.38E-07 | 0.25 |
| **brown** | Mid | 136 | 504 | 402 | 1609 | 33.83 | 31.32 | 2.51 | 8 | 0.111079837 | 0.214831271 | 0.214831271 | 0.25 |
| **brown** | Old | 251 | 937 | 402 | 1609 | 62.44 | 58.23 | 4.2 | 7.22 | 0.100535236 | 0.053955604 | 0.080933406 | 0.25 |
| **brown** | Young | 15 | 168 | 402 | 1609 | 3.73 | 10.44 | -6.71 | -64.26 | -1.484529908 | 4.88E-08 | 1.46E-07 | 0.25 |
| **cyan** | Mid | 3 | 25 | 40 | 160 | 7.5 | 15.62 | -8.12 | -52 | -1.058893689 | 0.132935784 | 0.236002058 | 0.25 |
| **cyan** | Old | 36 | 131 | 40 | 160 | 90 | 81.88 | 8.13 | 9.92 | 0.136502 | 0.157334705 | 0.236002058 | 0.25 |
| **cyan** | Young | 1 | 4 | 40 | 160 | 2.5 | 2.5 | 0 | 0 | 0 | 1 | 1 | 0.25 |
| **green** | Mid | 128 | 446 | 337 | 1351 | 37.98 | 33.01 | 4.97 | 15.05 | 0.202307278 | 0.027382421 | 0.027382421 | 0.25 |
| **green** | Old | 184 | 660 | 337 | 1351 | 54.6 | 48.85 | 5.75 | 11.76 | 0.16044692 | 0.016791837 | 0.025187755 | 0.25 |
| **green** | Young | 25 | 245 | 337 | 1351 | 7.42 | 18.13 | -10.72 | -59.09 | -1.289574571 | 3.40E-10 | 1.02E-09 | 0.25 |
| **greenyellow** | Mid | 7 | 35 | 59 | 241 | 11.86 | 14.52 | -2.66 | -18.31 | -0.291681808 | 0.670994648 | 0.85294721 | 0.25 |
| **greenyellow** | Old | 48 | 193 | 59 | 241 | 81.36 | 80.08 | 1.27 | 1.59 | 0.02275175 | 0.85294721 | 0.85294721 | 0.25 |
| **greenyellow** | Young | 4 | 13 | 59 | 241 | 6.78 | 5.39 | 1.39 | 25.68 | 0.329806569 | 0.52625133 | 0.85294721 | 0.25 |
| **grey** | Mid | 237 | 998 | 814 | 3257 | 29.12 | 30.64 | -1.53 | -4.98 | -0.073709736 | 0.292156008 | 0.292156008 | 0.25 |
| **grey** | Old | 404 | 1448 | 814 | 3257 | 49.63 | 44.46 | 5.17 | 11.64 | 0.158808616 | 0.000625465 | 0.001876395 | 0.25 |
| **grey** | Young | 173 | 811 | 814 | 3257 | 21.25 | 24.9 | -3.65 | -14.65 | -0.228486856 | 0.005733282 | 0.008599923 | 0.25 |
| **magenta** | Mid | 5 | 63 | 83 | 335 | 6.02 | 18.81 | -12.78 | -67.97 | -1.642373975 | 0.000310073 | 0.00046511 | 0.25 |
| **magenta** | Old | 75 | 244 | 83 | 335 | 90.36 | 72.84 | 17.53 | 24.06 | 0.311059207 | 1.52E-05 | 4.57E-05 | 0.25 |
| **magenta** | Young | 3 | 28 | 83 | 335 | 3.61 | 8.36 | -4.74 | -56.76 | -1.209414567 | 0.106615239 | 0.106615239 | 0.25 |
| **midnightblue** | Mid | 8 | 24 | 33 | 135 | 24.24 | 17.78 | 6.46 | 36.36 | 0.447458977 | 0.297731136 | 0.673144929 | 0.25 |
| **midnightblue** | Old | 25 | 109 | 33 | 135 | 75.76 | 80.74 | -4.98 | -6.17 | -0.091906657 | 0.448763286 | 0.673144929 | 0.25 |
| **midnightblue** | Young | 0 | 2 | 33 | 135 | 0 | 1.48 | -1.48 | -100 | NA | 1 | 1 | 0.25 |
| **pink** | Mid | 21 | 112 | 119 | 475 | 17.65 | 23.58 | -5.93 | -25.16 | -0.418071559 | 0.082033928 | 0.082033928 | 0.25 |
| **pink** | Old | 97 | 339 | 119 | 475 | 81.51 | 71.37 | 10.14 | 14.21 | 0.191737319 | 0.004822394 | 0.014467182 | 0.25 |
| **pink** | Young | 1 | 24 | 119 | 475 | 0.84 | 5.05 | -4.21 | -83.37 | -2.587996561 | 0.013729411 | 0.020594116 | 0.25 |
| **purple** | Mid | 17 | 89 | 62 | 249 | 27.42 | 35.74 | -8.32 | -23.29 | -0.382464968 | 0.127990965 | 0.360167806 | 0.25 |
| **purple** | Old | 39 | 140 | 62 | 249 | 62.9 | 56.22 | 6.68 | 11.88 | 0.161924824 | 0.240111871 | 0.360167806 | 0.25 |
| **purple** | Young | 6 | 20 | 62 | 249 | 9.68 | 8.03 | 1.65 | 20.48 | 0.268840028 | 0.59352435 | 0.59352435 | 0.25 |
| **red** | Mid | 40 | 211 | 260 | 1043 | 15.38 | 20.23 | -4.85 | -23.95 | -0.395015464 | 0.025870306 | 0.025870306 | 0.25 |
| **red** | Old | 215 | 783 | 260 | 1043 | 82.69 | 75.07 | 7.62 | 10.15 | 0.139479982 | 0.000916829 | 0.002750488 | 0.25 |
| **red** | Young | 5 | 49 | 260 | 1043 | 1.92 | 4.7 | -2.77 | -59.07 | -1.28862612 | 0.016567726 | 0.024851589 | 0.25 |
| **salmon** | Mid | 16 | 62 | 42 | 170 | 38.1 | 36.47 | 1.62 | 4.45 | 0.062877203 | 0.854223186 | 1 | 0.25 |
| **salmon** | Old | 21 | 86 | 42 | 170 | 50 | 50.59 | -0.59 | -1.16 | -0.016873819 | 1 | 1 | 0.25 |
| **salmon** | Young | 5 | 22 | 42 | 170 | 11.9 | 12.94 | -1.04 | -8.01 | -0.12043001 | 1 | 1 | 0.25 |
| **tan** | Mid | 26 | 76 | 54 | 218 | 48.15 | 34.86 | 13.29 | 38.11 | 0.465809027 | 0.021580375 | 0.060381667 | 0.25 |
| **tan** | Old | 22 | 117 | 54 | 218 | 40.74 | 53.67 | -12.93 | -24.09 | -0.397636278 | 0.040254445 | 0.060381667 | 0.25 |
| **tan** | Young | 6 | 25 | 54 | 218 | 11.11 | 11.47 | -0.36 | -3.11 | -0.045596866 | 1 | 1 | 0.25 |
| **turquoise** | Mid | 474 | 1742 | 1104 | 4417 | 42.93 | 39.44 | 3.5 | 8.87 | 0.122541001 | 0.006202476 | 0.006202476 | 0.25 |
| **turquoise** | Old | 536 | 1880 | 1104 | 4417 | 48.55 | 42.56 | 5.99 | 14.07 | 0.189898904 | 4.07E-06 | 6.11E-06 | 0.25 |
| **turquoise** | Young | 94 | 795 | 1104 | 4417 | 8.51 | 18 | -9.48 | -52.69 | -1.079895538 | 8.61E-24 | 2.58E-23 | 0.25 |
| **yellow** | Mid | 76 | 344 | 364 | 1456 | 20.88 | 23.63 | -2.75 | -11.63 | -0.178337241 | 0.175748091 | 0.175748091 | 0.25 |
| **yellow** | Old | 262 | 981 | 364 | 1456 | 71.98 | 67.38 | 4.6 | 6.83 | 0.095313675 | 0.033116463 | 0.099349389 | 0.25 |
| **yellow** | Young | 26 | 131 | 364 | 1456 | 7.14 | 9 | -1.85 | -20.61 | -0.332983283 | 0.169624486 | 0.175748091 | 0.25 |

**Table S6. Cross-species conservation of WGCNA co-expression modules**. The table reports module pairs showing significant ortholog overlap between species, integrated with eigengene trajectory similarity across shared developmental stages. Columns indicate: species A and B and their respective module colours; module sizes (number of genes); number of shared single-copy orthologs between modules; Jaccard index (intersection / union of ortholog sets); odds ratio, nominal p-value (one-sided Fisher's exact test) and Benjamini–Hochberg adjusted p-value for module overlap; number of shared developmental stages used for trajectory comparison; Pearson correlation coefficient between mean module eigengenes across shared stages and its nominal p-value; and logical flags indicating whether each pair showed significant ortholog overlap (padj < 0.05 and at least 50 shared orthologs) and correlated eigengene trajectories (|r| > 0.6 and p < 0.05). Module pairs meeting both criteria were considered conserved.

| **species_A** | **module_A** | **size_A** | **species_B** | **module_B** | **size_B** | **n_overlap** | **jaccard** | **odds_ratio** | **pval** | **padj** | **n_stages** | **rho** | **pval_L3** | **L1_overlap** | **Zsummary_min** | **L3_correlated** |
| --- | --- | --- | --- | --- | --- | --- | --- | --- | --- | --- | --- | --- | --- | --- | --- | --- |
| **Scy** | yellow | 518 | Und | blue | 943 | 171 | 0.13255814 | 1.690668875 | 3.05E-07 | 1.26E-05 | 5 | -0.984763272 | 0.00225256 | TRUE | 3.389930503 | TRUE |
| **Ddic** | red | 422 | Scy | yellow | 667 | 85 | 0.084661355 | 1.618904682 | 0.000206129 | 0.003710328 | 6 | 0.880926322 | 0.020423666 | TRUE | 2.623596794 | TRUE |
| **Ddic** | blue | 737 | Und | brown | 320 | 94 | 0.09761163 | 1.829539214 | 4.99E-06 | 9.42E-05 | 5 | 0.950391027 | 0.013164848 | TRUE | 2.064554652 | TRUE |
| **Ddic** | brown | 565 | Scy | blue | 897 | 130 | 0.097597598 | 1.327793237 | 0.00557772 | 0.047809031 | 6 | 0.965863513 | 0.00172806 | TRUE | 1.149865904 | TRUE |
| **Ddic** | red | 422 | Scy | blue | 897 | 110 | 0.090984285 | 1.581662167 | 9.61E-05 | 0.001921932 | 6 | -0.625798046 | 0.183841446 | TRUE | 5.226372788 | FALSE |
| **Scy** | red | 437 | Und | blue | 943 | 170 | 0.140495868 | 2.247292528 | 8.99E-14 | 4.94E-12 | 5 | 0.671647765 | 0.214389083 | TRUE | 4.825933402 | FALSE |
| **Ddic** | pink | 222 | Scy | turquoise | 831 | 54 | 0.054054054 | 1.547409672 | 0.005434897 | 0.047809031 | 6 | 0.071615019 | 0.892761118 | TRUE | 4.464385781 | FALSE |
| **Ddic** | blue | 950 | Scy | pink | 215 | 61 | 0.055253623 | 1.619870731 | 0.001709233 | 0.021975855 | 6 | 0.37073457 | 0.469375789 | TRUE | 4.420057092 | FALSE |
| **Ddic** | yellow | 424 | Scy | red | 518 | 93 | 0.109540636 | 2.571651541 | 3.79E-12 | 1.71E-10 | 6 | -0.085607477 | 0.871902478 | TRUE | 4.091313223 | FALSE |
| **Ddic** | greenyellow | 233 | Scy | red | 518 | 52 | 0.074391989 | 2.490677661 | 2.03E-07 | 6.08E-06 | 6 | -0.666854104 | 0.147991986 | TRUE | 4.091313223 | FALSE |
| **Scy** | turquoise | 777 | Und | turquoise | 1173 | 423 | 0.277013752 | 3.841304273 | 5.61E-59 | 9.26E-57 | 5 | 0.155511434 | 0.802797692 | TRUE | 3.386891715 | FALSE |
| **Ddic** | yellow | 353 | Und | blue | 924 | 166 | 0.149414941 | 3.138506057 | 1.28E-22 | 8.46E-21 | 5 | 0.56967848 | 0.316069888 | TRUE | 2.861767327 | FALSE |
| **Ddic** | greenyellow | 184 | Und | blue | 924 | 81 | 0.078870497 | 2.578610031 | 1.66E-09 | 4.37E-08 | 5 | -0.684561121 | 0.202305218 | TRUE | 2.861767327 | FALSE |
| **Ddic** | black | 197 | Und | turquoise | 1106 | 103 | 0.085833333 | 2.830781527 | 2.31E-12 | 1.02E-10 | 5 | -0.251593178 | 0.683073976 | TRUE | 1.845328135 | FALSE |
| **Ddic** | black | 230 | Scy | turquoise | 831 | 91 | 0.093814433 | 3.334165199 | 4.28E-16 | 2.57E-14 | 6 | 0.336613415 | 0.514150473 | TRUE | 1.419427488 | FALSE |
| **Scy** | green | 181 | Und | turquoise | 1173 | 83 | 0.065302911 | 2.072682011 | 2.14E-06 | 7.06E-05 | 5 | -0.603835273 | 0.28085745 | TRUE | 1.18415707 | FALSE |
| **Ddic** | brown | 449 | Und | yellow | 424 | 68 | 0.08447205 | 1.49732631 | 0.003803396 | 0.041837361 | 5 | 0.815731069 | 0.092284372 | TRUE | 0.497611366 | FALSE |
| **Ddic** | turquoise | 1081 | Scy | turquoise | 831 | 287 | 0.176615385 | 2.069418274 | 8.74E-18 | 7.87E-16 | 6 | 0.427691861 | 0.397578976 | TRUE | 0.282054245 | FALSE |
| **Ddic** | turquoise | 1081 | Scy | green | 209 | 68 | 0.055646481 | 1.674644177 | 0.000623039 | 0.009345581 | 6 | 0.731955518 | 0.098142558 | TRUE | 0.282054245 | FALSE |
| **Ddic** | turquoise | 1081 | Scy | black | 191 | 60 | 0.04950495 | 1.582470957 | 0.003247125 | 0.036989489 | 6 | -0.553420754 | 0.254618209 | TRUE | 0.282054245 | FALSE |
| **Ddic** | turquoise | 862 | Und | turquoise | 1106 | 370 | 0.231539424 | 2.241272745 | 4.21E-23 | 5.56E-21 | 5 | -0.856833733 | 0.063612357 | TRUE | -0.372065962 | FALSE |

**Table S7. RNA-seq samples used to genotype parental strains and generate the VCFs used for allele-specific expression analyses**. For each sample, the species, strain, sex, developmental stage, sequencing method (Bulk or Direct Low Input), number of PCR cycles, data source, and SRA/BioSample/BioProject accessions are indicated.

| **species** | **strain** | **sex** | **stage** | **Ectraction protocol** | **PCR cycles** | **Data source** | **SRR** | **BioSample** |
| --- | --- | --- | --- | --- | --- | --- | --- | --- |
| ***S. promiscuus*** | As6 | male | immature gametophyte | Bulk | NA | Lotharukpong et al. in prep. | RAP9_125_Scytosiphon_promiscuus_As6m_immGA_1 |  |
|  |  |  | immature gametophyte | Bulk | NA | Lotharukpong et al. in prep. | RAP9_126_Scytosiphon_promiscuus_As6m_immGA_2 |  |
|  |  |  | immature gametophyte | Bulk | NA | Lotharukpong et al. in prep. | RAP9_127_Scytosiphon_promiscuus_As6m_immGA_3 |  |
|  |  |  | immature parthenosporophyte | Bulk | NA | Lotharukpong et al. in prep. | RAP34_592_Scytosiphon_promiscuus_As6m_immPSP_1 |  |
|  |  |  | immature parthenosporophyte | Bulk | NA | Lotharukpong et al. in prep. | RAP34_593_Scytosiphon_promiscuus_As6m_immPSP_2 |  |
|  |  |  | immature parthenosporophyte | Bulk | NA | Lotharukpong et al. in prep. | RAP34_594_Scytosiphon_promiscuus_As6m_immPSP_3 |  |
|  |  |  | mature gametophyte | Bulk | NA | Lotharukpong et al. in prep. | Scytosiphon lomentaria_Asbm_male_matGA_1 |  |
|  |  |  | mature gametophyte | Bulk | NA | Lotharukpong et al. in prep. | Scytosiphon lomentaria_Asbm_male_matGA_2 |  |
|  |  |  | mature gametophyte | Bulk | NA | Lotharukpong et al. in prep. | Scytosiphon lomentaria_Asbm_male_matGA_3 |  |
|  |  |  | male gamete | Direct Low input | 15 | Lotharukpong et al. in prep. | RAP12_179_Scytosiphon_promiscuus_sxs103f_gametes_DirectLowIN_4_S16 |  |
|  |  |  | male gamete | Direct Low input | 15 | Lotharukpong et al. in prep. | RAP12_178_Scytosiphon_promiscuus_sxs103f_gametes_DirectLowIN_3_S15 |  |
|  |  |  | male gamete | Direct Low input | 15 | Lotharukpong et al. in prep. | RAP12_177_Scytosiphon_promiscuus_sxs103f_gametes_DirectLowIN_2_S14 |  |
|  |  |  | male gamete | Bulk | NA | Lotharukpong et al. in prep. | Scytosiphon lomentaria_Asbm_male_Gametes_1 |  |
|  |  |  | male gamete | Bulk | NA | Lotharukpong et al. in prep. | Scytosiphon lomentaria_Asbm_male_Gametes_2 |  |
|  |  |  | male gamete | Bulk | NA | Lotharukpong et al. in prep. | Scytosiphon lomentaria_Asbm_male_Gametes_3 |  |
|  | As9 | female | female gamete | Direct Low input | 20 | this study | SRR37846298 | SAMN56777093 |
|  |  |  | female gamete | Direct Low input | 20 | this study | SRR37846297 | SAMN56777093 |
|  |  |  | female gamete | Direct Low input | 20 | this study | SRR37846251 | SAMN56777093 |
|  | Mr5 (sxs103) | female | immature gametophyte | Bulk | NA | Lotharukpong et al. in prep. | RAP9_112_Scytosiphon_promiscuus_sxs103f_immGA_1 |  |
|  |  |  | immature gametophyte | Bulk | NA | Lotharukpong et al. in prep. | RAP9_113_Scytosiphon_promiscuus_sxs103f_immGA_2 |  |
|  |  |  | immature gametophyte | Bulk | NA | Lotharukpong et al. in prep. | RAP9_114_Scytosiphon_promiscuus_sxs103f_immGA_3 |  |
|  |  |  | mature gametophyte | Bulk | NA | Lotharukpong et al. in prep. | RAP7_72_Scytosiphon_promiscuus_sxs103f_matGA_1 |  |
|  |  |  | mature gametophyte | Bulk | NA | Lotharukpong et al. in prep. | RAP7_73_Scytosiphon_promiscuus_sxs103f_matGA_2 |  |
|  |  |  | mature gametophyte | Bulk | NA | Lotharukpong et al. in prep. | RAP7_74_Scytosiphon_promiscuus_sxs103f_matGA_3 |  |
|  |  |  | immature parthenosporophyte | DirectLowInput | 20 | Lotharukpong et al. in prep. | RAP17_287_Scytosiphon_promiscuus_Mr5f_earlyPSP5cells_Dlowin_1 |  |
|  |  |  | immature parthenosporophyte | DirectLowInput | 20 | Lotharukpong et al. in prep. | RAP17_288_Scytosiphon_promiscuus_Mr5f_earlyPSP5cells_Dlowin_2 |  |
|  |  |  | immature parthenosporophyte | DirectLowInput | 20 | Lotharukpong et al. in prep. | RAP17_289_Scytosiphon_promiscuus_Mr5f_earlyPSP5cells_Dlowin_3 |  |
|  |  |  | immature parthenosporophyte | DirectLowInput | 20 | Lotharukpong et al. in prep. | RAP17_290_Scytosiphon_promiscuus_Mr5f_earlyPSP5cells_Dlowin_4 |  |
|  |  |  | female gamete | Direct Low input | 15 | Lotharukpong et al. in prep. | RAP12_175_Scytosiphon_promiscuus_As6m_gametes_DirectLowIN_4_S13 |  |
|  |  |  | female gamete | Direct Low input | 15 | Lotharukpong et al. in prep. | RAP12_174_Scytosiphon_promiscuus_As6m_gametes_DirectLowIN_3_S12 |  |
|  |  |  | female gamete | Direct Low input | 15 | Lotharukpong et al. in prep. | RAP12_172_Scytosiphon_promiscuus_As6m_gametes_DirectLowIN_1_S11 |  |
|  | Mr8 | male | male gamete | Direct Low input | 21 | this study | SRR37846215 | SAMN56777104 |
|  |  |  | male gamete | Direct Low input | 21 | this study | SRR37846204 | SAMN56777104 |
|  |  |  | male gamete | Direct Low input | 21 | this study | SRR37846193 | SAMN56777104 |
| ***U. pinnatifida*** | Un1 | female | mature gametophyte | Bulk | NA | this study | SRR37846281 | SAMN56777110 |
|  |  |  | mature gametophyte | Bulk | NA | this study | SRR37846280 | SAMN56777110 |
|  |  |  | mature gametophyte | Bulk | NA | this study | SRR37846279 | SAMN56777110 |
|  |  |  | immature gametophyte | Bulk | NA | this study | SRR37846278 | SAMN56777107 |
|  |  |  | immature gametophyte | Bulk | NA | this study | SRR37846277 | SAMN56777107 |
|  |  |  | immature gametophyte | Bulk | NA | this study | SRR37846276 | SAMN56777107 |
|  |  |  | gametophyte | Bulk | NA | Vigneau et al. 2026 | SRR35498842 | SAMN51363669 |
|  |  |  | gametophyte | Bulk | NA | Vigneau et al. 2026 | SRR35498841 | SAMN51363669 |
|  |  |  | gametophyte | Bulk | NA | Vigneau et al. 2026 | SRR35498841 | SAMN51363669 |
|  |  | male | matGA | Bulk | NA | this study | SRR37846275 | SAMN56777109 |
|  |  |  | matGA | Bulk | NA | this study | SRR37846273 | SAMN56777109 |
|  |  |  | matGA | Bulk | NA | this study | SRR37846272 | SAMN56777109 |
|  |  |  | immGA | Bulk | NA | this study | SRR37846271 | SAMN56777106 |
|  |  |  | immGA | Bulk | NA | this study | SRR37846270 | SAMN56777106 |
|  |  |  | immGA | Bulk | NA | this study | SRR37846269 | SAMN56777106 |
|  |  |  | gametophyte | Bulk | NA | Vigneau et al. 2026 | SRR35498839 | SAMN51363670 |
|  |  |  | gametophyte | Bulk | NA | Vigneau et al. 2026 | SRR35498837 | SAMN51363670 |
|  |  |  | gametophyte | Bulk | NA | Vigneau et al. 2026 | SRR35498836 | SAMN51363670 |
|  | UnpinK2 | female | gametophyte | Bulk | NA | this study | SRR37846268 | SAMN56777115 |
|  |  |  | gametophyte | Bulk | NA | this study | SRR37846267 | SAMN56777115 |
|  |  | male | gametophyte | Bulk | NA | this study | SRR37846266 | SAMN56777116 |
|  |  |  | gametophyte | Bulk | NA | this study | SRR37846265 | SAMN56777116 |
| ***D. dichotoma*** | ODC1387 | male | immature gametophyte | Bulk | NA | Lotharukpong et al. in prep. | RAP_3_1_Dyctiota_dichotoma_ODC1387male_immGA_1 |  |
|  |  |  | immature gametophyte | Bulk | NA | Lotharukpong et al. in prep. | RAP_3_2_Dyctiota_dichotoma_ODC1387male_immGA_2 |  |
|  |  |  | immature gametophyte | Bulk | NA | Lotharukpong et al. in prep. | RAP_3_3_Dyctiota_dichotoma_ODC1387male_immGA_3 |  |
|  |  |  | mature gametophyte | Bulk | NA | Lotharukpong et al. in prep. | RAP_4_4_Dyctiota_dichotoma_ODC1387male_matGA_1 |  |
|  |  |  | mature gametophyte | Bulk | NA | Lotharukpong et al. in prep. | RAP_4_5_Dyctiota_dichotoma_ODC1387male_matGA_2 |  |
|  |  |  | mature gametophyte | Bulk | NA | Lotharukpong et al. in prep. | RAP_4_6_Dyctiota_dichotoma_ODC1387male_matGA_3 |  |
|  | ODC2377 | female | gametophyte | Bulk | NA | this study | SRR37846264 | SAMN56777087 |
|  |  |  | gametophyte | Bulk | NA | this study | SRR37846262 | SAMN56777087 |
|  |  |  | female gamete | Direct Low input | 21 | this study | SRR37846174 | SAMN56777075 |
|  |  |  | female gamete | Direct Low input | 21 | this study | SRR37846173 | SAMN56777075 |
|  |  |  | female gamete | Direct Low input | 21 | this study | SRR37846172 | SAMN56777075 |
|  | KB07m | male | gametophyte | Bulk | NA | this study | SRR37846261 | SAMN56777077 |
|  |  |  | gametophyte | Bulk | NA | this study | SRR37846260 | SAMN56777077 |
|  | KB07f | female | gametophyte | Bulk | NA | this study | SRR37846259 | SAMN56777076 |
|  |  |  | gametophyte | Bulk | NA | this study | SRR37846258 | SAMN56777076 |
|  |  |  | immature gametophyte | Bulk | NA | Lotharukpong et al. in prep. | RAP11_166_Dictyota_dichotoma_KB07f_immGA_1_S10 |  |
|  |  |  | immature gametophyte | Bulk | NA | Lotharukpong et al. in prep. | RAP11_167_Dictyota_dichotoma_KB07f_immGA_2_S11 |  |
|  |  |  | immature gametophyte | Bulk | NA | Lotharukpong et al. in prep. | RAP11_168_Dictyota_dichotoma_KB07f_immGA_3_S12 |  |
|  |  |  | female gamete | Bulk | NA | Lotharukpong et al. in prep. | RAP30_548_Dictyota_dichotoma_KB07f_gametes_3_S18 |  |
|  |  |  | female gamete | Bulk | NA | Lotharukpong et al. in prep. | RAP30_547_Dictyota_dichotoma_KB07f_gametes_2_S17 |  |
|  |  |  | female gamete | Bulk | NA | Lotharukpong et al. in prep. | RAP30_546_Dictyota_dichotoma_KB07f_gametes_1_S16 |  |
|  |  |  | female gamete | Direct Low input | 18 | Lotharukpong et al. in prep. | RAP15_245_Dictyota_dichotoma_KB07f_gametes_DLowIN_1 |  |
|  |  |  | female gamete | Direct Low input | 18 | Lotharukpong et al. in prep. | RAP15_246_Dictyota_dichotoma_KB07f_gametes_DLowIN_2 |  |
|  |  |  | female gamete | Direct Low input | 18 | Lotharukpong et al. in prep. | RAP15_247_Dictyota_dichotoma_KB07f_gametes_DLowIN_3 |  |

**Table S8. VCF statistics.** For each VCF, the number of SNPs, the number of genes containing at least one SNP, the percentage of genes containing at least one SNP, the mean and median number of SNP per gene are reported.

| **species** | **cross** | **number of SNPs differentiating the strains** | **number of genes containing at least one SNP** | **percentage of genes containing at least one SNP** | **mean number of SNP per gene** | **median number of SNP per gene** |
| --- | --- | --- | --- | --- | --- | --- |
| ***Scytosiphon promiscuus*** | Mr5 x As6 | 773,941 | 16,141 | 84% | 32 | 15 |
| ***Scytosiphon promiscuus*** | As9 x Mr8 | 90,976 | 5,700 | 29.60% | 3.5 | 0 |
| ***Undaria pinnatifida*** | Un1 x UpinK2 | 373,406 | 9,148 | 57.40% | 11.9 | 1 |
| ***Dictyota dichotoma*** | KB07f x ODC1387 | 113,592 | 8,972 | 43.60% | 3.6 | 0 |
| ***Dictyota dichotoma*** | ODC2377 x KB07f | 323,323 | 13,481 | 65.50% | 9.2 | 4 |
| ***Dictyota dichotoma*** | KB07f x KB07m | 61,049 | 3,302 | 16% | 1.4 | 0 |

Data S1. (separate file)
Full GO-term datasets obtained with InterProScan expanded to include ancestral terms on the full proteome of *Dictyota dichotoma*, *Undaria pinnatifida*, and *Scytosiphon promiscuus* respectively.

**Data S2. (separate file)**
GO-term dataset obtained by mapping the GO terms from the full GO-term datasets on the generic GO-slim database.

**Data S3. (separate file)**
Cross specific vcf files obtained by filtering of parental-strain-specific SNPs in each studied cross (see Methods).

**Data S4. (separate file)**

Aggregation of various analysis results. DESeq2 analysis for each stage vs egg cell for oogamous species or gametes for Scytosiphon promiscuus; columns report log₂ fold change (log2FoldChange), standard error (lfcSE), and adjusted p-value (padj) relative to female gametes for each developmental stage (zygote, polarisation, 2-celled, 4-celled, patterning), together with a binary significance flag (padj < 0.05 and |log₂FC| ≥ 1) (sig). WGCNA co-expression module assignment (module), module membership score (MM) and its p-value (MM_pval), and gene significance (GS) and its p-value (GS_pval). Parental allele contribution per stage for each cross, with mean TPM across replicates (meanTPM), mean maternal fraction (mean_maternal_fraction), log₂ fold change (logFC), log₂ counts per million (logCPM), p-value (PValue), and FDR from edgeR's exact test comparing maternally and paternally assigned read counts normalised by total number of sorted reads; and a genome-of-origin bias category (category): maternally biased (FDR < 0.05 and logFC < −1), paternally biased (FDR < 0.05 and logFC > 1), unbiased (FDR ≥ 0.05 or |logFC| ≤ 1), not expressed (filtered by filterByExpr and mean TPM < 5), or undetermined (no test result but mean TPM ≥ 5).

**Data S5. (separate file)**

Single-copy orthologous genes acrossorthogroups used for cross-species transcriptome comparison. Each row represents an orthogroup containing exactly one gene from each of the three brown algal species. Columns indicate the orthogroup identifier and the corresponding gene identifiers for *Dictyota dichotoma*, *Scytosiphon promiscuus*, and *Undaria pinnatifida*.

**Data S6. (separate file)**

Average PC3 loadings of single-copy orthogroups from cross-species PCA. Orthogroups ranked by their loading on PC3 of the cross-species PCA on log2(TPM + 1)-transformed orthogroup expression. PC3 captures a conserved developmental trajectory shared across the three species, with sample scores along this axis ordering consistently with developmental progression in each
